# Complete elucidation and heterologous reconstruction of the biosynthetic pathway of camptothecin

**DOI:** 10.64898/2026.06.23.733941

**Authors:** Tong Zhang, Ying Xiong, Kuangqin Chen, Shiwen Wu, Xing Yan, Junsu Zhou, Yan Wang, Chengshuai Yang, Pingping Wang, Zhihua Zhou

## Abstract

Camptothecin derivatives are first-line anticancer drugs used worldwide for the treatment of diverse malignant tumors. However, the biosynthetic pathway of camptothecin has remained elusive for five decades. Here, we fully map its entire biosynthetic route. We discovered five key missing enzymes (OpCAR, OpSDR11, OpCS, OpGH1, and OpSTR) *via* the combination of MALDI mass spectrometry imaging, single-cell RNA sequencing and co-expression analysis. Meanwhile, we demonstrated a free flavin mononucleotide triggered the non-enzymatic 6-5-6 to 6-6-5 fused-ring skeleton rearrangement, filling the last gap in camptothecin biosynthesis. Finally, we validated this identified pathway and achieved the *de novo* biosynthesis of camptothecin in *Saccharomyces cerevisiae*. These discoveries uncover the long-standing mystery underlying camptothecin and pave the way for manufacturing camptothecin and its derivatives through synthetic biology approaches.

## Introduction

Camptothecin **9** and its derivatives, exhibiting the potent inhibitory activity against DNA topoisomerase I and represent the second-largest class of anticancer drugs worldwide (*1*). Since 1996, two derivatives topotecan and irinotecan have been approved by US Food and Drug Administration for the treatment of malignant tumors, such as small cell lung cancer, colorectal cancer, recurrent ovarian cancer and cervical cancer, generating annual sales of approximately one billion dollars worldwide (*2*). Camptothecin **9**, together with the famous antimitotic drug vinblastine, antihypertensive drug ajmalicine and antimalarial drug quinine, belongs to monoterpene indole alkaloids (MIAs) class. MIAs comprise over 3,000 structurally and pharmacologically diverse compounds and are regarded as one of the most diverse groups of plant secondary metabolites (*3, 4*). Several taxonomically diverse plant species are capable of synthesizing camptothecin **9**, among which, *Camptotheca acuminata*, *Ophiorrhiza pumila*, and *Nothapodytes nimmoniana* exhibit higher camptothecin **9** accumulations than others (*5*).

Total chemical synthesis of camptothecin **9** requires harsh reaction condition and incurs high cost, hindering its industrial application (*6–8*). At present, camptothecin **9** is mainly extracted from *C. acuminata* and *N. nimmoniana*, and subsequently converted into various derivatives through chemical methods (*9–11*). However, its content is very low (0.012-0.236% dry weight) in the woody plants *C. acuminata* and *N. nimmoniana* (*12–14*), and producing one ton of high-purity camptothecin **9** consumes 1,000 to 1,500 tons of dry plant raw materials (*15, 16*). In 2024, irinotecan and topotecan collectively accounted for 85%–90% of the total consumption of pharmaceutical-grade camptothecin **9**, corresponding to roughly 12 tons of camptothecin **9** equivalent (*2, 17*). This indicates a massive annual consumption of plant resources. Therefore, achieving sustainable production of camptothecin **9** is urgently required. Heterologous biosynthesis of camptothecin **9** using microbial fermentation represents a promising approach to achieve a sustainable supply, yet its implementation depends on a thorough understanding of the biosynthetic pathway (*18–21*).

Camptothecin **9** is biosynthesized *via* a branch pathway of the MIAs metabolism, starting from the canonical MIAs precursor strictosidine **10** in *O. pumila* (*22, 23*). The upstream metabolic route from primary metabolites to strictosidine **10** has been fully elucidated (table S1) (*24–26*). Strictosidine **10** is generated by the Pictet-Spenglerase strictosidine synthase (STR) through intermolecular cyclization of tryptamine **1** and the monoterpene glucoside secologanin **2** (*27*). Nevertheless, the downstream biosynthetic steps from strictosidine **10** to camptothecin **9** have remained unclear for over 50 years (*28, 29*). In particular, the mechanisms underlying formation of lactam ring skeleton, the 6-5-6 to 6-6-5 fused-ring rearrangement and the subsequent tailoring modifications remain elusive (Fig. 1). Previous isotope-labeling experiments and chemical deduction have implied a putative biosynthetic cascade involving strictosamide **3**, strictosamide epoxide **29**, strictosamide diol **30**, strictosamide ketolactam **31**, pumiloside **11**, and (3*S*)-deoxypumiloside **12** prior to the formation of the camptothecin skeleton (fig. S1), however, the enzymes involved have not yet been identified (*1, 30*).

**Fig. 1.**
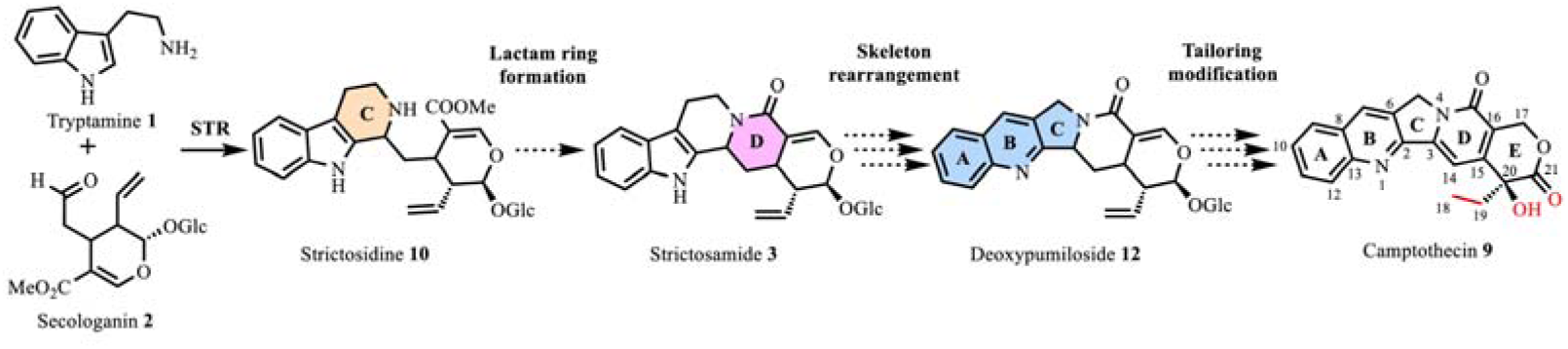
The proposed biosynthetic route of camptothecin 9 in *O. pumila*. Black solid arrow denotes previously characterized enzymatic reaction, and black dashed arrows denote proposed reactions catalyzed by enzymes that remained unclear. The upstream biosynthetic pathway of tryptamine **1** and secologanin **2** as well as strictosidine **10** have been elucidated, whereas these enzymes responsible for lactam formation, skeleton rearrangement, and skeleton tailoring modification at late-stage to produce camptothecin **9** were still unknown.

Here we report the complete elucidation of the camptothecin **9** biosynthetic pathway in *O. pumila* through Matrix-Assisted Laser Desorption Ionization-Mass Spectrometry Imaging (MALDI-MSI), single-cell RNA sequencing and co-expression analysis. Moreover, reconstitution of this identified complete pathway in *S. cerevisiae* achieved *de novo* camptothecin **9** production (0.55 ug/L), marking a milestone in the biosynthesis of the anticancer drug camptothecin **9** and its derivatives.

## Results

### Discovery of the key candidate *OpCAR*

*O. pumila* naturally accumulates camptothecin **9** at a relatively high content and is a Rubiaceae plant with the smallest known genome among the camptothecin-producing species (*1, 28*). To analyze the distribution of camptothecin **9** in this plant, we measured its content in the root, leaf and stem tissues of *O. pumila* and found that the content in the root tissue was higher than that in the leaf and stem tissues (Fig. 2A). However, RNA-sequencing analysis of these three tissues revealed that the expression pattern of four known genes involved in the early camptothecin **9** biosynthetic pathway (*OpTDC*, *OpSLS*, *OpLAMT*, and *OpSTR*) were not correlated to the distribution pattern of camptothecin **9** across the three tissues (figs. S2 to S4). Thus, co-expression analysis based solely on these four upstream-pathway genes was insufficient to identify candidate genes for the unknown downstream steps. We therefore performed MALDI-MSI of *O. pumila* root and found that camptothecin **9** and other known intermediates strictosidine **10**, strictosamide **3**, (3*S*)-deoxypumiloside **12**, and deoxycamptothecin **8** mainly accumulated in the root cortex (Fig. 2, B and C). These results imply that some reactions involved in the biosynthesis of camptothecin **9** may occur predominantly in cortex cells, raising the possibility that specifically expressed genes in this tissue could serve as candidates. Based on this hypothesis, we performed single-cell RNA sequencing (scRNA-seq) using the *O. pumila* hairy root (Fig. 2D, figs. S5 to S11 and table S3). Expression quantification analysis of the known genes in the camptothecin **9** biosynthetic route revealed that *OpSLS*, *OpTDC* and *OpSTR* were specifically and highly expressed in the cortex-3 (Fig. 2E).

**Fig. 2.**
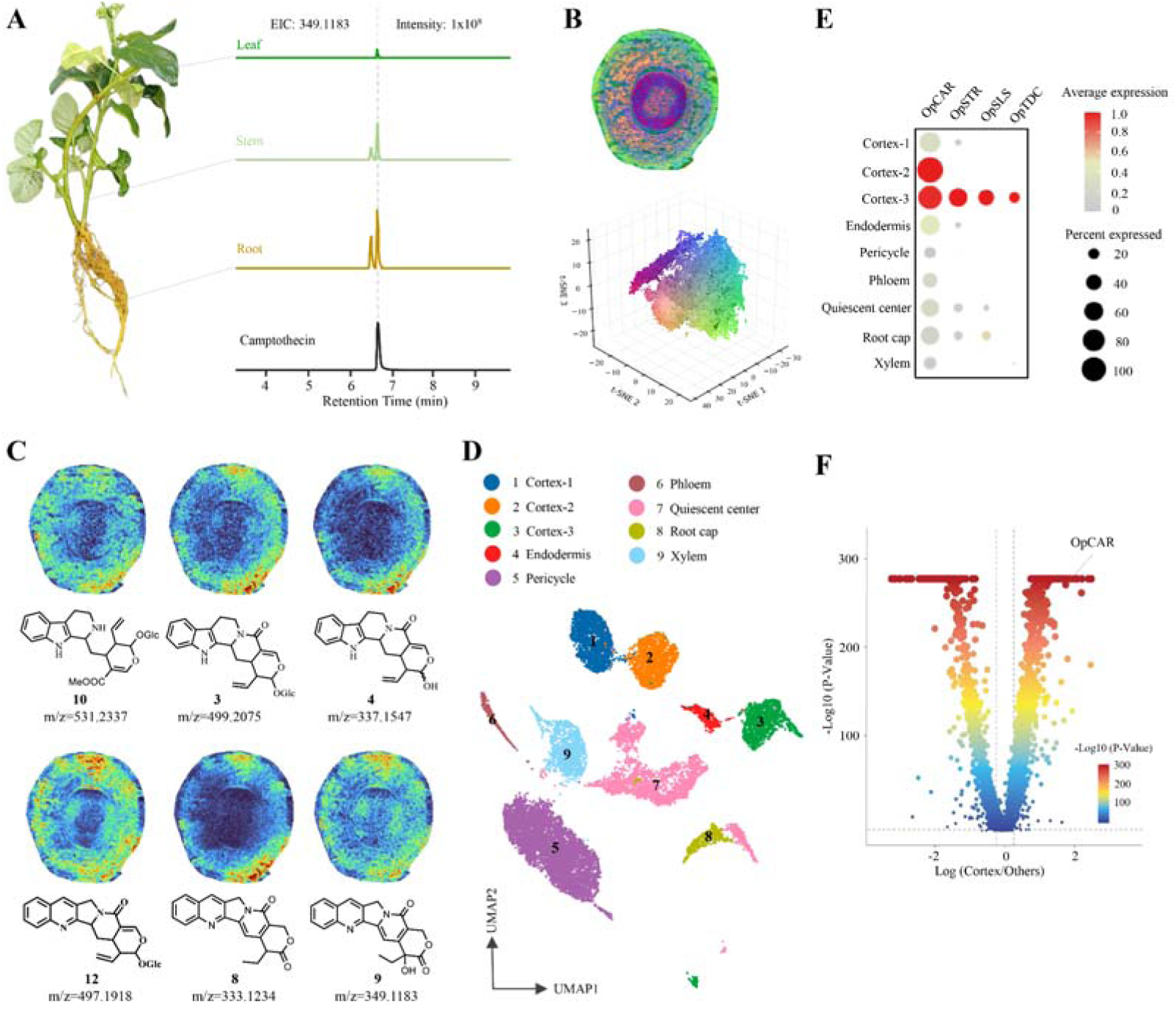
Spatial metabolomics and single-cell transcriptomics to discover candidate genes for the camptothecin biosynthesis. **(A)** LC-MS quantification analysis of camptothecin **9** in root, stem, and leaf tissues of *O. pumila*. **(B)** Cross-section of an *O. pumila* root showing the spatial distribution of metabolites. Colors indicate compounds originating from distinct root regions. **(C)** Mass spectrometry imaging of the spatial distribution of strictosidine **10**, strictosamide **3**, strictosamide aglycone **4**, deoxypumiloside **12**, deoxycamptothecin **8**, and camptothecin **9** in *O. pumila* root cross-sections. **(D)** UMAP visualization of seven cell types identified by scRNA-seq. Colors represent distinct cell types; cortex cells are subdivided into three subgroups. **(E)** Dot plot of expression levels of *OpCAR*, *OpSTR*, *OpTDC*, and *OpSLS* across cell types. Color intensity indicates relative expression level (red, high; gray, low); dot size represents the fraction of cells expressing each gene. **(F)** Differential gene expression analysis comparing cortex cells with all other cell types, highlighting genes specifically enriched in cortex cells.

Given these findings, genes highly and specifically expressed in the cortex cells were considered candidates involved in camptothecin **9** biosynthesis. Differential gene expression analysis between cortex cells and other cells led to the discovery of *OpCAR*, the sole gene encoding a reductase that potentially catalyzes the reduction of the C18-C19 double bond to form the C20-ethyl of camptothecin **9** (Fig. 2F and table S4).

### Three key enzymes for the tailoring modification of camptothecoside aglycone to camptothecin

For functional characterization, we expressed the candidate gene *OpCAR* in *Escherichia coli* and then performed *in vitro* enzymatic reaction by incubating the OpCAR crude proteins and camptothecoside aglycone **6** as a substrate. A corresponding ion peak for reduction product, compound **7** (*m/z* [M+H]^+^ = 335.1390), could be observed in liquid chromatography mass spectrum (LC-MS) analysis (Fig. 3, A and B). Through MS/MS analysis (Fig. 3B), the produced compound **7** by OpCAR was proposed to be the double bond (C18, C19) reduction product of camptothecoside aglycone **6**.

**Fig. 3.**
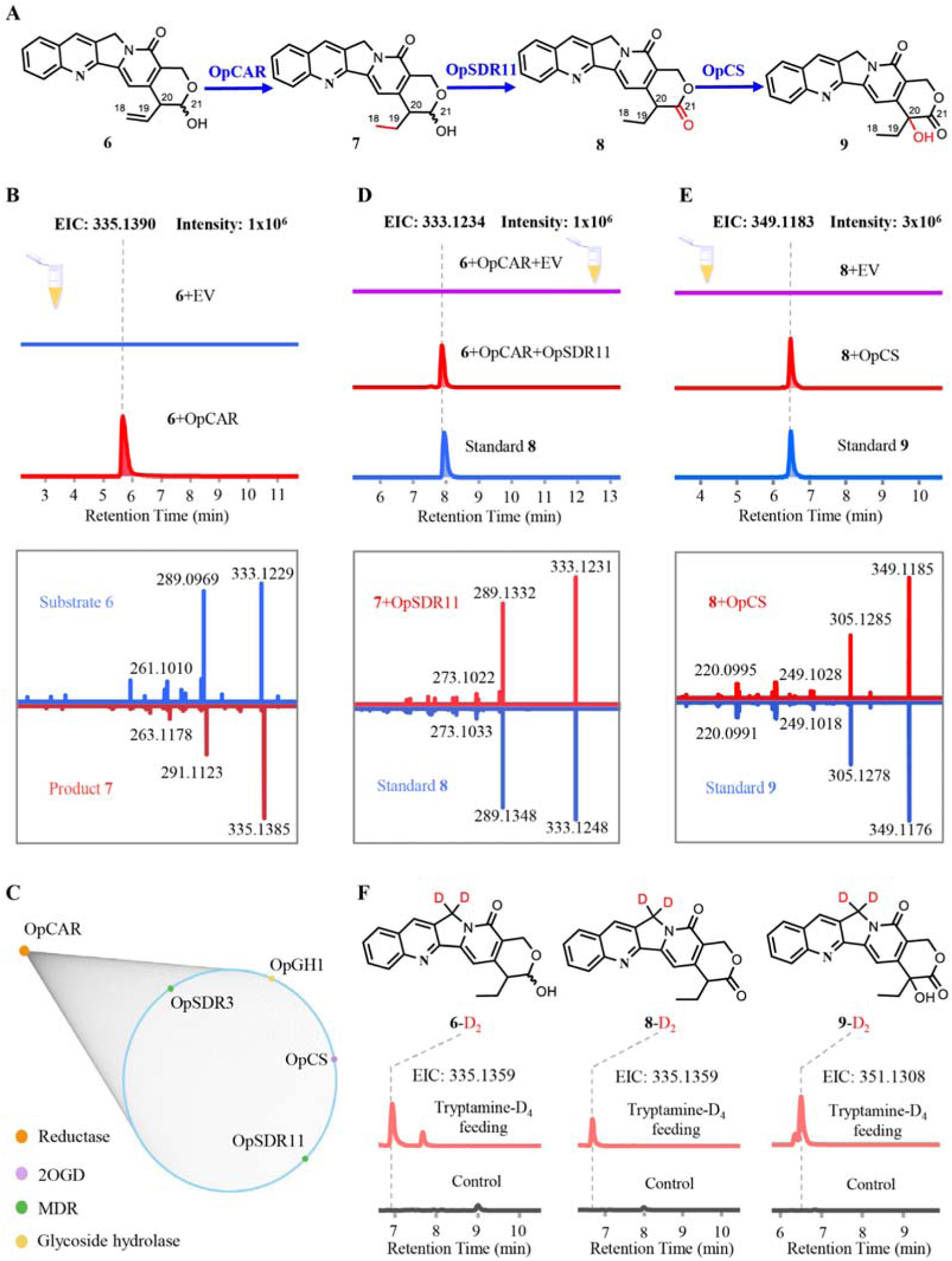
Biochemical characterization of tailoring enzymes for camptothecin 9 biosynthesis. **(A)** The biosynthetic route from camptothecoside aglycone **6** to camptothecin **9**. **(B)** LC-MS analysis of compound **7** generated by OpCAR using camptothecoside aglycone **6** as the substrate. EIC for compound **7** (m/z [M+H]^+^ = 335.1390) are shown. MS/MS spectrum of **7** produced by purified OpCAR; diagnostic fragment ions are compared with those of authentic standard. **(C)** Co-expression network of the *O. pumila* transcriptome using *OpCAR* as a bait. Each point represents one of 300 genes with a Pearson correlation coefficient (PCC) > 0.85 relative to *OpCAR*. **(D)** LC-MS analysis of compound **8** produced by OpSDR11 and OpCAR. Crude compound **7** generated by OpCAR was incubated with OpSDR11 expressed in *E. coli*. EIC for compound **8** (m/z [M+H]^+^ = 333.1234) are shown. MS/MS spectrum of compound **8** produced by purified OpSDR11; diagnostic fragment ions are compared with those of authentic deoxycamptothecin **8**. **(E)** LC-MS analysis of compound **9** produced by OpCS expressed in *E. coli*. EIC for compound **9** (m/z [M+H]^+^ = 349.1183) are shown. MS/MS spectrum of compound **9** produced by purified OpCS; diagnostic fragment ions are compared with those of authentic camptothecin **9**. **(F)** Detection of key intermediates by isotope labeling. *O. pumila* hairy roots were cultured in medium supplemented with tryptamine-D4. Deuterium-labeled compounds were identified by comparison with corresponding unlabeled authentic standards by LC-MS analysis. Ethanol, the solvent of tryptamine-D_4_, was added into the *O. pumila* hairy roots as the negative control. EV, empty vector.

Starting from compound **7**, the dehydrogenation at C21 and oxidation at C20 were required to generate camptothecin **9**. According to previous studies, short-chain dehydrogenase/reductase (SDR) have been proposed to be involved in this dehydrogenation reaction and cytochrome P450 (P450) or 2-oxoglutarate-dependent dioxygenase (2OGD) enzymes were supposed to catalyze this hydroxylation reaction (Fig. 3A) (*31–36*). Given that *OpCAR* was demonstrated to be involved in the camptothecin **9** biosynthesis and was highly expressed in the *O. pumila* root, we performed co-expression analysis across the *O. pumila* tissue transcriptomes using *OpCAR* as bait. Genes encoding SDR, P450, and 2OGD that exhibited co-expression patterns with *OpCAR* were selected as candidate genes for functional characterization (Fig. 3C).

To identify the SDR responsible for the C21 dehydrogenation of compound **7** to form deoxycamptothecin **8**, six *SDR* candidate genes were cloned and functionally characterized *via in vitro* reactions by incubating these expressed and purified SDR proteins and compound **7** (obtained from the reaction of OpCAR with camptothecoside aglycone **6**) as a substrate, respectively. LC-MS analysis revealed the extracted ion chromatograms (EIC) peak (*m/z* [M+H]^+^ = 333.1234) corresponding to a newly produced compound was detected only in OpSDR3 and OpSDR11 reactions, and its retention time and fragments matched those of the standard deoxycamptothecin **8**, indicating OpSDR3 and OpSDR11 are responsible for C21 dehydrogenation to produce the target deoxycamptothecin **8** (Fig. 3D). The yield of deoxycamptothecin **8** in the OpSDR11 reaction was higher than that in the OpSDR3 reaction, indicating the higher catalytic activity of OpSDR11 (fig. S12). Moreover, we also found that OpSDR11 could catalyze the C21 dehydrogenation of camptothecoside aglycone **6** to form compound **15**, which could then be converted into deoxycamptothecin **8** by OpCAR, indicating that two parallel routes from camptothecoside aglycone **6** to deoxycamptothecin **8** exist (fig. S13).

The conversion of deoxycamptothecin **8** to camptothecin **9**, hydroxylation at the C20 position is required. We first screened twelve P450 genes highly expressed in *O. pumila* root *via* expressing each of them in *S. cerevisiae* and obtained microsomes for *in vitro* reaction using deoxycamptothecin **8** as a substrate. However, none of these proteins could catalyze deoxycamptothecin **8** to produce camptothecin **9**. We therefore turned our attention to 2OGDs, and four 2OGD candidate genes were functionally characterized *via in vitro* reaction using the crude enzyme of each 2OGD candidate gene expressed in *E. coli*. LC-MS analysis indicated a newly produced compound, sharing the same retention time and MS fragments as standard camptothecin **9**, was detected in the OpOGD1 reaction (Fig. 3E). To further confirm the chemical structure of this newly produced compound, we performed a large-scale reaction to convert deoxycamptothecin **8** and obtained purified product for NMR analysis. ^1^H NMR and ^13^C NMR analyses demonstrated that the reaction product of OpOGD1 was camptothecin **9** (figs. S34 and S35) and OpOGD1 was accordingly renamed as OpCS. Collectively, the discovery and identification of the three tailoring enzymes OpSDR, OpCAR and OpCS uncovered the biosynthetic route from camptothecoside aglycone **6** to camptothecin **9**.

To determine whether above key intermediates exist in *O. pumila* plant, we performed an isotope feeding experiment by adding tryptamine-D_4_ into *O. pumila* hairy roots cultures. Isotopically labeled camptothecin **9** and two key intermediates, camptothecoside aglycone **6** and deoxycamptothecin **8**, could be detected in the *O. pumila* hairy roots (Fig. 3F and figs. S25 to S31). Compounds **7** and **15** were not detected, possibly due to their structural instability or rapid conversion to downstream products.

### OpGH1 and FMN trigger skeleton rearrangement to generate camptothecoside aglycone

After elucidating the skeleton tailoring modification route from camptothecoside aglycone **6** to camptothecin **9**, we then attempted to unearth the skeleton rearrangement process linking strictosamide **3** to camptothecoside aglycone **6**. Conversion of strictosamide **3** to camptothecoside aglycone **6** requires a glycoside hydrolase to cleave the glucosyl moiety (Fig. 4A). To obtain the responsible enzyme, five glycoside hydrolase candidates exhibiting co-expression pattern with OpCAR were cloned and functionally characterized *via in vitro* enzymatic reactions using each of their crude enzymes expressed in *E. coli* and strictosamide **3** as a substrate. Among them, OpGH1 efficiently consumed strictosamide **3** (fig. S14, A and B), and a newly produced compound, showing loss of a glucose group (*m/z* [M+H]^+^ = 337.1547) was detected by LC-MS analysis (Fig. 4B). MS/MS analysis results revealed the fragments of the newly produced compound-337, 319, 267, 237, and 171-were identical with the substrate strictosamide **3**, and that the molecular weight of the newly produced compound was 162 less than that of strictosamide **3** (fig. S14, C and D), indicating OpGH1 hydrolyzes the glucose group of strictosamide **3** to produce strictosamide aglycone **4**.

**Fig. 4.**
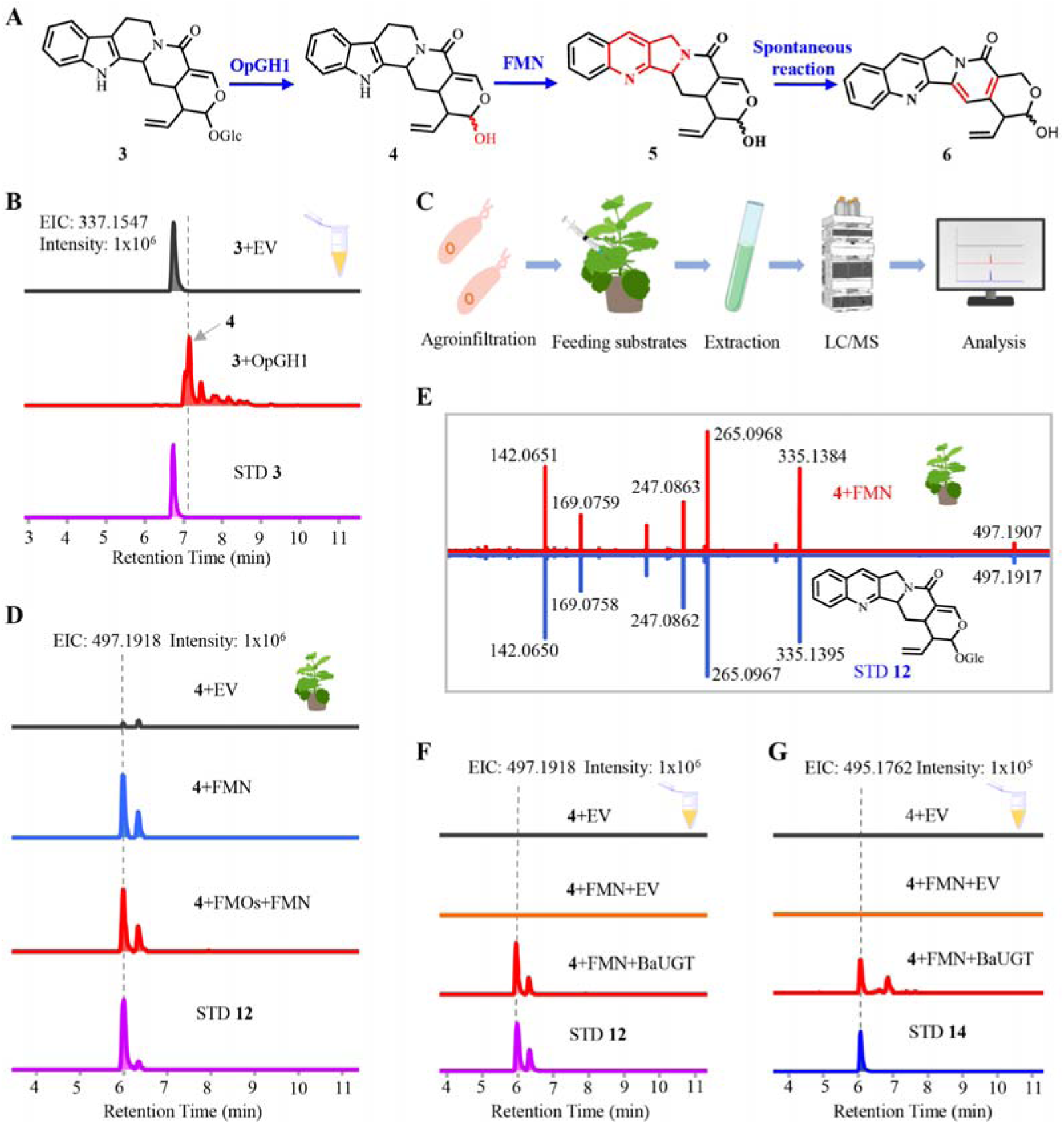
Biochemical characterization of OpGH1 and FMN responsible for the formation of camptothecin 9 skeleton. **(A)** Biosynthetic route to camptothecoside aglycone **6** from strictosamide **3**. **(B)** LC-MS analysis of the product of *in vitro* reaction of OpGH1 using strictosamide aglycone **4** as substrate. EIC for strictosamide aglycone **4** (m/z [M+H]^+^ = 337.1547) are shown. **(C)** Schematic of the *N. benthamiana* transient expression system. *Agrobacterium tumefaciens* strains harboring target genes were infiltrated into *N. benthamiana* leaves, followed by substrate feeding and LC-MS analysis. **(D)** EIC for (3*S*)-deoxypumiloside **12** (m/z [M+H]^+^ = 497.1918) from *N. benthamiana* expressing the indicated genes and fed with FMN and substrate strictosamide aglycone **4**. **(E)** MS/MS spectrum of (3*S*)-deoxypumiloside **12** produced in *N. benthamiana* fed with FMN; diagnostic fragment ions are compared with those of the authentic standard. **(F)** LC-MS analysis of product **12** in reactions containing substrate strictosamide aglycone **4**, BaUGT, and FMN. EIC for (3*S*)-deoxypumiloside **12** (m/z [M+H]^+^ = 497.1918) are shown. **(G)** LC-MS analysis of camptothecoside **14** in *in vitro* reactions containing substrate strictosamide aglycone **4**, BaUGT, and FMN. EIC for camptothecoside **14** (m/z [M+H]^+^ = 495.1762) are shown. STD, standard; EV, empty vector.

Subsequently, the biosynthesis of camptothecin **9** from strictosamide aglycone **4** entails an intricate skeletal rearrangement, transitioning from a 6-5-6 to a 6-6-5 fused ring system, and was previously deduced to be initiated by the electron loss of the secondary amine (*39, 40*). We hypothesized that a flavin-containing monooxygenase (FMO) might catalyze this ring rearrangement and accordingly cloned and expressed several candidate *FMO* genes in *Nicotiana benthamiana* (Fig. 4C). When strictosamide aglycone **4** was co-infiltrated with FMN-the redox cofactor of FMO proteins-into *N. benthamiana* plants expressing FMO candidates as well as *N. benthamiana* plants without expressing FMO as controls, we did not detect the target product deoxypumiloside aglycone **5**. Unexpectedly, (3*S*)-deoxypumiloside **12** (*m/z* [M+H]^+^ = 497.1918), the corresponding glycosylated product of deoxypumiloside aglycone **5**, which have already had the 6-6-5 fused ring system, were observed in all FMO-expressing and control plants (Fig. 4D). Given that FMN has been reported to mediate electron-transfer-driven skeletal rearrangement (*39*), we speculated that the skeleton rearrangement from strictosamide aglycon **4** to (3*S*)-deoxypumiloside **12** is likely catalyzed by FMN (fig. S15). Indeed, when only strictosamide aglycon **4** and FMN were injected into the wide type *N. benthamiana*, LC-MS analysis also confirmed the formation of (3*S*)-deoxypumiloside **12** (Fig. 4E). These results indicated that FMN may trigger the skeleton rearrangement to form the 6-6-5 fused ring system of camptothecin **9**. The formation of (3*S*)-deoxypumiloside **12** instead of its deglycosylated product, deoxypumiloside aglycone **5**, from the substrate strictosamide aglycon **4** likely results from the glycosylation of deoxypumiloside aglycone **5** by endogenous tobacco glycosyltransferases. To investigate whether FMN can promote strictosamide **3** into (3*S*)-deoxypumiloside **12**, we incubated FMN and strictosamide **3** *in vitro*. Under this condition, instead of (3*S*)-deoxypumiloside **12**, only compound **11** was observed, compared with the quinoline moiety in camptothecin **9**, the quilolin-4(1H)-one moiety in compound **11** is markedly different (figs. S23, S24, and S38). Thus, we continued to focus our attention on the non-enzymatic 6-5-6 to 6-6-5 skeleton rearrangement from strictosamide aglycon **4**.

To exclude the potential background interference from *N. benthamiana*, we directly incubated FMN with **4** *in vitro* (Fig. 4F), none of the 6-6-5 ring product deoxypumiloside aglycone **5** or (3*S*)-deoxypumiloside **12** was detected. We speculated that the intermediate deoxypumiloside aglycone **5** without glucosyl moiety might be unstable under these reaction conditions. Indeed, when OpGH1 was incubated with (3*S*)-deoxypumiloside **12**, the deglycosylated product deoxypumiloside aglycone **5** accumulated at very low levels (fig. S16, A, B and E), even though the substrate was largely consumed (fig. S16D). We therefore sought to confirm the formation of deoxypumiloside aglycone **5** and camptothecoside aglycone **6** *via* detecting their more stable glycosylated product by *in vitro* reaction system. We screened a group of UDP-glycosyltransferases from various sources to find an enzyme capable of glycosylating these aglycones (fig. S17). When a multifunctional UDP-glycosyltransferase from *Bacillus subtilis*, BaUGT, together with sugar donor UDP-glucose, was incubated with strictosamide aglycone **4** and FMN reaction, (3*S*)-deoxypumiloside **12** (Fig. 4F) and camptothecoside **14** (Fig. 4G), the respective glycosylated correspondents of deoxypumiloside aglycone **5** and camptothecoside aglycone **6**, were clearly detected. The above results demonstrated that FMN could directly catalyze strictosamide aglycone **4** to form unstable products **5** and **6** that could be protected by further glycosyl modification. When (3*S*)-deoxypumiloside **12** was catalyzed by OpGH1, camptothecoside aglycone **6** was also detected (fig. S16F), implying that camptothecoside aglycone **6** may be produced by spontaneous reaction of the unstable intermediate deoxypumiloside aglycone **5** (fig. S16, A and C). Collectively, these results indicated that the glycoside hydrolase OpGH1 and the cofactor FMN act sequentially to convert strictosamide **3** to deoxypumiloside aglycone **5**, which spontaneously rearranges to camptothecoside aglycone **6** (Fig. 4A).

### OpSTR directly catalyzes the formation of strictosamide

Previous research reported that strictosidine synthase (OpSTR) from *O. pumila* could catalyze tryptamine **1** and secologanin **2** to generate strictosidine **10** (*30*). The lactam formation of strictosamide **3** from strictosidine **10** was supposed to be catalyzed by an unknown P450 (*41*). To discover possible gene involved in the formation of strictosamide **3**, we co-expressed *OpSTR* with eleven cortex-enriched, *OpSTR*-co-expressed P450 candidates in *N. benthamiana*. Unexpectedly, when tryptamine **1** and secologanin **2** were fed to *N. benthamiana* as substrates, two compounds could be detected in the *N. benthamiana* leaves expressing OpSTR alone, and their retention time and fragments matched those of the standard strictosamide **3** and strictosidine **10**, respectively (Fig. 5, B and C). The addition of P450 genes has no effect on the yield of the strictosamide **3** in the *N. benthamiana* (fig. S19). These results imply that either OpSTR could directly catalyze the formation of strictosamide **3** from tryptamine **1** and secologanin **2**, or that unknown native enzymes of *N. benthamiana* could catalyze the conversion of strictosidine **10** into strictosamide **3** (Fig. 5A). To further verify the function of OpSTR, we expressed OpSTR in *E. coli* and obtained the pure protein to catalyze tryptamine **1** and secologanin **2** *via in vitro* enzymatic reaction. Both strictosamide **3** and strictosidine **10** could be detected in the OpSTR reaction system (Fig. 5, D and E). Then, these two products were purified from large-scale reaction system and confirmed as strictosamide **3** and strictosidine **10** by NMR analysis, respectively (figs. S32, S36, and S37). These results clearly demonstrated that OpSTR alone can catalyze the formation of strictosamide **3**. However, OpSTR could not catalyze strictosidine **10** to form strictosamide **3**, indicating that strictosamide **3** is formed directly from tryptamine **1** and secologanin **2**, and that strictosidine **10** is not its precursor in this reaction (fig. S20). We speculate OpSTR may generate distinct transition states from substrates tryptamine **1** and secologanin **2**, thereby yielding two separate products strictosamide **3** and strictosidine **10** (fig. S21). The discovery of this novel function of OpSTR uncovers a long-term unknown mechanism underlying six-membered ring formation (lactam scaffold formation) during early camptothecin **9** biosynthesis (Fig. 1).

**Fig. 5.**
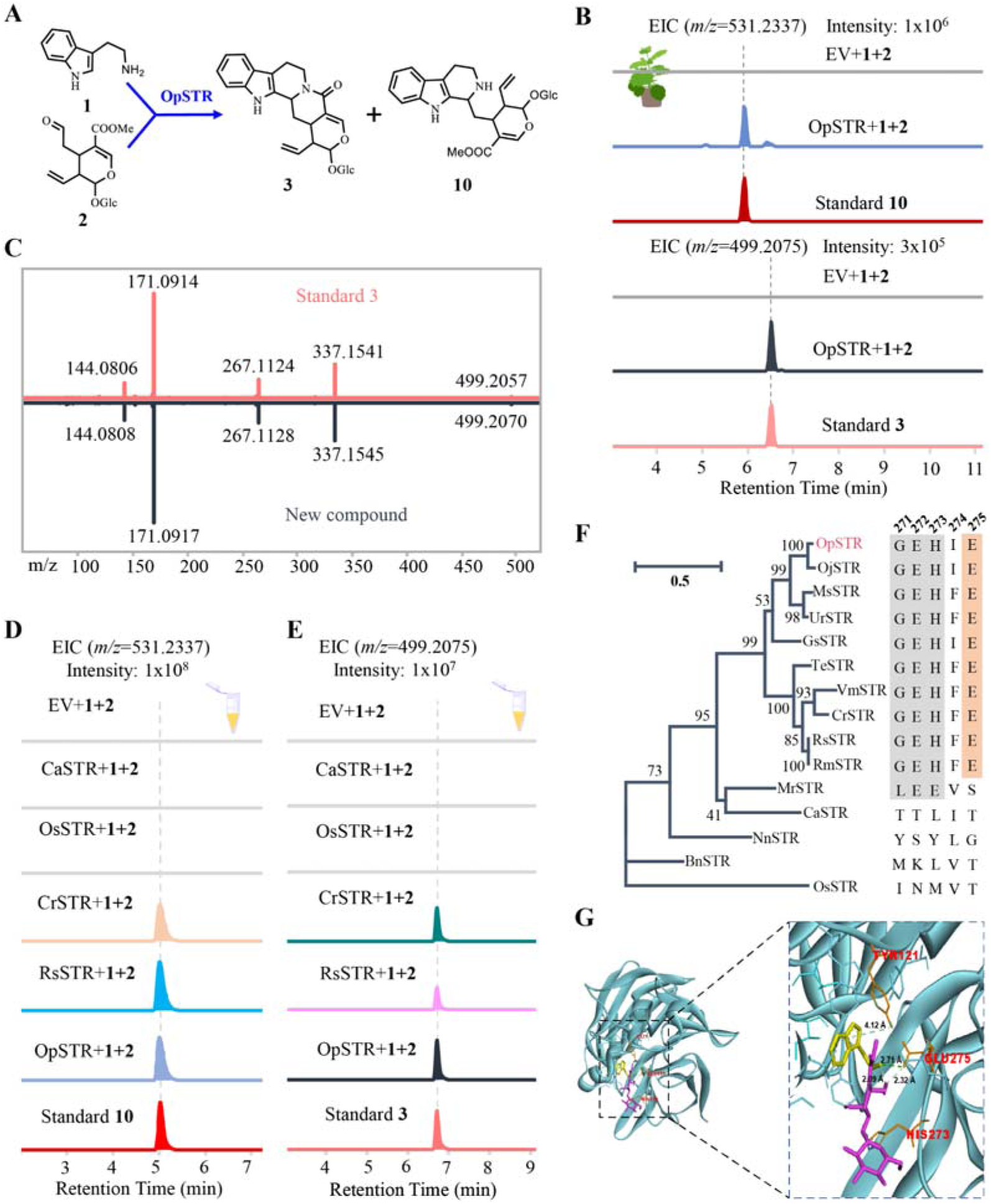
Biochemical characterization of *OpSTR*. **(A)** Reaction catalyzed by OpSTR (*O. pumila* strictosidine synthase). **(B)** LC-MS analysis of strictosidine **10** and strictosamide **3** produced in *N. benthamiana* expressing *OpSTR* and fed with tryptamine **1** and secologanin **2** as substrates. EIC for strictosidine **10** (m/z [M+H]^+^ = 531.2337) and strictosamide **3** (m/z [M+H]^+^ = 499.2075) are shown. EV, empty vector. **(C)** MS/MS spectrum of strictosamide **3** produced in *N. benthamiana* expressing *OpSTR* and fed with tryptamine **1** and secologanin **2** as substrates. Diagnostic fragment ions of the enzymatic product are consistent with those of the authentic standard. **(D)** LC-MS analysis of product of *in vitro* reaction using crude enzyme from STR orthologs of diverse plant species with tryptamine **1** and secologanin **2** as substrates. All STR proteins were expressed in *E. coli* and purified for *in vitro* assays. CrSTR, strictosidine synthase from *Catharanthus roseus*; RsSTR, from *Rauvolfia serpentina*; CaSTR, from *Camptotheca acuminata*; OsSTR, from *Oryza sativa*. EIC for strictosidine **10** (m/z [M+H]^+^ = 531.2337) are shown. **(E)** LC-MS analysis of product of *in vitro* reaction using crude enzyme from STR orthologs of diverse plant species with tryptamine **1** and secologanin **2** as substrates. EIC for strictosamide **3** (m/z [M+H]^+^ = 499.2075) are shown. The top trace shows the empty vector control. **(F)** Maximum likelihood phylogenetic tree of OpSTR and STR orthologs from diverse plant species, constructed using MEGA 7. Inset shows alignment of residues 271–275 of OpSTR and selected STR orthologs. **(G)** Molecular docking model of OpSTR in complex with substrates tryptamine **1** and secologanin **2**. Tryptamine **1** is shown in yellow; secologanin **2** is shown in purple. Residues making contacts with the substrates are highlighted in orange.

To investigate whether this novel function of STR is conserved in other plants, its homologous genes from several plants were selected for functional characterization. We found that CrSTR from *Catharanthus roseus* and RsSTR from *Rauvolfia serpentine* could catalyze the formation of strictosamide **3** and strictosidine **10**, while CaSTR and OsSTR from *C. acuminata* and *Oryza sativa*, respectively, failed to generate strictosamide **3** (Fig. 5, D and E). Phylogenetic analysis indicated CrSTR and RsSTR share a closer phylogenetic relationship with OpSTR compared with CaSTR and OsSTR (Fig. 5F). Sequence alignment revealed the amino acid E275 is conserved among the bifunctional active STRs (OpSTR, CrSTR, and RsSTR) (Fig. 5F), and molecular docking showed that the amino acids E275 may play a key role in enzymatic activity (Fig. 5G). We verified the function of E275 by mutating E275 to corresponding amino acids (S, T, and G) found in the non-bifunctional STR. LC-MS analysis indicated that neither strictosamide **3** nor strictosidine **10** could not be formed, suggesting that E275 plays a key role in the catalytic activity of STR (fig. S22).

### *De novo* biosynthesis of camptothecin in engineered *Saccharomyces cerevisiae*

Through the discovery of all the key missing enzymes mediating lactam ring formation and tailoring modifications, together with the identification of an FMN-triggered non-enzymatic skeleton rearrangement, we have uncovered the complete biosynthetic pathway of camptothecin **9**. To verify this pathway and explore a novel synthetic biology approach for camptothecin **9** production, we sought to reconstruct it for *de novo* biosynthesis in an engineered *S. cerevisiae* chassis (Fig. 6A). A series of metabolic engineering approaches were adopted in this process, including (1) overexpressing three mevalonate pathway genes, *ScERG12*, *ScERG8*, and *ScERG19*, together with a geraniol-producing mutant of *ScERG20* (*ScERG20*^F96W-N119W^) to augment the endogenous geranyl pyrophosphate (GPP) precursor pool; (2) The topoisomerase I gene from *O. pumila*, which confers tolerance to camptothecin, was overexpressed in order to avoid the toxicity of camptothecin **9** towards *S. cerevisiae* cell factory; (3) more than 20 heterologous and endogenous genes were overexpressed through iterative rounds of chromosomal integration (table S5). When the engineered *S. cerevisiae* CPT120 was cultivated in YPD medium supplemented with FMN (0.1 mM), camptothecin **9** was detected by LC-MS analysis with a titer of 0.55 μg/L (Fig. 6B).

**Fig. 6.**
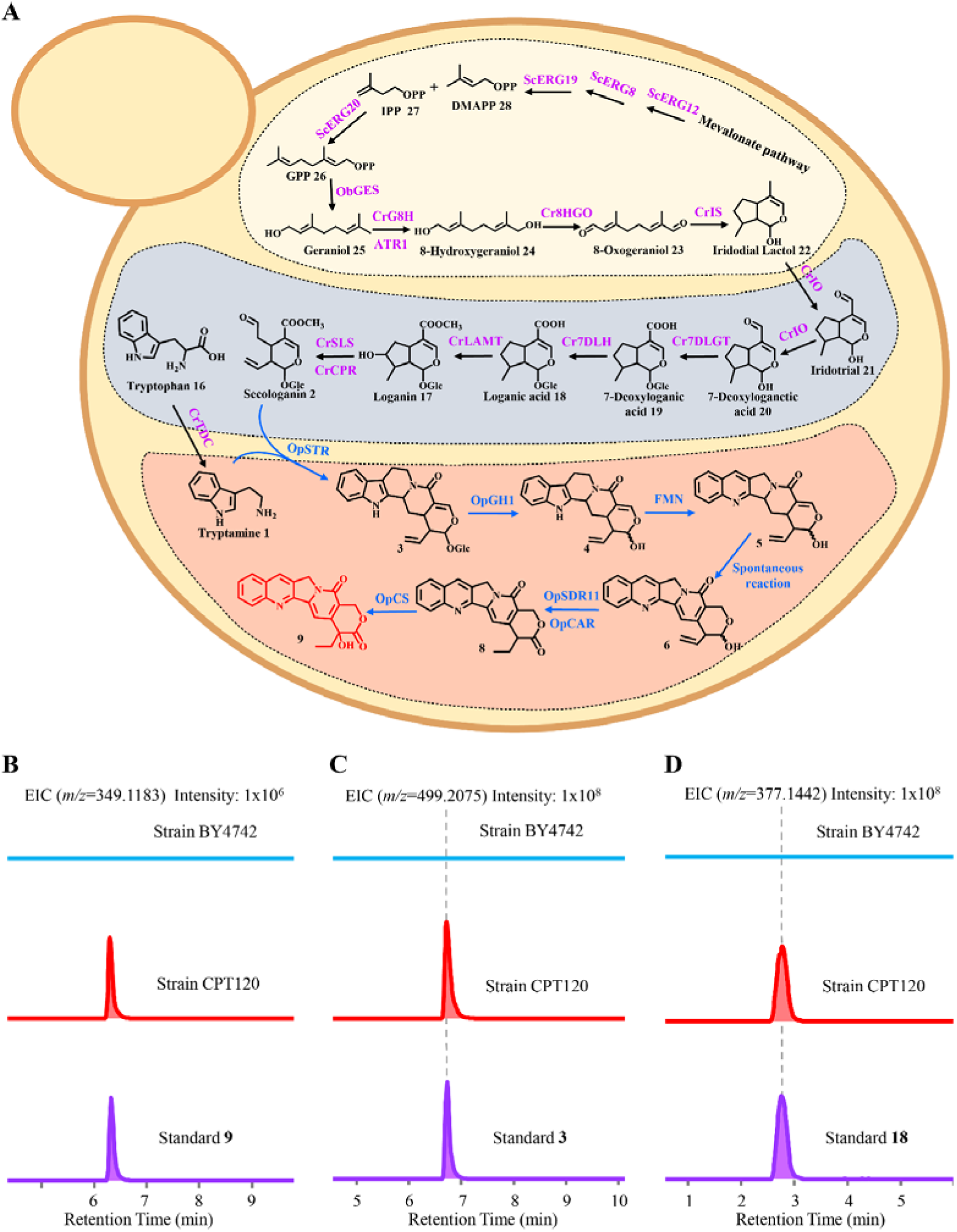
*De novo* biosynthesis of camptothecin 9 in *S. cerevisiae*. **(A)** Schematic of the engineered *S. cerevisiae* strain CPT120 for camptothecin **9** production. **(B)** LC-MS detection of camptothecin **9** in fermentation broth of *S. cerevisiae* strain CPT120. EIC for camptothecin **9** (m/z [M+H]^+^ = 349.1183) from the engineered strain (middle) and authentic standard (bottom) are shown. **(C)** LC-MS detection of strictosamide **3** in fermentation broth of *S. cerevisiae* strain CPT120. EIC for strictosamide **3** (m/z [M+H]^+^ = 499.2075) from the engineered strain (middle) and authentic standard (bottom) are shown. **(D)** LC-MS detection of loganic acid **18** in fermentation broth of *S. cerevisiae* strain CPT120. EIC for loganic acid **18** (m/z [M+H]^+^ = 377.1442) from the engineered strain (middle) and authentic standard (bottom) are shown. Abbreviations: ScERG12, mevalonate kinase from *S. cerevisiae*; ScERG8, phosphomevalonate kinase from *S. cerevisiae*; ScERG19, diphosphomevalonate decarboxylase from *S. cerevisiae*; ScERG20, GPP synthase from *S. cerevisiae*; ObGES, geraniol synthase from *Ocimum basilicum*; CrG8H, geraniol 8-hydroxylase from *C. roseus*; ATR1, NADPH-cytochrome P450 reductase from *Arabidopsis thaliana*; CrCPR, NADPH-cytochrome P450 reductase from *C. roseus*; Cr8HGO, 8-hydroxygeraniol oxidoreductase from *C. roseus*; CrIS, iridoid synthase from *C. roseus*; CrIO, iridoid oxidase from *C. roseus*; Cr7DLGT, 7-deoxyloganetic acid glucosyltransferase from *C. roseus*; Cr7DLH, 7-deoxyloganic acid hydroxylase from *C. roseus*; CrLAMT, loganic acid *O*-methyltransferase from *C. roseus*; CrSLS, secologanin synthase from *C. roseus*; CrTDC, tryptophan decarboxylase from *C. roseus*; OpSTR, strictosidine synthase from *O. pumila*; OpGH1, glycoside hydrolase 1 from *O. pumila*; FMN, flavin mononucleotide; OpCAR, camptothecoside aglycone reductase from *O. pumila*; OpSDR11, short-chain dehydrogenase/reductase 11 from *O. pumila*; OpCS, camptothecin synthase from *O. pumila*. Black arrows denote previously characterized enzymatic reactions, and blue arrows denote the reactions we characterized in this study.

Metabolite profiling of CPT120 also revealed the accumulation of several key pathway intermediates, including strctosamide **3**, camptothecoside aglycone **6**, deoxycamptothecin **8**, geraniol and loganic acid **18**, supporting the validity of the elucidated complete biosynthetic pathway of camptothecin **9** (Fig. 6, C and D). Moreover, the major product of OpSTR strictosidine **10**, which could not enter camptothecin **9** biosynthesis, accumulated to 10.91 mg/L, while strictosamide **3** accumulated to only 0.45 mg/L, suggesting that OpSTR-catalyzed formation of strictosamide **3** represents a primary bottleneck and motivating future efforts to improve camptothecin **9** production through enzyme and pathway engineering.

In summary, we verified the complete biosynthetic pathway of camptothecin **9** and its *de novo* biosynthesis through the construction of engineered microbial CPT120 producing camptothecin **9**. This proof-of-concept strain assembles the complete ≥20-step biosynthetic route from primary metabolism to camptothecin **9** production. The accumulation profiles of key intermediates delineate the principal flux bottlenecks and establish a tractable chassis for systematic metabolic engineering to drive camptothecin **9** titers toward industrially meaningful levels.

## Discussion

Camptothecin **9** biosynthesis has remained unresolved for five decades, owing to three challenging transformations: lactam scaffold formation, carbon-skeleton rearrangement, and late-stage tailoring (*43*). Advances in single-cell transcriptomics and spatial metabolomics now enable pathway discovery by linking specialized metabolites to cell-type-specific gene expression. In this study, by integrating MALDI-MSI, single-cell RNA sequencing and co-expression analysis, we identified cortex-associated metabolic and transcriptional signatures in *O. pumila*, which guided the discovery of the key missing genes in the camptothecin **9** pathway from a narrowed down candidate gene pool (Figs. 2 to 4). This spatially informed strategy provided the critical entry point for uncovering the enzymatic and chemical logic of camptothecin **9** biosynthesis efficiently and enabled its complete reconstruction.

The discovery of a previously unrecognized function of OpSTR revises the entry point into the camptothecin branch of monoterpene indole alkaloid metabolism. STR has been considered the canonical gatekeeper of MIA biosynthesis, uniquely producing strictosidine **10** from which approximately 3,000 alkaloids are derived (*44*). Our data show that OpSTR can also directly produce strictosamide **3** from the same substrates through divergent cyclization transition states (Fig. 5). This finding resolves a key uncertain lactam formation in early camptothecin **9** biosynthesis: the lactam-bearing strictosamide scaffold can arise directly at the STR step rather than through a separate downstream oxidation of strictosidine **10**. The identification of STR E275 as a conserved residue required for the bifunctional catalytic activity to produce strictosamide **3** and strictosidine **10** provides a molecular foothold for future studies of how sequence variation and active-site remodeling have driven functional divergence within the STR family. Thus, OpSTR expands the role of STR from a strictosidine-forming gatekeeper to a branch-point enzyme that initiates the camptothecin-specific lactam scaffold, with broader implications for the evolution of pathway branching and the biocatalytic diversification of alkaloid skeletons. CaSTR from *C. acuminata* could only catalyze the production of strictosidinic acid from tryptamine **1** and secologanic acid (*4, 30*) instead of the production of strictosidine **10** and strictosamide **3** from tryptamine **1** and secologanin **2** (Fig. 5), implying some plant species may produce strictosamide **3** *via* divergent biosynthetic route.

The 6-5-6 to 6-6-5 skeletal rearrangement is the key process within the camptothecin **9** biosynthesis (*28, 34*). We found that when combining strictosamide **3**, OpGH1 and FMN, FMN could directly catalyze the compound **4** generated by OpGH1 to form compound **6**, a product structurally much closer to camptothecin **9** (Fig. 4). We demonstrated that this skeleton rearrangement process could be achieved *via* a reaction without enzymes, which was triggered by free FMN, and is therefore mechanistically notable. Reduced flavin are known to act as free cofactors in other biosynthetic pathways (*40*) and rarely reported to take participate in this complex skeleton rearrangement process (fig. S15). During the preparing of this manuscript, this FMN-catalyzing skeleton rearrangement of strictosamide **3** was also reported by other researchers (*46*), they also found that FMN can convert strictosamide **3** to pumiloside **11**, as shown in our study (figs. S23, S24, and S38). However, the pumiloside **11** bearing a quilolin-4(1H)-one moiety as observed, which need several uncharacterized modifications to afford quinoline moiety in the target camptothecin 9. (3*S*)-deoxypumiloside **12** was not detected as expected when strictosamide **3** was incubated with FMN, we proposed the exist of free hydroxyl or glucosyl moiety at C21 is key to form a quinoline moiety or a quilolin-4(1H)-one moiety during the skeleton rearrangement catalyzed by FMN. The mechanism underlying this reaction is unknown. Isotope feeding experiment and theory calculation analysis were necessary to unearth it. Currently, the efficient of this process catalyzed by FMN is low, it is necessary to confirm whether some enzymes may catalyze or accelerate this process in future.

In the late stage of biosynthetic pathway for camptothecin **9**, three enzymes (OpCAR, OpSDR11 and OpCS) from different families achieved the tailoring modification process including reduction, oxidation and hydroxylation reactions (Fig.3), implying the complexity of camptothecin **9** biosynthesis. Although two parallel routes from camptothecoside aglycone **6** to deoxycamptothecin **8** were observed (Fig. 3 and S13), the strict substrate specificity of OpCS for deoxycamptothecin **8**, excluding aglycone analogs and other derivatives, established the C20 hydroxylation as the final committed step (table S2). Meanwhile, recently more studies indicated that the biosynthetic pathway of camptothecin **9** is not linear (*28, 46*). The elucidation of parallel biosynthetic pathway to camptothecin **9** and discovery of related enzymes will contribute to the construction and optimization of cell factory for camptothecin **9** in microorganism or plant chassis.

The *de novo* production of camptothecin **9** in *S. cerevisiae* at 0.55 μg/L establishes a proof of principle that this complex plant alkaloid could be formed in a microbial cell factory (Fig. 6). Detection of camptothecin **9** together with key intermediates confirms that the identified complete biosynthetic route is functional even in a heterologous microorganism chassis. This achievement in this study removes the barrier for heterologous biosynthesis of camptothecin **9** using microbial fermentation and paves a way for the sustainable production of camptothecin **9** and its derivatives. Analysis of limiting steps and enzyme engineering for enzymes involved in the limiting steps in microorganism cell factory will benefit the yield improvement of target compound camptothecin and boost the manufacturing of it and its derivatives.

## Acknowledgments

Authors sincerely appreciate Professor Yu Zhang from CAS Center for Excellence in Molecular Plant Sciences (CEMPS) for his valuable advices on the preparation of this manuscript and Yulu Zhang, Yuanyuan Gao, Wenxian Lan, Lianyan Jing, and Xiaoyan Xu from the Core Facility Centre of CEMPS for mass spectrometry or NMR analyses assistance.

## Author contributions

Z.Z., P.W., C.Y. and Y.W. supervised and coordinated the experiments. T.Z. and Y.X. performed the most of functional characterization including gene mining and cloning, protein expression, catalytic reaction and product analyses. K.C. conducted the co-expression analysis and help T.Z. and Y.X. with the functional characterization. S.W. and X.Y. performed the bioinformation analysis and help T.Z. and Y.X. with the mining of candidate genes. J.Z. speculated the catalytic mechanism of FMN and help T.Z. and Y.X. with the NMR analysis. T.Z., P.W., C.Y., and Z.Z. wrote the manuscript.

## Data and materials availability

Coding sequences for the functional *Ophiorrhiza pumila* genes characterized in this study has been deposited in the National Center Biotechnology Information (NCBI).

