## Supplementary material for "Complete elucidation and heterologous reconstruction of the biosynthetic pathway of camptothecin": SI

**The PDF file includes:**

Materials and Methods

Supplementary Text

Figs. S1 to S40

Tables. S1 to S4

**Materials and Methods**

Plant materials, strains, and chemicals

*Ophiorrhiza pumila* plants were collected from Longjuan Town, Anxi County, Quanzhou City, Fujian, China, and then frozen for RNA preparation. *Camptotheca acuminata* plants were grown in a greenhouse under a 16 h light/8 h dark cycle at 28℃. *Escherichia coli* strain TOP10 and *Agrobacterium tumefaciens* strain GV3101 were purchased from Shanghai Weidi Biotechnology Co., Ltd. Chemical standards for loganic acid **18**, tryptamine **1**, strictosamide **3**, and pumiloside **11** were purchased from TargetMol (USA). The strictosidine **10** standard was purchased from Shanghai Yiqi Biotechnology Co., Ltd. Acetosyringone and nicotinamide adenine dinucleotide phosphate (NADPH) were purchased from Shanghai Yuanye Biological Co., Ltd. Flavin mononucleotide (FMN) was obtained from Shanghai Titan Technology Co., Ltd. The camptothecoside **14** standard was obtained from *C. acuminata* seeds, using an isolation method performed as previously reported (*48*). Camptothecoside aglycone **6** was obtained from the large-scale reaction of OpGH1 with camptothecoside **14** (figs. S18 and S33). Chemical reagents such as methanol, acetonitrile, and dimethyl sulfoxide were of HPLC grade.

Camptothecin analysis of *O. pumila*

Three one-year-old *O. pumila* plants were washed with tap water and sterile water, three times each, to ensure the removal of impurities such as soil. Each plant was divided into three tissue types—root, stem, and leaf—and these air-dried tissues were subsequently ground into powder. A 0.1 g sample of powder was thoroughly soaked in 10 mL methanol at 30°C for 3 h, followed by centrifugation at 12,000 rpm, and the sediment was extracted with methanol again. The combined supernatants were combined and stored at −20°C for liquid chromatography mass spectrum (LC-MS) analysis. LC-MS analysis was performed on a Q Exactive Plus mass spectrometer coupled with a UPLC-Q-Orbitrap-HRMS system, and instrument parameters were set as previously described with minor modifications (*28*). Compounds were separated on a Welch BioUltimate-C18 column (2.1 × 100 mm, 2.7 μm) at a flow rate of 0.35 mL/min. The elution gradient was as follows: 0–1 min, 10% acetonitrile; 1–11 min, 10%–55% acetonitrile; 11–11.5 min, 55%–95% acetonitrile; 11.5–15 min, 95% acetonitrile; 15–15.1 min, 95%–10% acetonitrile; 15.1–18 min, 10% acetonitrile.

Mass spectrometry imaging analysis

The roots of one-year-old *O. pumila* were frozen and sectioned at a thickness of 20 μm using a Leica CM1950 cryostat (Leica Microsystems GmbH, Wetzlar, Germany) at −20°C. The tissue sections were then placed on electrically conductive slides coated with indium tin oxide (ITO) and dried in a vacuum desiccator for 30 min. Desiccated tissue sections were sprayed using a TM-Sprayer with 2,5-dihydroxybenzoic acid (DHB, 15 mg/mL in 90% acetonitrile). The flow rate of the sprayer was set to 0.1 mL/min, with a temperature of 60°C and a pressure of 6 psi. The matrix was sprayed onto the tissue sections in twenty passes, with 5 s of drying time between each pass. Compounds were desorbed, ionized, and imaged using a prototype Bruker timsTOF flex MS system (Bruker Daltonics, Bremen, Germany) equipped with a 10 kHz SmartBeam 3D laser. Laser power was set to 80%, and mass spectra were acquired in positive mode throughout the experiment. Mass spectral data were collected over a mass range of m/z 50–1300 Da. The imaging spatial resolution was 20 μm, and each spectrum consisted of 400 laser shots. Matrix-Assisted Laser Desorption Ionization (MALDI) mass spectrometry data were analyzed using SCiLS Lab software and normalized using the root mean square; normalized intensity was used to display the signal intensity in each image (*49*). MS/MS fragmentation spectra obtained using the timsTOF flex MS system were used to confirm the identities of the detected metabolites.

Isotope feeding experiment of *O. pumila* hairy root

The *O. pumila* hairy roots without genetic modification were obtained by Agrobacterium infection as previously reported (*50*). Briefly, 1 cm-long stems cut from sterile *O. pumila* plants were incubated on solid B5 medium for 48 h in the dark. *Agrobacterium rhizogenes* strain C58C1 was cultured at 28°C until the OD_600_ reached 0.5; the cells were then collected by centrifugation (5,000 rpm, 10 min) and resuspended in liquid B5 medium, followed by cultivation for 30 min at 28°C and 110 rpm. The stems were then immersed in the *A. rhizogenes* suspension for 8 min, blotted dry on sterile paper, and cultured on solid B5 medium for 48 h. The stems were then cultured on solid B5 medium containing 300 mg/L carbenicillin to induce hairy roots formation, and the resulting hairy roots were subcultured sequentially on solid B5 medium containing 200, 100, and 0 mg/L carbenicillin. Once the hairy roots reached a stable growth stage on solid B5 medium, 3 cm long roots were excised from the petri dish and cultured in liquid B5 medium. After 14 days, 2 mg of deuterium-labeled d4-tryptamine, dissolved in 1 mL ethanol and filtered through a 0.22 μm membrane filter, was added to the hairy root culture. Thirty days later, the hairy roots were blotted dry with absorbent paper and dried in an oven at 35°C until a constant dry weight was achieved. Subsequently, 0.1 g of hairy roots was ground into powder and extracted three times with 1 mL methanol each time. The combined supernatants were collected after centrifugation (12,000 rpm, 10 min) and transferred to an injection vial for LC-MS analysis.

Single-cell RNA sequencing

Protoplast preparation was performed using the GEXSCOPE® Single Cell RNA-seq Kit, purchased from Singleron Biotechnologies Co., Ltd. The 45-day-old *O. pumila* hairy roots were sliced into 1–2 mm long strips and quickly transferred into a sterile dish containing 20 mL of enzyme digestion solution. The dish was kept under vacuum for 30 min, after which it was shaken at 60 rpm in the dark until digestion was complete. The protoplast solution was filtered through a cell strainer and centrifuged at 100 × g for 5 min, and the supernatant was discarded. Subsequently, the protoplasts were washed twice with washing buffer and stained with fluorescein diacetate (FDA) to assess viability. A protoplast suspension at a concentration of 3.0–3.5 × 10⁵ cells/mL was loaded onto a microfluidic chip, and scRNA-seq library construction was performed according to the manufacturer's instructions (Singleron Biotechnologies). An Illumina NovaSeq 6000 with 150 bp paired-end reads was used to sequence the scRNA-seq libraries, and the raw reads were analyzed to generate gene expression profiles using CeleScope v2.0.7 (Singleron Biotechnologies). Briefly, unique molecular identifiers (UMIs) and barcodes were extracted from R1 reads and error-corrected (*51*). Poly-A tails and adapter sequences were trimmed from R2 reads, and the trimmed R2 reads were aligned against the *O. pumila* transcriptome (https://pumila.kazusa.or.jp/) using STARSolo (STAR v2.7.11a). Successfully Assigned Reads (SAR) sharing the same cell barcode, UMI, and gene were aggregated to construct the gene expression matrix for downstream analysis.

Single-cell RNA-seq analysis

Quality control, dimensionality reduction, and clustering were performed using Scanpy v1.9.3 (*52*) in Python 3.10. For each sample, the expression matrix was filtered according to the following criteria: (1) cells with fewer than 200 detected genes or with gene counts in the top 2% were excluded; (2) cells in the top 2% of UMI counts were removed; and (3) genes expressed in fewer than 5 cells were excluded. Following filtering, the retained cells were used for downstream analyses. Raw counts were normalized to total counts per cell and subsequently log-transformed to generate a normalized data matrix. The top 2,000 variable genes were selected using the 'seurat_v3' flavor setting. Principal component analysis (PCA) was performed on the scaled variable gene matrix, and the top 2,000 principal components were retained for clustering and dimensionality reduction. Potential ambient RNA contamination was estimated and removed using DecontX v1.6.1 (*53*). To correct for batch effects between samples, Harmony v0.0.6 was applied to the top 2,000 principal components derived from PCA (*54*). Cell clusters were identified using the Louvain algorithm with a resolution parameter of 1.2 and visualized using Uniform Manifold Approximation and Projection (UMAP). To identify differentially expressed genes (DEGs), the Wilcoxon rank-sum test was applied using the scanpy.tl.rank_genes_groups function with default parameters. DEGs were defined as genes expressed in more than 10% of cells in either comparison group and with an average log10 fold change (logFC) > 1.5. Statistical significance was assessed using Benjamini–Hochberg adjusted p-values, with a significance threshold of p < 0.05. The volcano plot was generated using the ggplot2 package in R. Cell types were assigned to each cluster based on the expression of canonical marker genes from Arabidopsis thaliana. Sequence homology between the protein sequences of the detected genes and those of *A. thaliana* was assessed by sequence alignment. The canonical markers used and their corresponding cell types are listed in table S3.

RNA sequencing and analysis

The 1-year-old *O. pumila* plant was washed with sterile water and separated into three tissues (leaf, stem, and root). The tissues were then rapidly frozen in liquid nitrogen and stored at −80°C. RNA was extracted using the RNAprep Pure Plant Plus Kit (Tiangen Biotech Co., Ltd.) and reverse-transcribed into cDNA using the PrimeScript™ RT Reagent Kit (Takara). The cDNA was fragmented into 150–800 bp segments for library construction, and sequencing was performed on the Illumina platform. The normalized expression matrix was imported into R version 4.3.2, and the gene *OpCAR* was selected as the bait gene. Pearson correlation coefficients between *OpCAR* and all other genes were calculated using the ‘cor()’ function with the ‘pearson’ method. Genes exhibiting strong positive correlations with *OpCAR* (Pearson's r ≥ 0.85) were identified as candidate co-expressed genes, and the bait gene *OpCAR* was excluded from the resulting candidate genes list. An edge table was subsequently generated using *OpCAR* as the source node and the candidate co-expressed genes as target nodes, along with the corresponding correlation coefficients. The resulting edge table was imported into Cytoscape for visualization of the co-expression network (*55*).

Gene cloning and vector building

Plant RNA was extracted from fresh leaves, and reverse transcription was performed using the PrimeScript™ RT Reagent Kit. Candidate genes from *O. pumila* were cloned from *O. pumila* cDNA using PrimeSTAR Max DNA Polymerase (TaKaRa). The strictosidine synthase genes *CaSTR*, *CrSTR*, and *OsSTR* were cloned from *Camptotheca acuminata*, *Catharanthus roseus*, and *Oryza sativa*, respectively (table S1). The gene *RsSTR*, codon-optimized for *E. coli*, was synthesized by Shanghai Rui Mian Biological Technology Co., Ltd. All genes were cloned into the appropriate vectors using the Hieff Clone Universal One Step Cloning Kit (Yeasen). Genes intended for functional characterization in *N. benthamiana* were cloned into the plasmid pCAMBIA2301 between the *BamH*I and *Sal*I sites, and the resulting constructs were transformed into *A. tumefaciens* strain GV3101 for transient expression. Genes intended for functional characterization in *E. coli* were amplified using gene-specific primers containing homologous arm sequences and then cloned into the pCold-TF vector between the *Nde*I and *Sal*I sites.

Functional characterization in *Nicotiana benthamiana*

The culture, collection, and washing of *A. tumefaciens* cells harboring the genes described above were performed as previously reported (*25, 28*). Briefly, the cells were cultured in Luria–Bertani (LB) medium at 28°C until the OD_600_ value exceeded 1.0 and harvested by centrifugation at 4,000 rpm. The cells were washed and resuspended in 5 mL of infiltration buffer (50 mM MES, 0.15 mM acetosyringone, 10 mM MgCl₂, pH 5.6). When testing the function of a single gene, the cell suspension was diluted to a final OD_600_ of 0.8 for infiltration. For simultaneous characterization of multiple genes, the *A. tumefaciens* suspensions were mixed such that the final OD_600_ of each strain was 0.3. The cells were then incubated at 28°C for 3 h and injected into *N. benthamiana* leaves for transient expression. *A. tumefaciens* harboring the empty pCAMBIA2301 vector was used as the negative control and each experiment was performed in triplicate. After four days, substrates were dissolved in MES buffer at a final concentration of 0.1 mM and injected into the *N. benthamiana* leaves for the catalytic reaction. Three days later, the *N. benthamiana* leaves were collected, ground, and extracted with methanol to obtain the crude extract. After centrifugation at 4,000 rpm for 5 min, 1 mL of the supernatant was dried using a rotary evaporator and reconstituted in 200 μL methanol for mass spectrometry analysis.

Protein structure prediction and molecular docking of STR

The three-dimensional structure of STR was predicted using AlphaFold (*56*). Molecular docking of tryptamine **1** and secologanin **2** into the predicted STR structure was performed using Discovery Studio. The protein was prepared by removing water molecules, adding hydrogen atoms, and applying the CHARMm force field. Ligand structures were obtained from PubChem and energy-minimized. Docking was conducted using the CDOCKER protocol, with the binding site defined based on the reported active site residues (*57*). The lowest-energy docked pose was selected for each ligand. Visualization of protein–ligand interactions, including hydrogen bonds and hydrophobic contacts, was performed using Discovery Studio.

Protein expression in *Escherichia coli*

E. *coli* cells harboring the vectors described above were first cultured in 10 mL LB medium with ampicillin (100 μg/mL) at 37°C until the OD_600_ value reached 2.0, and then inoculated into a 500 mL flask containing 100 mL LB medium at 37°C and 220 rpm. When the OD_600_ value reached 0.6, the flask was placed on ice for 10 min, and isopropyl-β-d-thiogalactoside (IPTG) was added to a final concentration of 0.5 mM. The cells were then cultured at 16°C and 120 rpm to induce protein expression. After 16 h, the cells were collected by centrifugation (4,000 rpm, 5 min) and resuspended in 5 mL PBS buffer (50 mM, pH 7.2). The cells were then disrupted by sonication and centrifuged (4,000 rpm, 5 min) to obtain the supernatant as the crude enzyme extract. Subsequently, the crude enzyme was purified using Ni-NTA His-Tag Purification Agarose (MedChemExpress) according to the manufacturer's instructions, and protein purity was assessed by sodium dodecyl sulfate–polyacrylamide gel electrophoresis (SDS-PAGE). The purified enzyme was dissolved in PBS buffer (50 mM, pH 7.0, 10% glycerol) to a final concentration of 0.1 mg/mL, flash-frozen in liquid nitrogen, and stored at −80°C.

Functional analysis of protein in vitro

OpSTR: For functional characterization of OpSTR in vitro, substrates tryptamine **1** and secologanin **2** were each added to a final concentration of 1 mM and mixed with 200 μL of purified OpSTR protein, and each experiment was performed in triplicate. The reaction was incubated at 30°C for 3 h and quenched by adding an equal volume of methanol. After mixing and centrifugation at 12,000 rpm for 5 min, the supernatant was collected and stored at −20°C for LC-MS analysis. Functional characterization of the other STRs was performed as described for OpSTR, and the TF-tag protein expressed from the empty vector was used as the negative control.

OpGH1: When testing the catalytic activity of OpGH1, the substrates strictosamide **3**, (3*S*)-deoxypuniloside **12**, camptothecoside **14**, and pumiloside **11** were each added to 200 μL of purified OpGH1 protein to a final concentration of 0.1 mM. The reaction was incubated at 30°C for 4 h and stopped by adding 400 μL of methanol. The reaction conditions for the TF-tag protein were identical to those for OpGH1 and served as the negative control. The supernatant collected by centrifugation was immediately stored at −20°C for subsequent LC-/MS analysis.

OpCAR: Camptothecoside aglycone **6** (0.1 mM) and NADPH (1 mM) were mixed with 200 μL of OpCAR protein to initiate the reaction, which was incubated at 30°C for 4 h. An equal volume of methanol was then added to stop the reaction, and the supernatant was collected after centrifugation (12,000 rpm, 5 min) and stored at −20°C for subsequent LC-MS analysis. Cells expressing the empty vector were used as the negative control.

OpSDR: Purified OpSDR protein (200 μL) containing NADPH (1 mM) was incubated with camptothecoside aglycone **6** (0.1 mM) at 30°C for 4 h. The reaction was then stopped by adding 200 μL of methanol, and the supernatant was collected after centrifugation (12,000 rpm, 5 min) for subsequent LC-MS analysis. To produce deoxycamptothecin **8** from camptothecoside aglycone **6**, 200 μL of the OpSDR reaction supernatant was evaporated to dryness and reconstituted in 200 μL of crude OpCAR protein solution containing NADPH (1 mM). After a 4 h reaction at 30°C, 200 μL of methanol was added to quench the reaction, followed by centrifugation (12,000 rpm, 5 min) and LC-MS analysis. Reactions using crude protein from the empty vector served as the negative control, and each experiment was performed in triplicate.

OpCS: Crude OpCS protein (200 μL) was combined with deoxycamptothecin **8** (30 μM), FeSO₄ (300 μM), L-ascorbic acid (300 μM), and α-ketoglutaric acid (300 μM) in PBS buffer. After a 24 h reaction at 30°C, 200 μL of methanol was added to stop the reaction, followed by thorough mixing. The supernatant was collected by centrifugation at 12,000 rpm for 5 min and stored at −80°C for subsequent analysis. The empty pCold-TF vector was used as the negative control, and each experiment was performed in triplicate.

Standard reactions of flavin mononucleotide

The standard reaction was conducted in 100 μL of PBS buffer (50 mM, pH 8.0) containing strictosamide **3** (0.5 mM) and flavin mononucleotide (FMN, 1 mM). Reactions without FMN were performed as the negative control. Centrifuge tubes containing the reaction mixture were placed in a water bath at 30°C for 24 h, after which 100 μL of methanol was added to stop the reaction. After centrifugation at 12,0 00 rpm for 5 min, the supernatant was collected and subjected to LC-MS analysis.

For the combined functional characterization of FMN, OpGH1, and BaUGT, strictosamide **3** (1 mM) and FMN (1 mM) were added to 200 μL of crude OpGH1 protein in Tris buffer (50 mM, pH 7.2). After a 6 h reaction, 200 μL of methanol was added to quench the reaction, and the supernatant obtained by centrifugation (12,000 rpm, 5 min) was evaporated to dryness. The dried residue was then reconstituted in 200 μL of crude BaUGT protein solution (Tris buffer, 50 mM, pH 7.2) containing uridine diphosphate glucose (UDPG) at a final concentration of 2 mM. The reaction was conducted at 30°C for 12 h and stopped by adding 200 μL of methanol, followed by centrifugation at 12,000 rpm for 5 min. The entire supernatant was evaporated to dryness at 30°C and reconstituted in 100 μL of methanol. After centrifugation at 12,000 rpm for 5 min, the supernatant was stored at −20°C for subsequent LC-MS analysis and each experiment was performed in triplicate.

Construction and analysis of the *S. cerevisiae* strains producing Camptothecin **9**

Genome editing of the strains producing camptothecin **9** was performed as previously reported (19). Briefly, genes were integrated into the chromosomes of *S. cerevisiae* BY4742 using the CRISPR/Cas9 system. The gene sequences of *ObGES, CrSLS, CrIS,* and *OpGH1* were codon-optimized for *S. cerevisiae*. Homologous arms (donor DNA fragments) were added to both ends of each expression cassette (promoter, gene, and terminator) by PCR (table S5), and the expression cassettes were integrated into the genome by homologous recombination. The expression cassette was transformed into *S. cerevisiae* using the LiAc/SS carrier DNA/PEG method, and the transformants were identified by PCR and sequencing. To evaluate camptothecin **9** production, the target strains were cultured in 10 mL of yeast extract peptone dextrose (YPD) medium with 0.1 mM FMN for 5 days (30°C, 250 rpm). The fermentation broth was extracted three times with an equal volume of ethyl acetate, and the extract was dried under vacuum. The dried extract was then reconstituted in 1 mL of methanol, and the supernatant obtained by centrifugation was stored at −20°C for LC-MS analysis.

Purification and NMR analysis of reaction products

The reaction product was first separated by silica gel column chromatography to remove most impurities, and the fractions eluted with petroleum ether at different concentrations were analyzed by high-performance liquid chromatography (HPLC) and mass spectrometry. The fraction containing the target compound was evaporated to dryness and reconstituted in methanol, followed by centrifugation (12,000 rpm, 5 min). The supernatant was then injected onto a preparative HPLC system equipped with an Agilent system and a Welch Ultimate XB-C18 column. Acetonitrile and ultrapure water were used as the mobile phase at a constant flow rate of 10 mL/min, and the product was monitored at 254 nm. The collected fraction was evaporated to dryness and dissolved in an appropriate deuterated solvent based on solubility. ¹H NMR and ¹³C NMR spectra were recorded on a Bruker AM-400 (400 MHz) or AM-500 (500 MHz) spectrometer in CDCl₃, CD₃OD, or (CD₃)₂SO. ¹H NMR chemical shifts (δ) of are reported in ppm relative to TMS (δ = 0.00 ppm) or DMSO-d_6_ (δ = 2.50 ppm). ¹³C NMR chemical shifts (δ) of are reported in ppm relative to CDCl₃ (δ = 77.16 ppm), CD₃OD (δ = 49.00 ppm), or (CD₃)₂SO (δ = 39.52 ppm).

Chemical synthesis of deoxypumiloside

Chemical synthesis route of (3*S*)-deoxypumiloside **12** and (3*R*)-deoxypumiloside **13** (*58*):

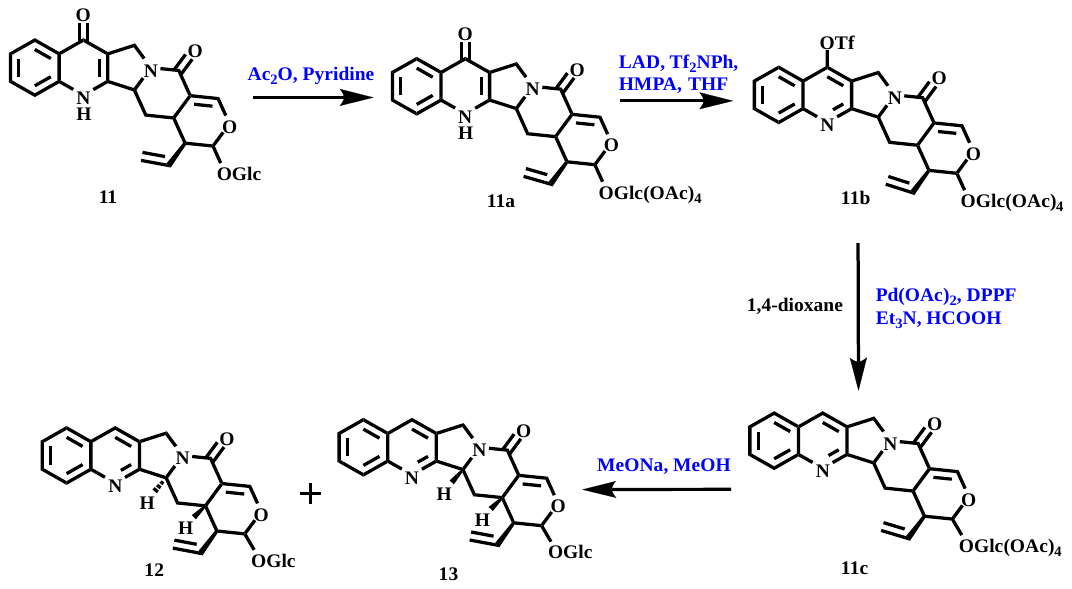

General: The solution of compound **11** (200 mg, 0.39 mmol) in pyridine (3.0 mL) and acetic anhydride (3.0 mL) was stirred overnight at room temperature. The solvent was removed in vacuum, and the residue was purified by C18 silica gel chromatography (CH_3_CN/H_2_O = 65/35) to obtain tetraacetate derivative **11a** as yellow solid (240 mg, 94% yield). The solution of compound **11a** (230 mg, 0.34 mmol) and HMPA (119 μL, 0.68 mmol) in anhydrous THF (11 mL) was cooled to –78 ℃, LDA (1.0 M in THF, 0.68 mL) was dropped and the resulted reaction system was stirred for 1 hour at the same temperature. Then the THF solution (1.1 mL) of N-phenyltriﬂuoro-methanesulfonimide (243 mg, 0.68 mmol) was added dropwise. The reaction system was stirred for 30 minutes, then warmed to room temperature gradually over1 h, and quenched with cold water. The organic solvent was removed under vacuum and the water phase was extracted with CH_2_Cl_2_ (3 x 50 mL), the combined organic phase was dried over anhydrous Na_2_SO_4_ and concentrated under vacuum. The residue was purified by C18 silica gel chromatography (CH_3_CN/H_2_O = 80/20) to obtain 7-triﬂate derivative **11b** as yellow solid (154 mg, 56% yield). To the solution of compound **11b** (150 mg, 0.18 mmol) in anhydrous 1, 4-dioxane (15mL), Pd (OAc)_2_ (15 mg, 0.05 mmol), DPPF (43 mg, 0.077 mmol), HCOOH (28 μL, 0.74 mmol) and Et_3_N (250 μL, 0.05 mmol) were added successively under N_2_ atmosphere at room temperature. The resulting mixture was heated at 60℃ for 30 minutes, then cooled to room temperature, quenched with cold water, and extracted with CH_2_Cl_2_. The combined organic phase was washed with 5% aqueous Na_2_CO_3_ and brine, then dried over anhydrous Na_2_SO_4_. The solvent was removed under vacuum and the resulting residue was purified by silico chromatography (DCM/MeOH = 20/1) to obtain compound **11c** as solid (80 mg, 66% yield). To the solution of compound **11c** (80 mg, 0.12 mmol) in MeOH (15 mL), MeONa (50% in MeOH, 79 mg, 0.73 mmol) was added at 0℃. The resulting reaction system was stirred at room temperature for 3 h. The solvent was removed in vacuum and the resulting residue was purified on silico chromatography (DCM/MeOH = 5/1) to obtain (3*S*)-deoxypumiloside **12** (12 mg, 20.2% yield) and (3*R*)-deoxypumiloside **13** as solid (10 mg, 16.8% yield) (figs. S39 and S40).

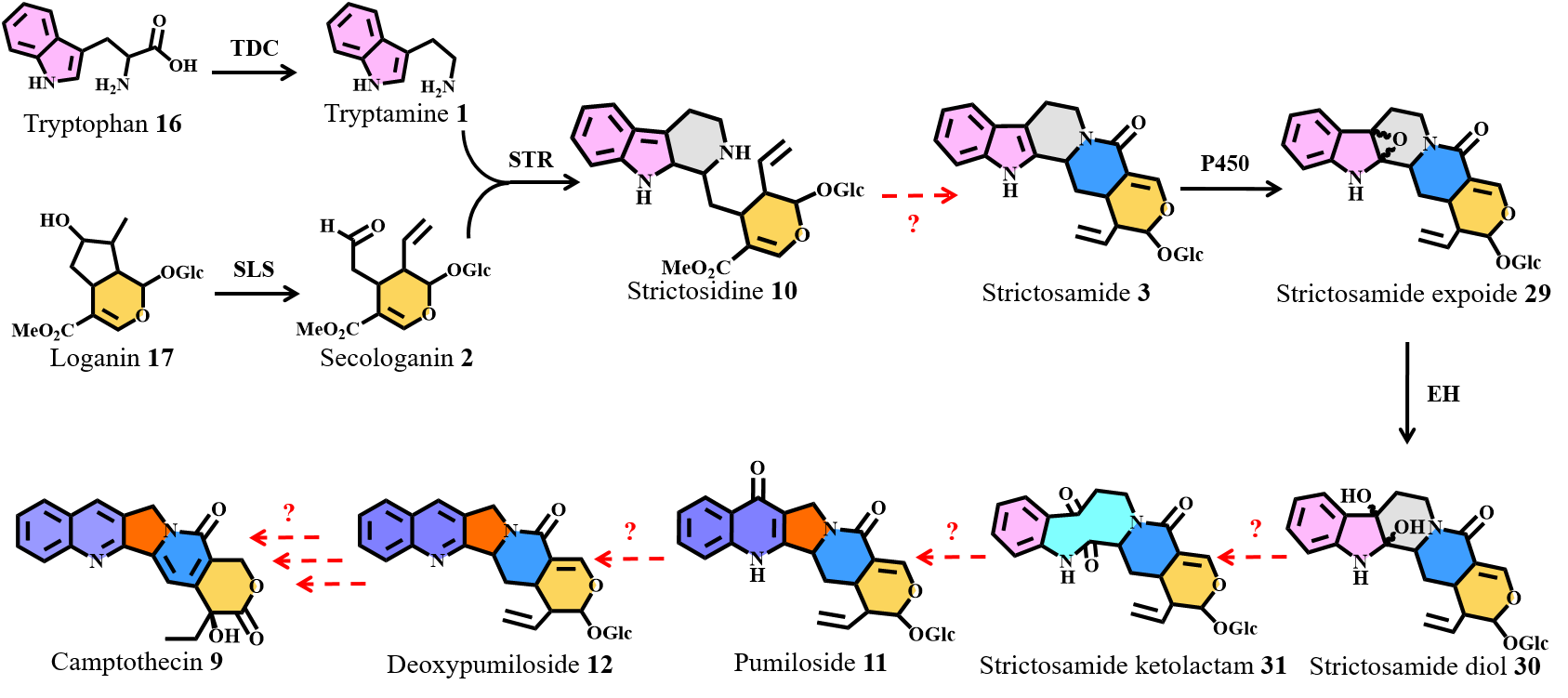

**Fig. S1. Proposed biosynthetic pathway of camptothecin*.*** Black solid arrows indicate reactions that have been identified previously; red dashed arrows indicate proposed but uncharacterized reactions. The biosynthesis of tryptamine **1** and secologanin **2** has been elucidated. The enzymes responsible for five-membered ring formation, skeleton rearrangement, and downstream tailoring steps leading to camptothecin **9** remain to be identified.

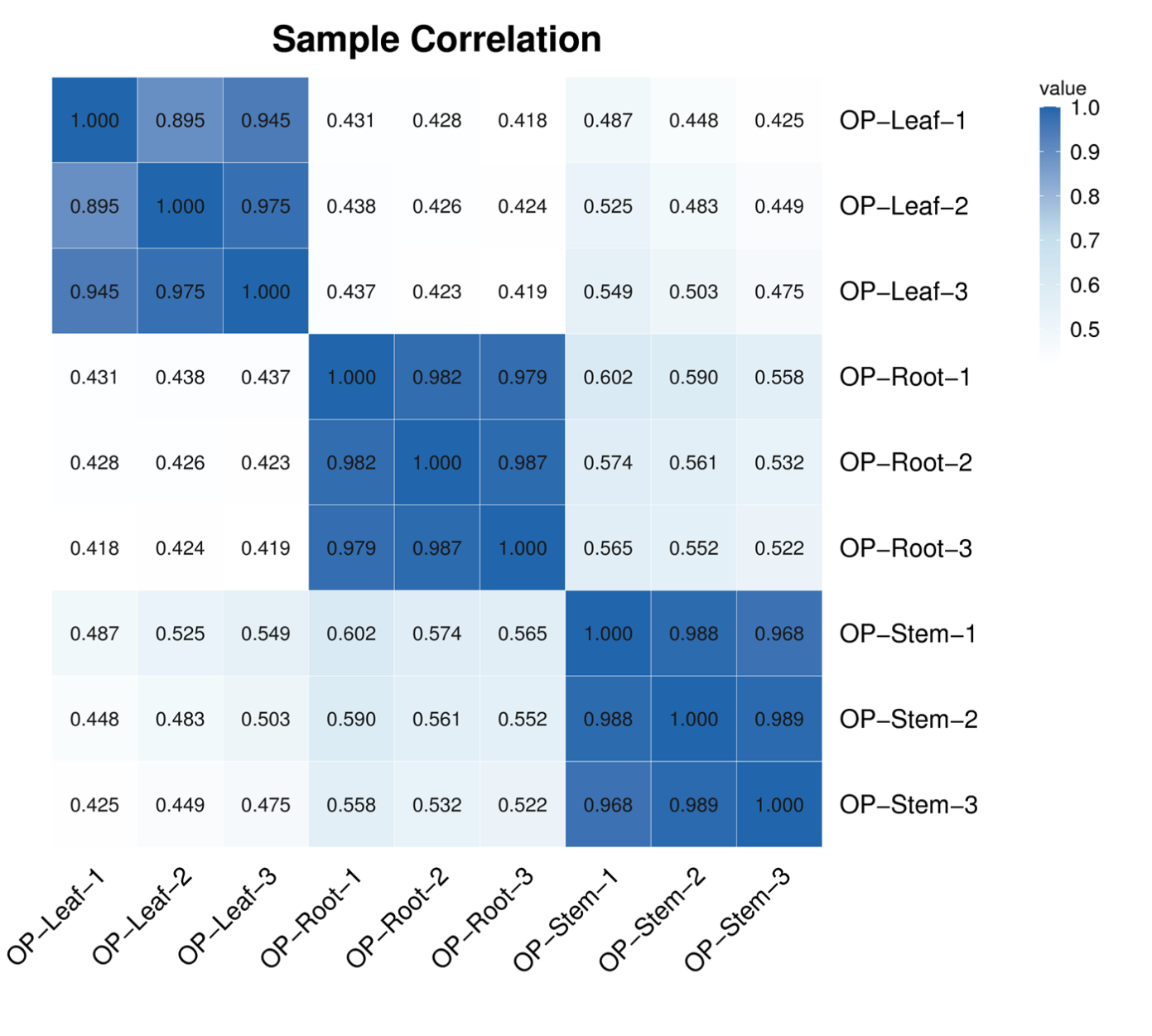

**Fig. S2. Correlation analysis of RNA-seq samples from *O. pumila* leaves, stems, and roots.** Pearson correlation coefficients were calculated for three biological replicates from each tissue. The color intensity indicates the strength of correlation between paired samples, as shown by the scale bar.

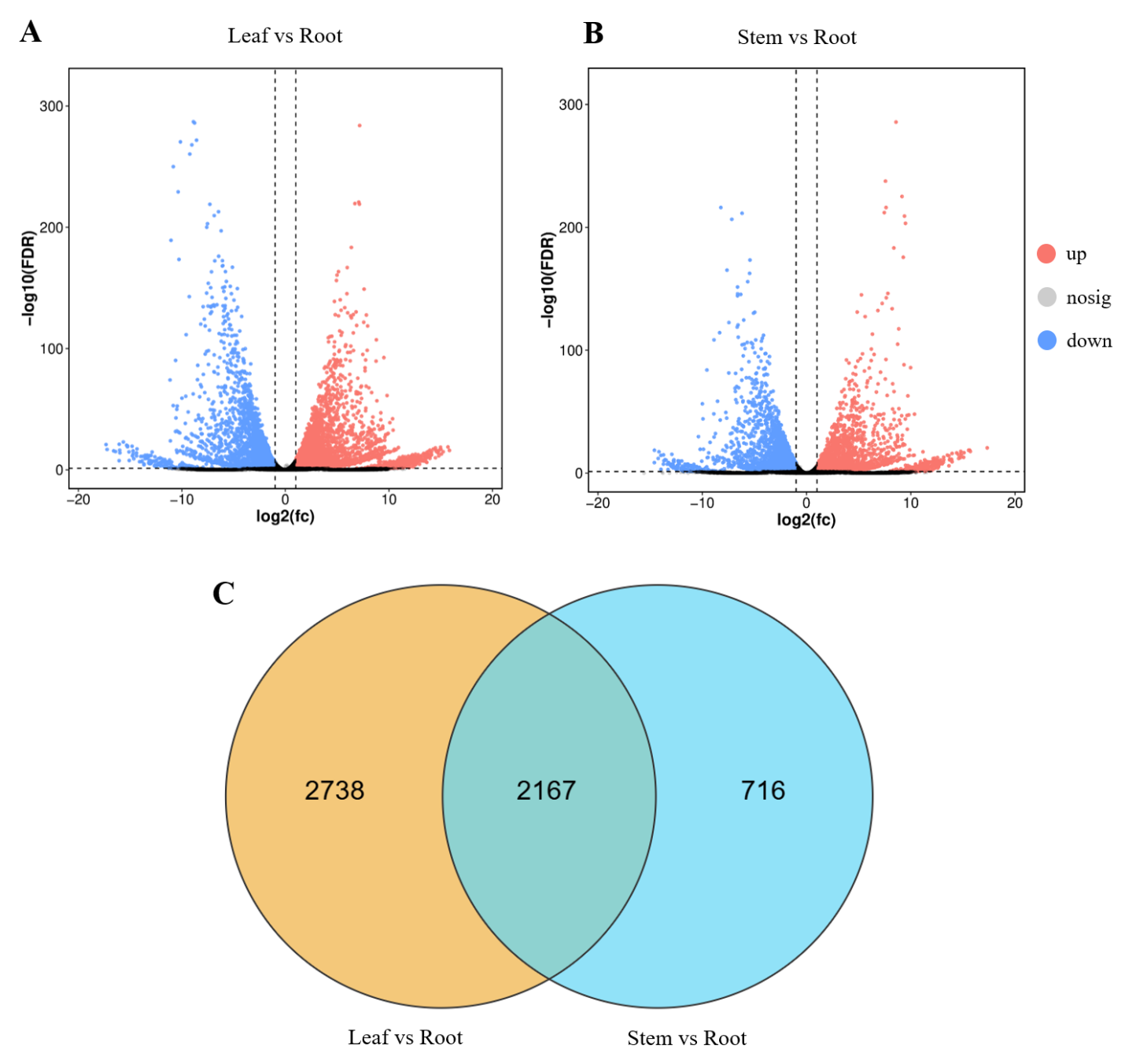

**Fig. S3. Difference analysis of genes in roots, stems and leaves from *O. pumila* plants*.* (A and B)** Volcano plots showing differential gene expression in root versus leaf **(A)** and root versus stem **(B)** comparisons. Red and blue points denote genes with higher and lower expression in roots, respectively; gray points denote genes without significant differential expression under the applied thresholds. **(C)** Venn diagram showing the overlap between genes highly expressed in roots relative to leaves and genes highly expressed in roots relative to stems. The overlap represents root-enriched genes shared by both comparisons.

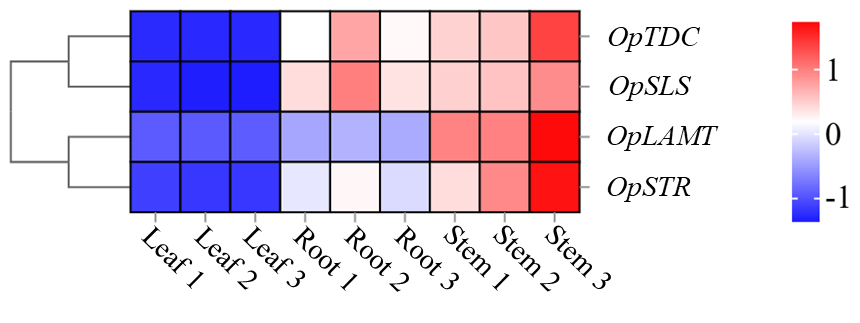

**Fig. S4. Expression patterns of known camptothecin biosynthetic genes in *O. pumila* tissues.** The heatmap shows TPM-normalized expression levels across leaf, root, and stem samples. Each tissue includes three biological replicates. The color scale indicates relative expression after normalization.

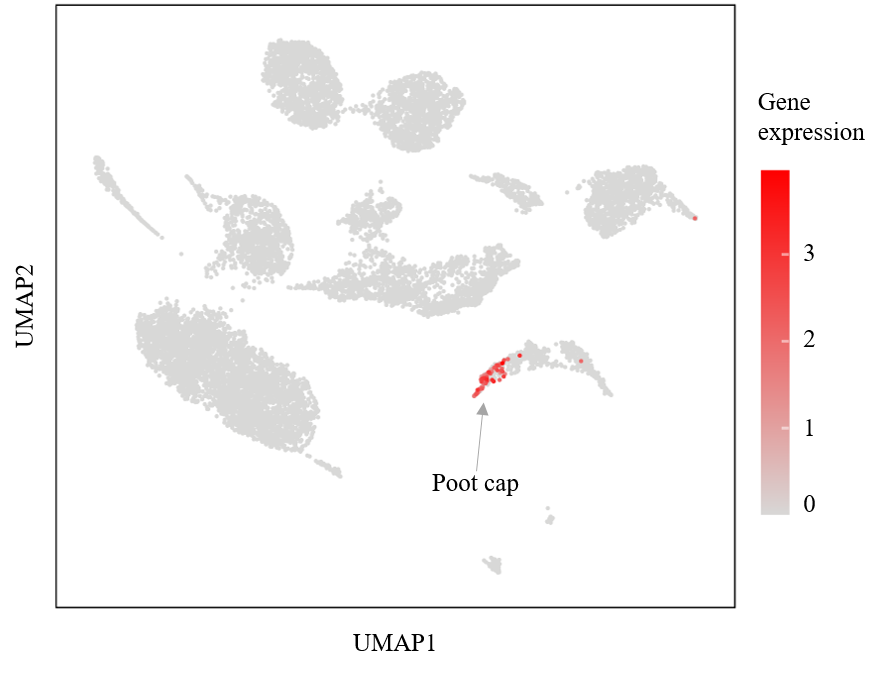

**Fig. S5. Expression profile of gene *Opuchr10-g0063950.1* at the single-cell level with the *O. pumila* hairy root.** The gene *Opuchr10-g0063950.1* was annotated as the marker gene of root cap according to homologous gene *AT1G33280* from *Arabidopsis thaliana*. The red dots represent the cells highly expressing *Opuchr10-g0063950.1*.

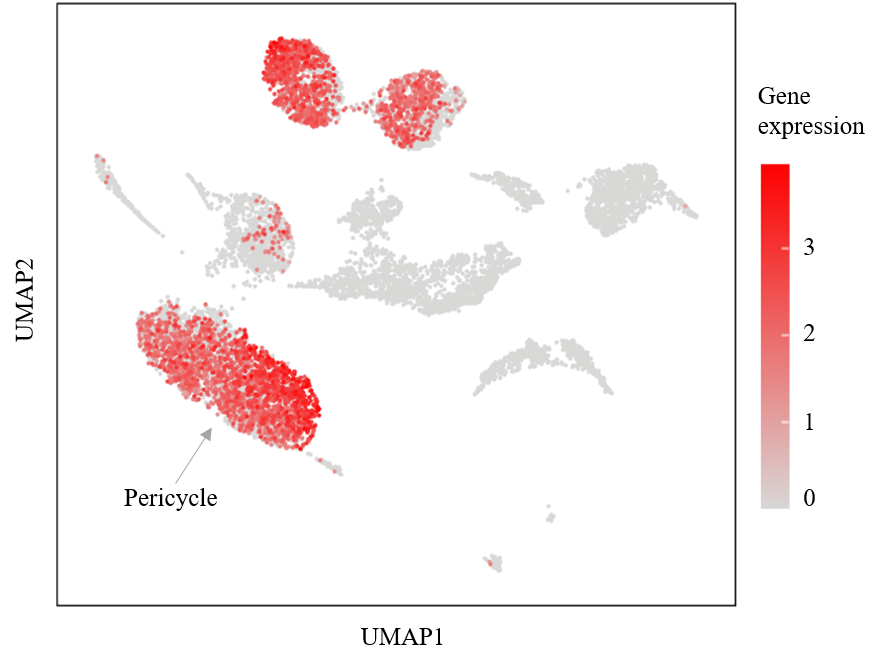

**Fig. S6. Expression profile of gene *Opuchr05-g0068550.1* at the single-cell level with the *O. pumila* hairy root.** The gene *Opuchr05-g0068550.1* was annotated as the marker gene of pericycle according to homologous gene *AT1G32450* from *A. thaliana*. The red dots represent the cells highly expressing *Opuchr05-g0068550.1*.

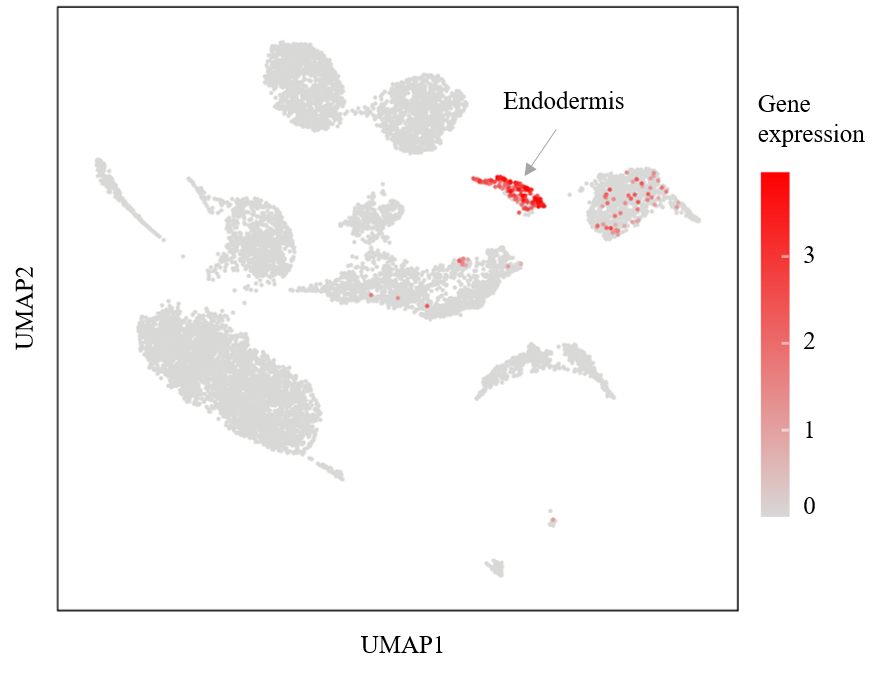

**Fig. S7. Expression profile of gene *Opuchr08-g0062080.1* at the single-cell level with the *O. pumila* hairy root.** The gene *Opuchr08-g0062080.1* was annotated as the marker gene of endodermis according to homologous gene *AT1G61590* from *A. thaliana*. The red dots represent the cells highly expressing *Opuchr08-g0062080.1*.

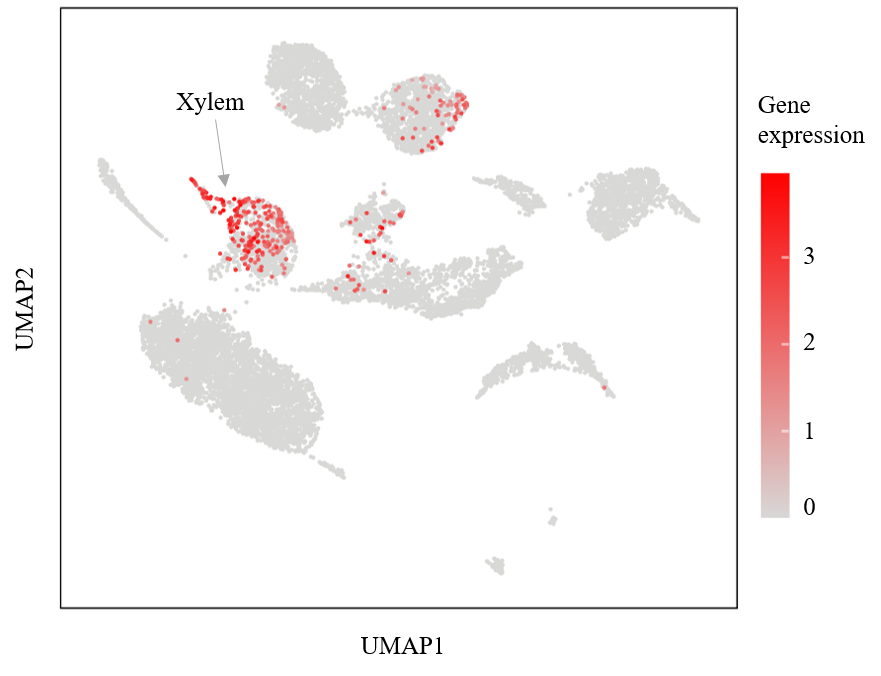

**Fig. S8. Expression profile of gene *Opuchr11-g0077910.1* at the single-cell level with the *O. pumila* hairy root.** The gene *Opuchr11-g0077910.1* was annotated as the marker gene of xylem according to homologous gene *AT1G68810* from *A. thaliana*. The red dots represent the cells highly expressing *Opuchr11-g0077910.1*.

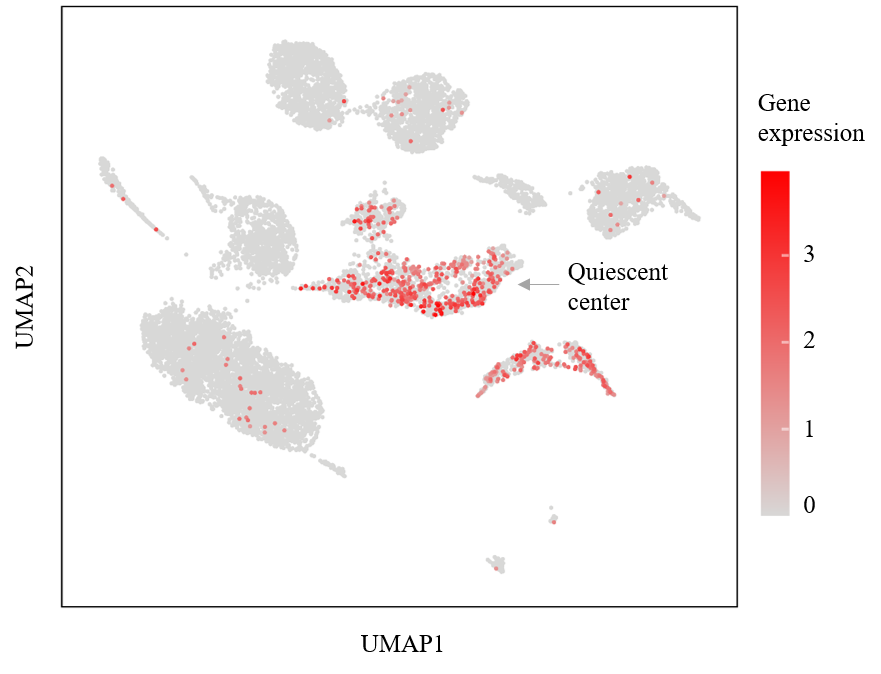

**Fig. S9. Expression profile of gene *Opuchr07-g0011840.1* at the single-cell level with the *O. pumila* hairy root.** The gene *Opuchr07-g0011840.1* was annotated as the marker gene of quiescent center according to homologous gene *AT3G20840* from *A. thaliana*. The red dots represent the cells highly expressing *Opuchr07-g0011840.1*.

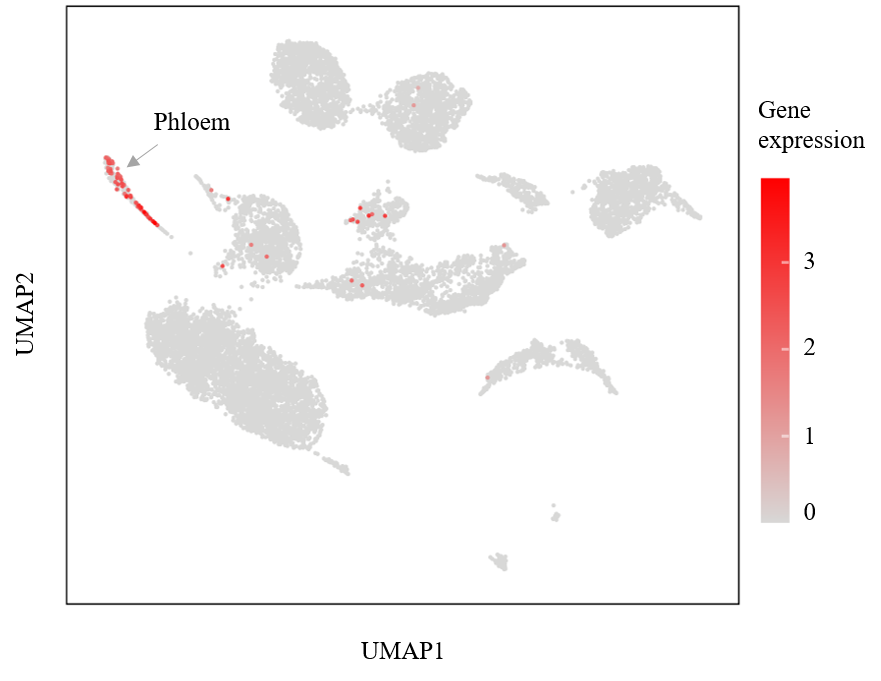

**Fig. S10. Expression profile of gene *Opuchr02-g0008300.1* at the single-cell level with the *O. pumila* hairy root.** The gene *Opuchr02-g0008300.1* was annotated as the marker gene of phloem according to homologous gene *AT1G79430* from *A. thaliana*. The red dots represent the cells highly expressing *Opuchr02-g0008300.1*.

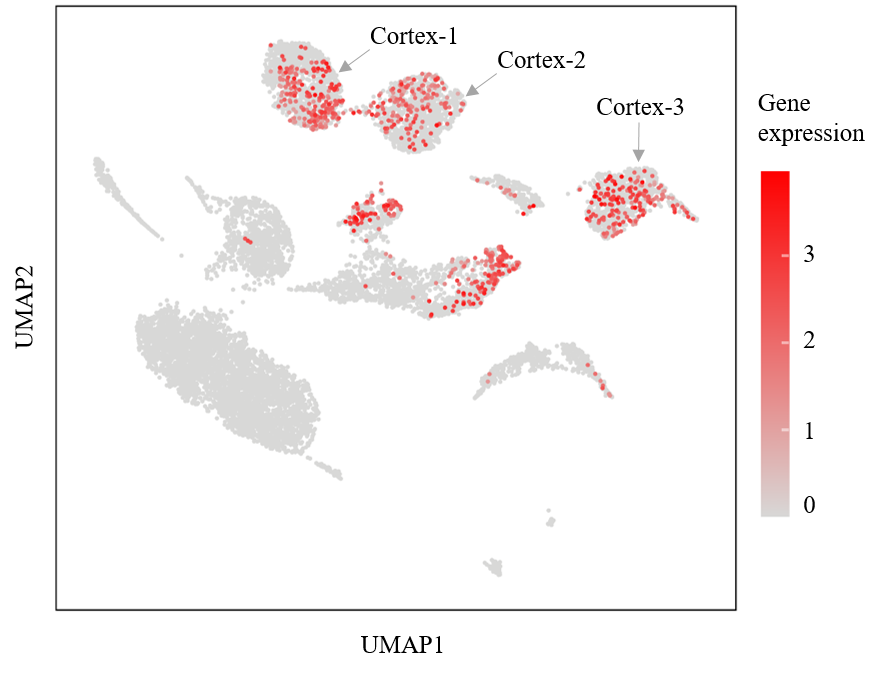

**Fig. S11. Expression profile of gene *Opuchr06-g0003570.1* at the single-cell level with the *O. pumila* hairy root.** The gene *Opuchr06-g0003570.1* was annotated as the marker gene of cortex according to homologous gene *AT5G53370* from *A. thaliana*. The red dots represent the cells highly expressing *Opuchr06-g0003570.1*.

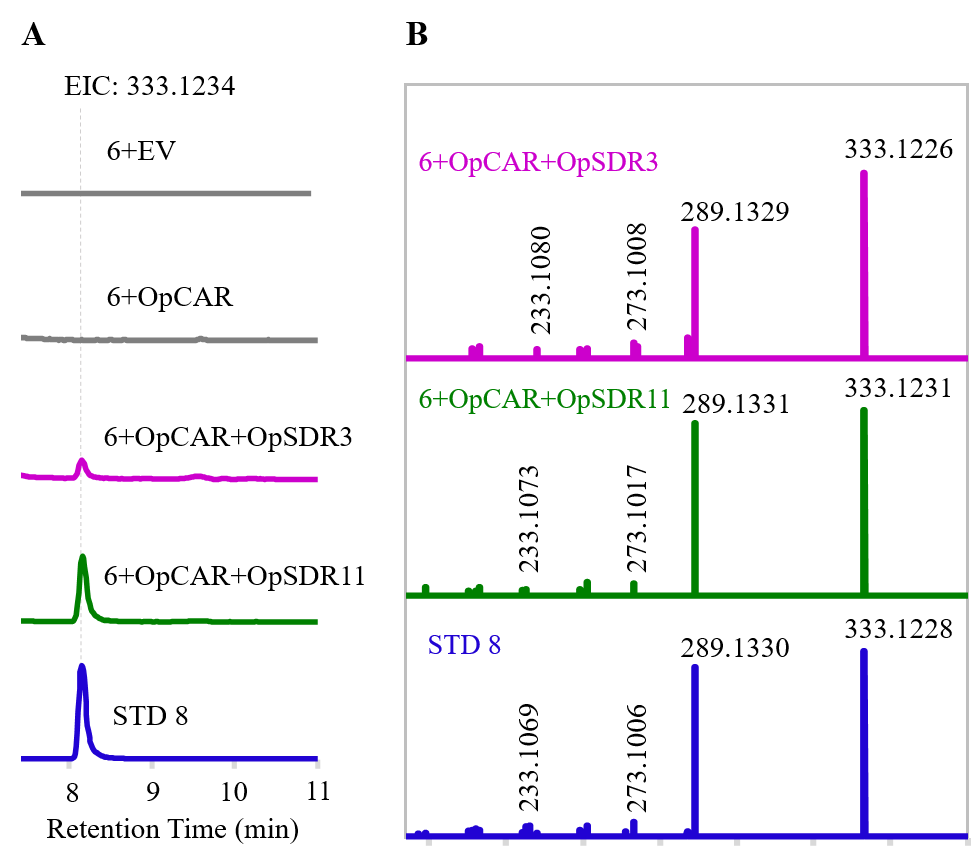

**Fig. S12. Functional characterization of the OpSDR and OpCAR.** (**A**). LC-MS analysis of the product deoxycamptothecin **8** in the reaction of OpSDR and OpCAR. The crude reaction product of OpCAR was mixed with OpSDR3 and OpSDR11, respectively. Extracted ion chromatogram (EIC) for deoxycamptothecin **8** (m/z [M+H]^+^ = 333.1234) produced in these reaction are shown. The reaction of EV and camptothecoside aglycone **6** was performed as the negative control. (**B**). MS/MS spectra of deoxycamptothecin **8** produced in the reaction of OpSDR3, OpSDR11, and OpCAR. The main fragments of product deoxycamptothecin **8** were compared with those of standard deoxycamptothecin **8**. STD, standard. EV, empty vector.

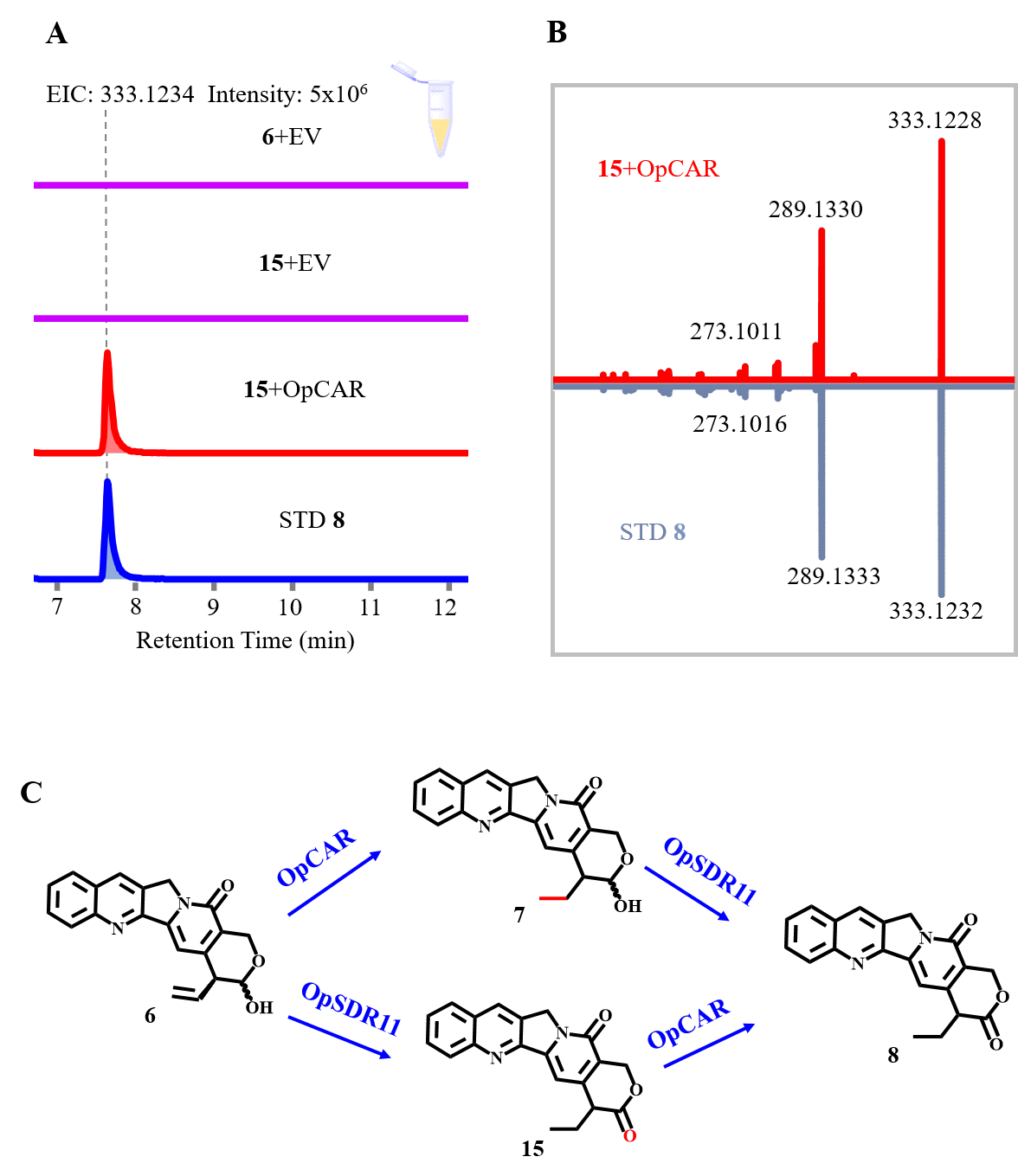

**Fig. S13. Functional characterization of the reaction of OpSDR11 and compound 15. (A)** LC-MS analysis of the product **8** in the reaction of crude **15** and pure OpCAR obtained by *E. coli*. OpSDR11 was used to catalyze camptothecoside aglycone **6** to obtain crude product **15** which was then mixed with OpCAR. Extracted ion chromatogram for deoxycamptothecin **8** (m/z [M+H]^+^ = 333.1234) are shown. **(B)** MS/MS spectra of deoxycamptothecin **8** produced by the reaction of pure protein OpSDR11 and **15**; the fragments are compared with the standard deoxycamptothecin **8**. (**C**) The biosynthetic pathway of deoxycamptothecin **8** from camptothecoside aglycone **6**.

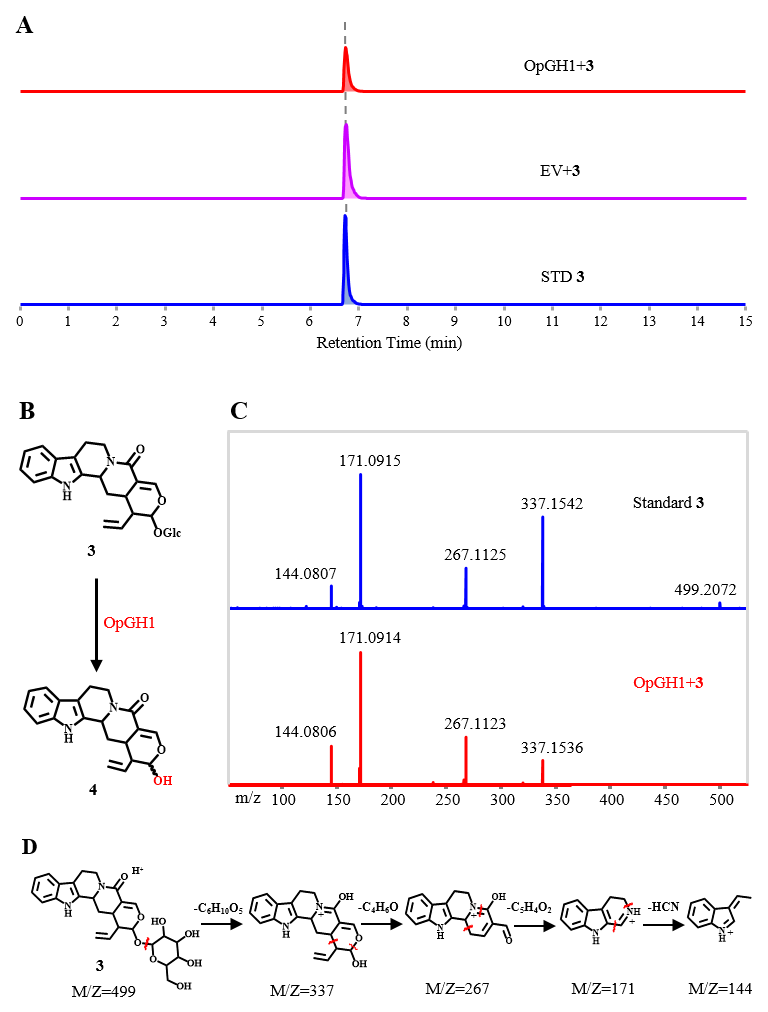

**Fig. S14. Characterization of reaction of OpGH1 and strictosamide 3.** (**A**). LC-MS analysis of substrate strictosamide **3** in the reaction of OpGH1. (**B**). Reaction catalyzed by OpGH1. (**C**). MS/MS spectra of product strictosamide aglycone **4** produced by OpGH1. (**D**). Cleavage pattern for product strictosamide aglycone **4**.

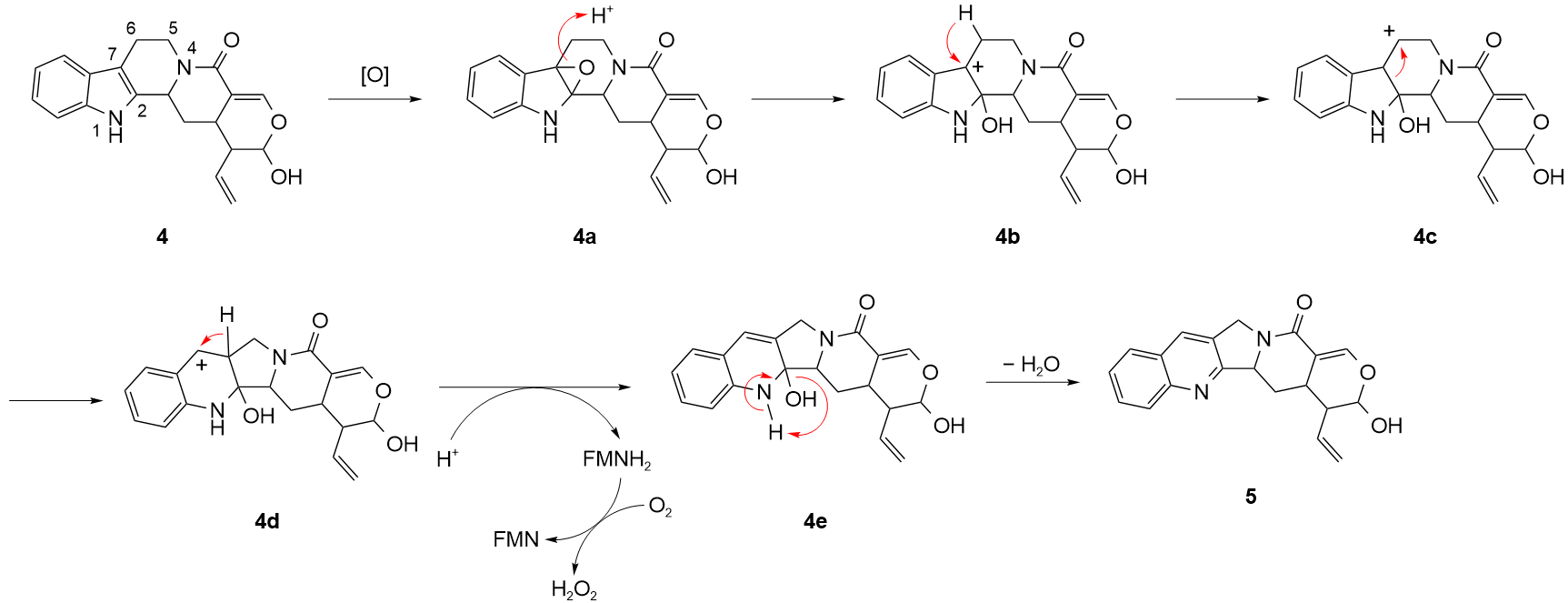
 **Fig. S15. Proposed mechanism for the FMN‑mediated conversion of strictosamide aglycone 4 to deoxypumiloside aglycone 5.** Based on the experimental observations, a detailed mechanism for the conversion of strictosamide aglycone **4** to the deoxypumiloside aglycone **5** is proposed. This transformation proceeds through a cascade of oxidative and rearrangement events, with FMN playing a pivotal role as a mediator of deprotonation. The reaction is initiated by the spontaneous oxidation of C2–C7 double bond of **4** under aerobic conditions to afford the corresponding epoxide **4a**. Subsequent protonation of the epoxide oxygen facilitates ring opening, generating a tertiary carbocation at C7 **4b**, which is stabilized by the adjacent π‑system of the indole moiety. The C7 carbocation **4b** then undergoes a 1, 2‑hydride shift from C6 to C7, relocating the positive charge to C6 and furnishing cation **4c**. The C6 cation **4c** then undergoes a cation‑driven Wagner–Meerwein‑type rearrangement, wherein the C2–C7 *σ*‑bond is cleaved and a new C2–C6 bond is concomitantly formed. This deep‑seated rearrangement converts the original 6–5–6–6–6 fused ring system into a 6–6–5–6–6 arrangement, generating a new carbocation at C7 **4d**. The C7 cation **4d** is converted to the corresponding alkene **4e** by elimination of the proton at C6. This deprotonation is facilitated by oxidized FMN, which abstracts a hydride equivalent from C6, yielding the C6–C7 double bond and regenerating FMNH₂. FMN acts as a two‑electron oxidant and is reduced to FMNH₂, which is subsequently reoxidised by molecular oxygen to reform the catalytically active flavin with concomitant production of H₂O₂. Finally, spontaneous intramolecular dehydration between the NH-1 and the hydroxyl group at C2 closes the pyridine ring, affording the pentacyclic deoxypumiloside aglycone **5**.

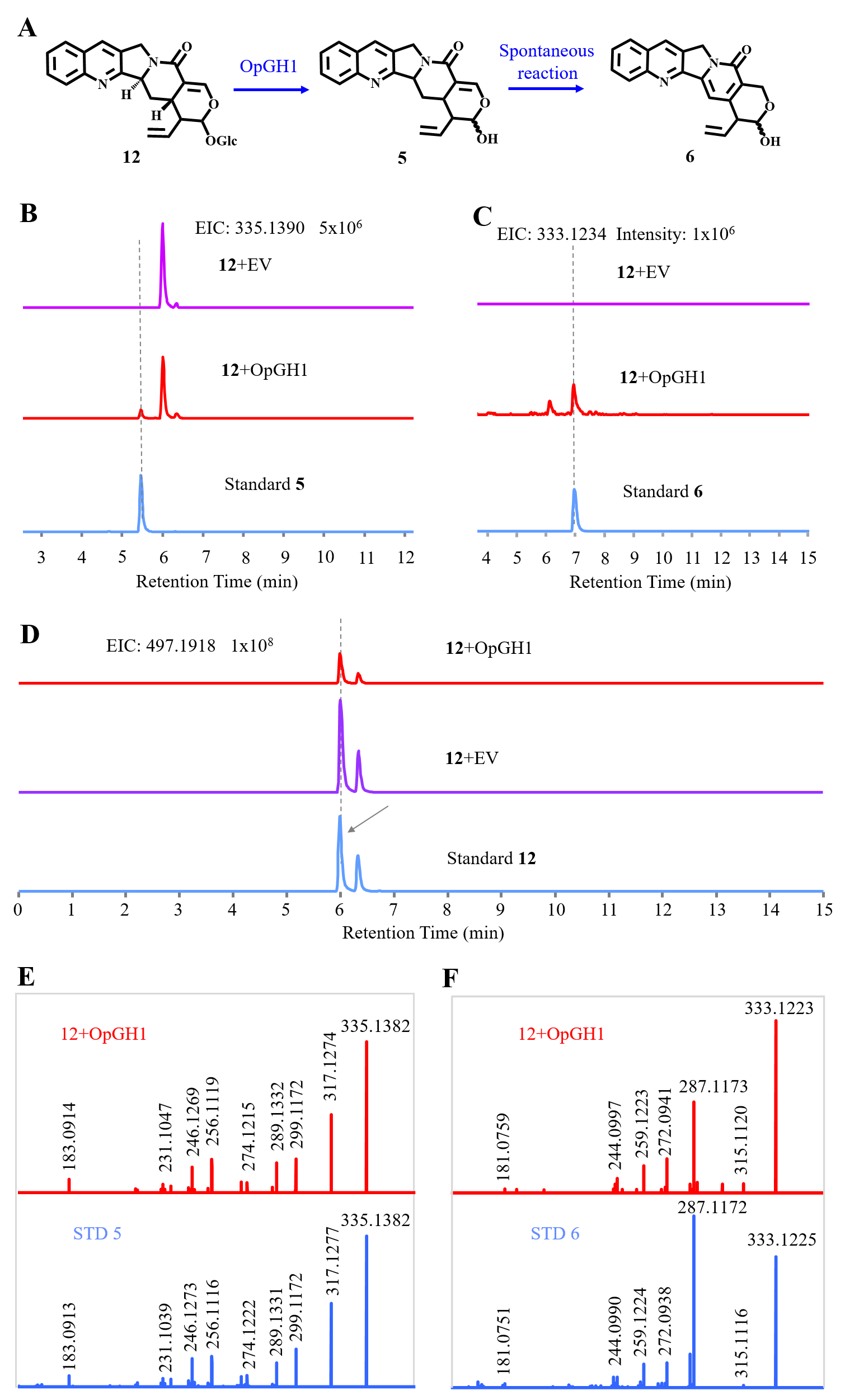

**Fig. S16. Characterization of catalytic activity of OpGH1 on (3*S*)-deoxypumiloside 12.** (**A**). Reaction catalyzed by OpGH1. (**B**). LC-MS analysis of the deoxypumiloside aglycone **5** in the reaction of OpGH1 and (3*S*)-deoxypumiloside **12**, extracted ion chromatogram for product deoxypumiloside aglycone **5** (m/z [M+H]^+^ = 335.1390) are shown. (**C**). LC-MS analysis of the camptothecoside aglycone **6** in the reaction of OpGH1 and (3*S*)-deoxypumiloside **12**, extracted ion chromatogram for product camptothecoside aglycone **6** (m/z [M+H]^+^ = 333.1234) are shown. (**D**). LC-MS analysis of substrate (3*S*)-deoxypumiloside **12** in the reaction of OpGH1. (**E**). MS/MS spectra of deoxypumiloside aglycone **5** produced by the reaction of OpGH1 and (3*S*)-deoxypumiloside **12**. The main fragments of product deoxypumiloside aglycone **5** were compared with those of standard deoxypumiloside aglycone **5**. (**F**). MS/MS spectra of camptothecoside aglycone **6** produced in the reaction of OpGH1 and (3*S*)-deoxypumiloside **12**. EV, empty vector.

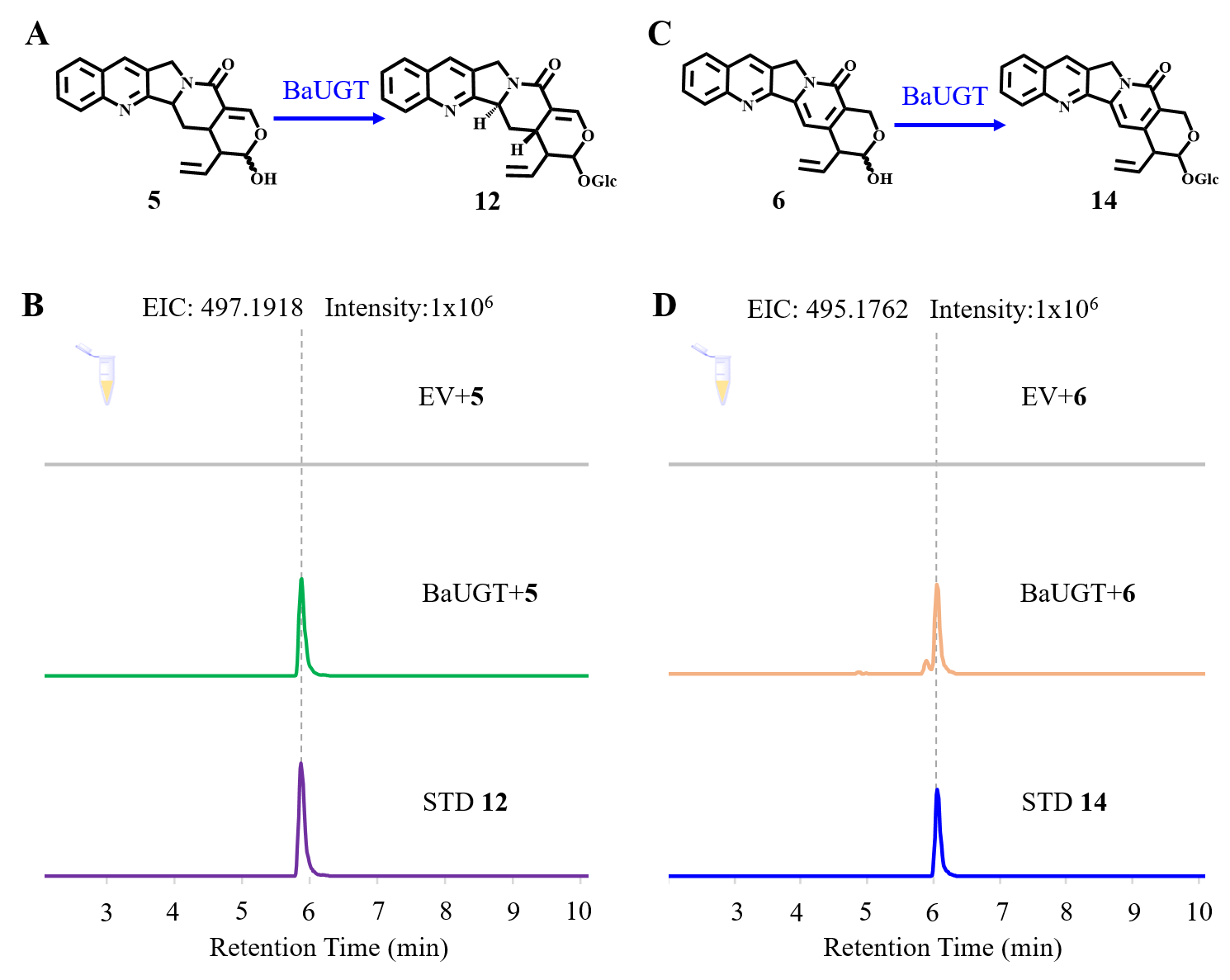

**Fig. S17. Functional characterization of BaUGT in vitro.** (**A**). Reaction catalyzed by BaUGT in vitro. (**B**). LC-MS analysis of the product **12** in the reaction of BaUGT; extracted ion chromatogram for (3*S*)-deoxypumiloside **12** (m/z [M+H]^+^ = 497.1918) are shown. (**C**). Biosynthesis of camptothecoside **14** catalyzed by BaUGT in vitro. (**D**). LC-MS analysis of the product **14** in the reaction of BaUGT; extracted ion chromatogram for camptothecoside **14** (m/z [M+H]^+^ = 495.1762) are shown.

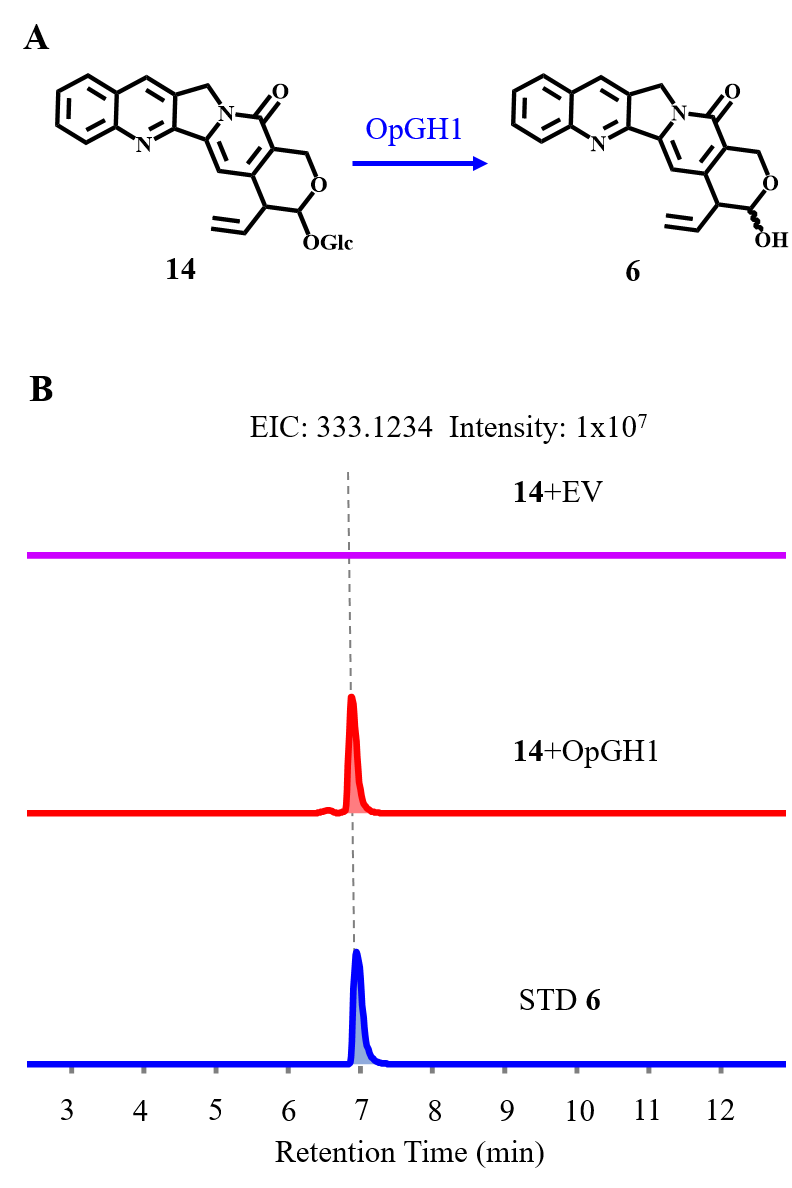

**Fig. S18. Functional characterization of the reaction of OpGH1 and** **camptothecoside 14.** (**A**). Reaction catalyzed by OpGH1 in vitro. (**B**). LC-MS analysis of the camptothecoside aglycone **6** in the reaction of OpGH1 and camptothecoside **14**, extracted ion chromatogram for product camptothecoside aglycone **6** (m/z [M+H]^+^ = 333.1234) are shown. STD, standard. EV, empty vector.

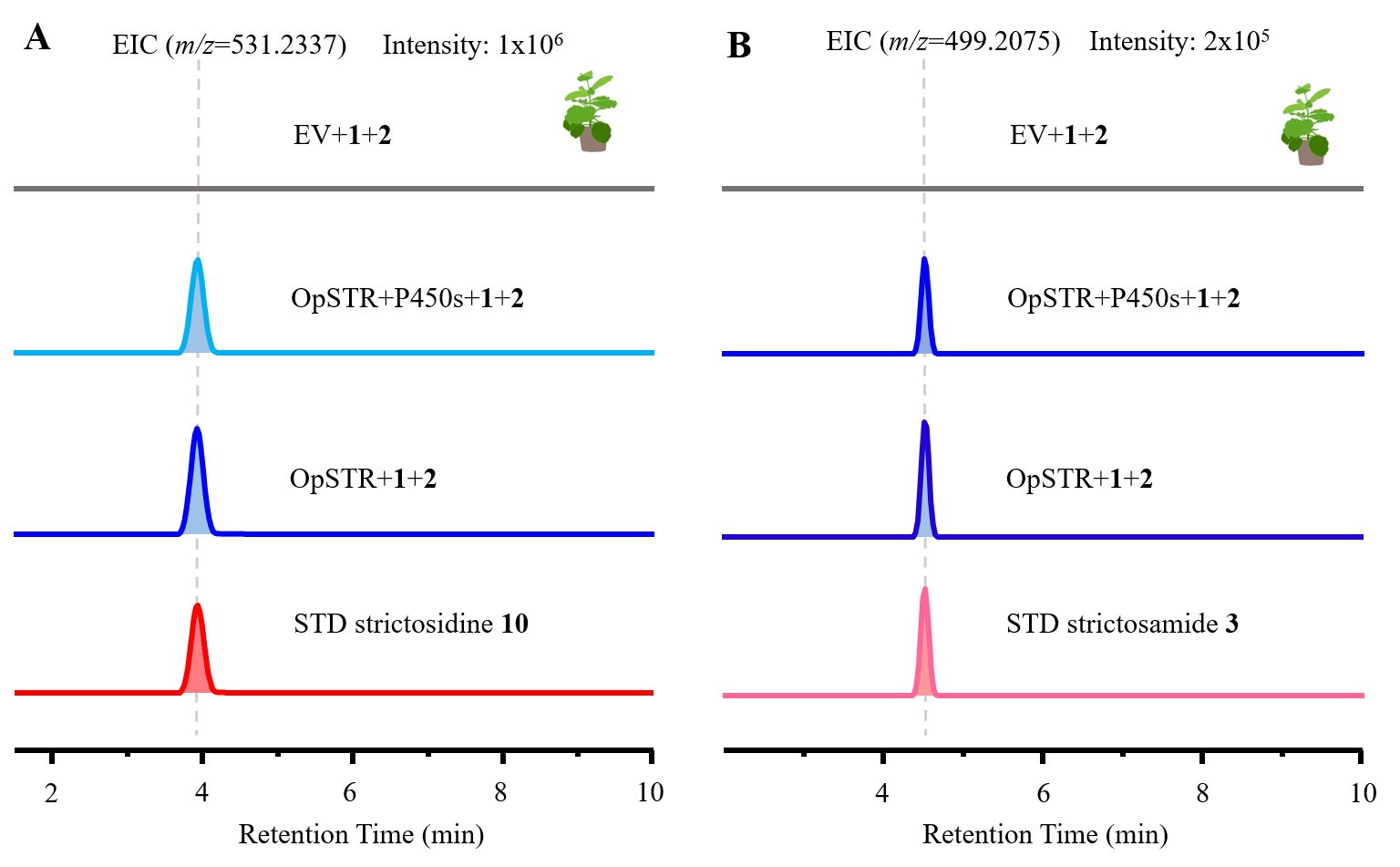

**Fig. S19. Functional characterization of OpSTR and cytochrome P450s** **in the *N. benthamiana*.** (**A**). Extracted ion chromatogram for strictosidine **10** (m/z [M+H]^+^ = 531.2337) produced in the reaction of single OpSTR, mixture cytochrome P450s and OpSTR. (**B**). Extracted ion chromatogram for strictosamide **3** (m/z [M+H]^+^ = 499.2075) produced in the reaction of single OpSTR, mixture containing cytochrome P450s and OpSTR. EV, empty vector; STD, standard.

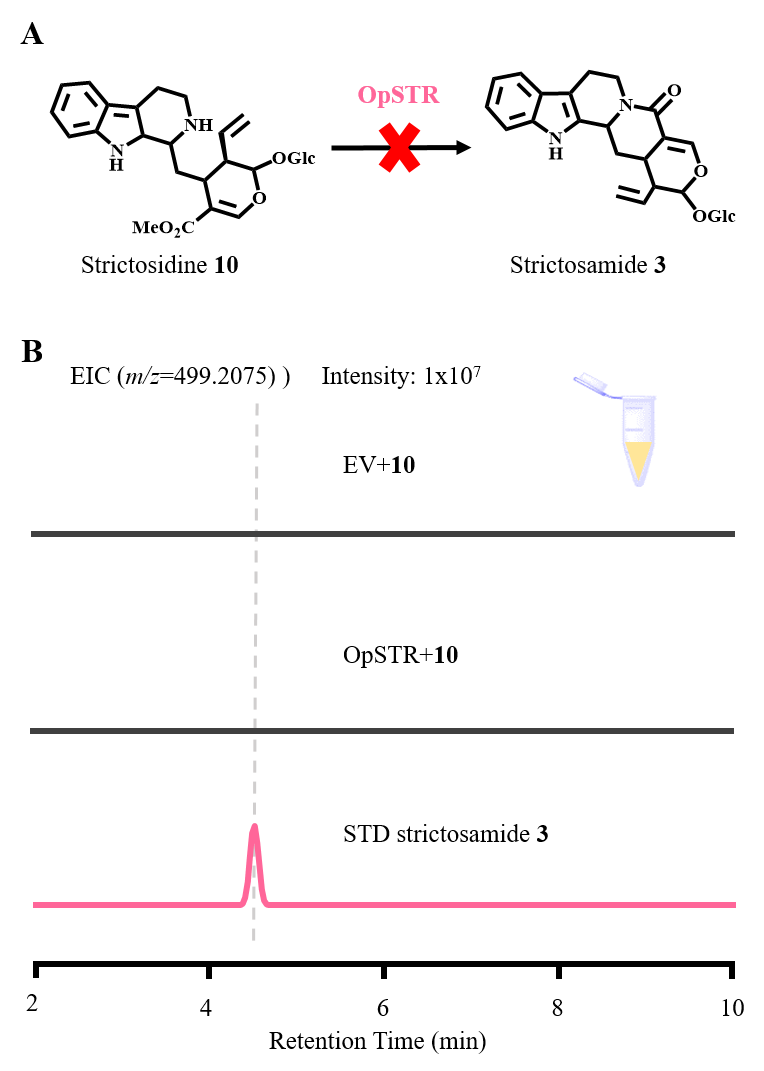

**Fig. S20. Characterization of the OpSTR reaction with strictosidine 10 in vitro.** (**A**). Proposed reaction catalyzed by OpSTR. (**B**). Extracted ion chromatogram for strictosamide **3** (m/z [M+H]^+^ = 499.2075) produced in the reaction of OpSTR and strictosidine **10**. Corresponding extracted ion chromatogram of strictosamide **3** could not be discovered, indicating OpSTR could not catalyze strictosidine **10** to yield strictosamide **3**. EV, empty vector; STD, standard.

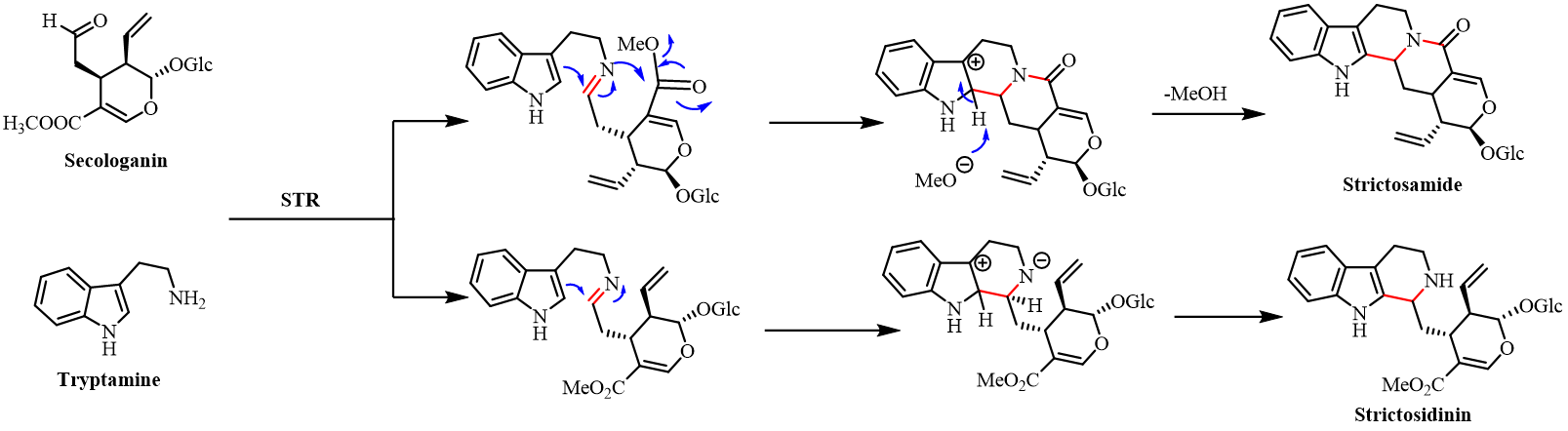

**Fig. S21. Proposed mechanism of OpSTR reaction produing stictosamide 3 and strictosidine 10.** The protein OpSTR may catalyze the formation of different transition states from the substrates tryptamine 1 and secologanin 2, thereby generating two products.

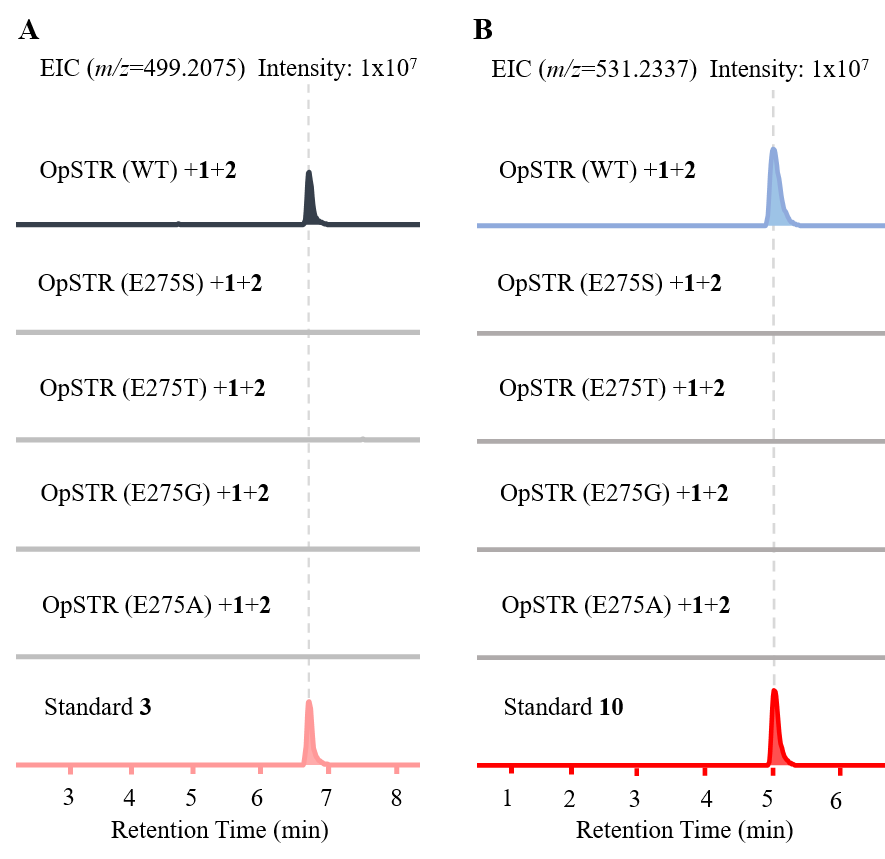

**Fig. S22. Functional characterization of the key amino acid 275E of OpSTR.** (**A**). LC-MS analysis of strictosamide **3** produced in the reaction of OpSTR mutants, extracted ion chromatogram for product strictosamide **3** (m/z [M+H]^+^ = 499.2075) are shown. (**B**). LC-MS analysis of strictosidine **10** produced in the reaction of OpSTR mutants, extracted ion chromatogram for product strictosidine **10** (m/z [M+H]^+^ = 531.2337) are shown. STD, standard. WT, wild type.

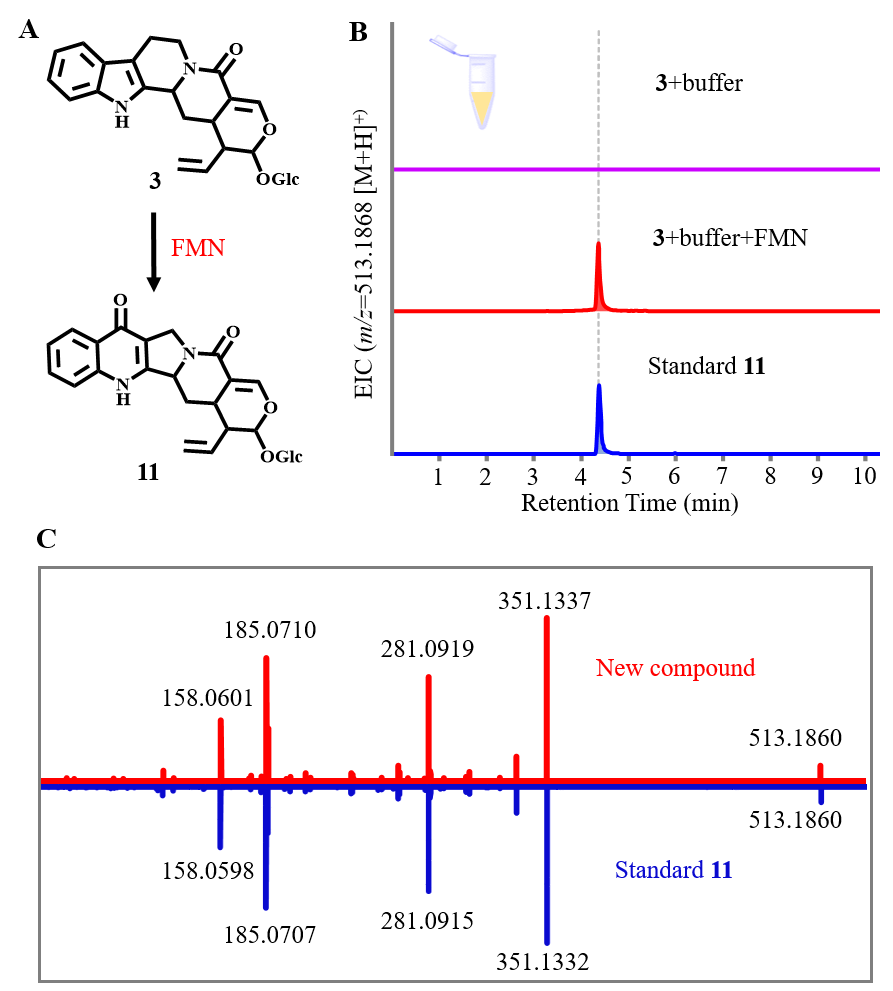

**Fig. S23. Reaction catalyzed by FMN in vitro.** (**A**). FMN catalyze strictosamide **3** to produce pumiloside **11**. (**B**). Extracted ion chromatogram for product pumiloside **11** (m/z [M+H]^+^ = 513.1868) in the reaction of strictosamide **3** and FMN. (**C**). MS/MS spectra of pumiloside **11** produced by FMN. FMN, flavin mononucleotide.

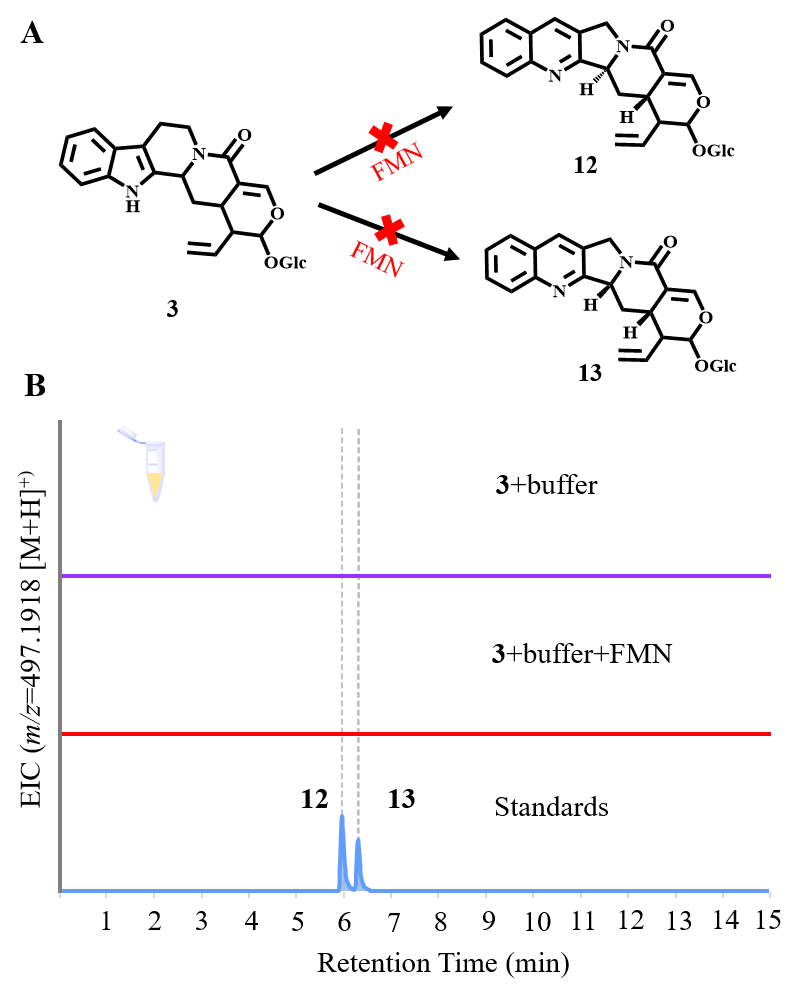

**Fig. S24. Characterization of the product in the reaction of FMN and strictosamide** **3.** (**A**). Proposed biosynthetic pathway from strictosamide **3** to (3*S*)-deoxypumiloside **12** and (3*R*)-deoxypumiloside **13**. (**B**). LC-MS analysis of the (3*S*)-deoxypumiloside **12** and (3*R*)-deoxypumiloside **13** in the reaction of FMN and strictosamide **3,** extracted ion chromatogram for product (3*S*)-deoxypumiloside **12** and (3*R*)-deoxypumiloside **13** (m/z [M+H]^+^ = 497.1918) could not be discovered. FMN, flavin mononucleotide.

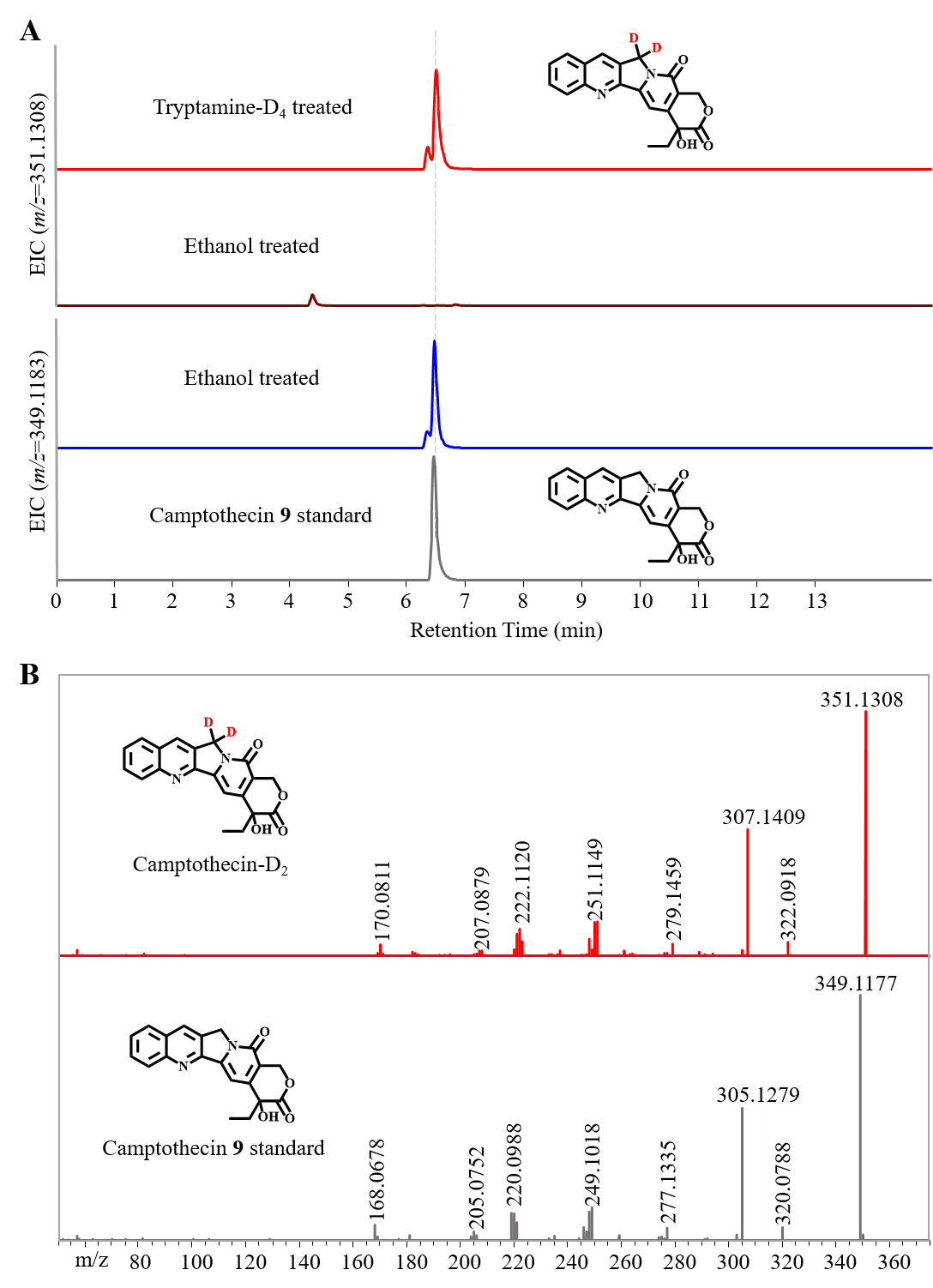

**Fig. S25. LC-MS analysis of camptothecin 9 in the isotope feeding experiment.** (**A**). Extracted ion chromatogram for deuterium-labeled camptothecin-D_2_ (m/z [M+H]^+^ = 351.1308) and camptothecin **9** (m/z [M+H]^+^ = 349.1183) are shown. Tryptamine-D_4_ dissolved in the ethanol was added into the *O. pumila* hairy roots to label the intermediates involved in the camptothecin **9** biosynthesis. Ethanol was solely added into the *O. pumila* hairy roots as the negative control. (**B**). MS/MS spectra of deuterium-labeled camptothecin-D_2_ and standard camptothecin **9**. The main fragments of deuterium-labeled camptothecin-D_2_ were compared with those of standard camptothecin **9**.

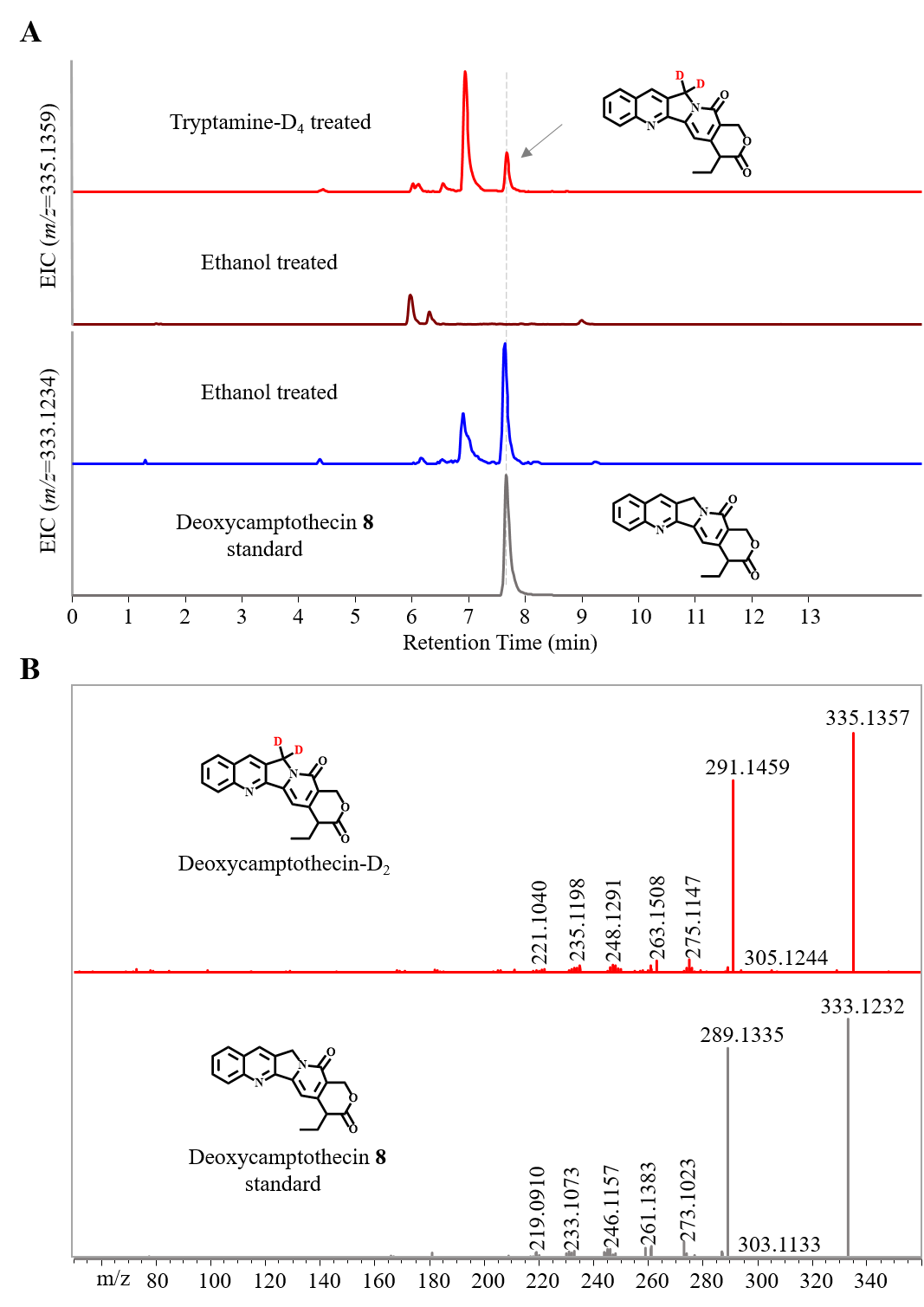

**Fig. S26. LC-MS analysis of deoxycamptothecin 8 in the isotope feeding experiment.** (**A**). Extracted ion chromatogram for deuterium-labeled deoxycamptothecin-D_2_ (m/z [M+H]^+^ = 335.1359) and deoxycamptothecin **8** (m/z [M+H]^+^ = 333.1234) are shown. Tryptamine-D_4_ dissolved in the ethanol was added into the *O. pumila* hairy roots to label the intermediates involved in the camptothecin **9** biosynthesis. Ethanol was solely added into the *O. pumila* hairy roots as the negative control. (**B**). MS/MS spectra of deuterium-labeled deoxycamptothecin-D_2_ and standard deoxycamptothecin **8**. The main fragments of deuterium-labeled deoxycamptothecin-D_2_ were compared with those of standard deoxycamptothecin **8**.

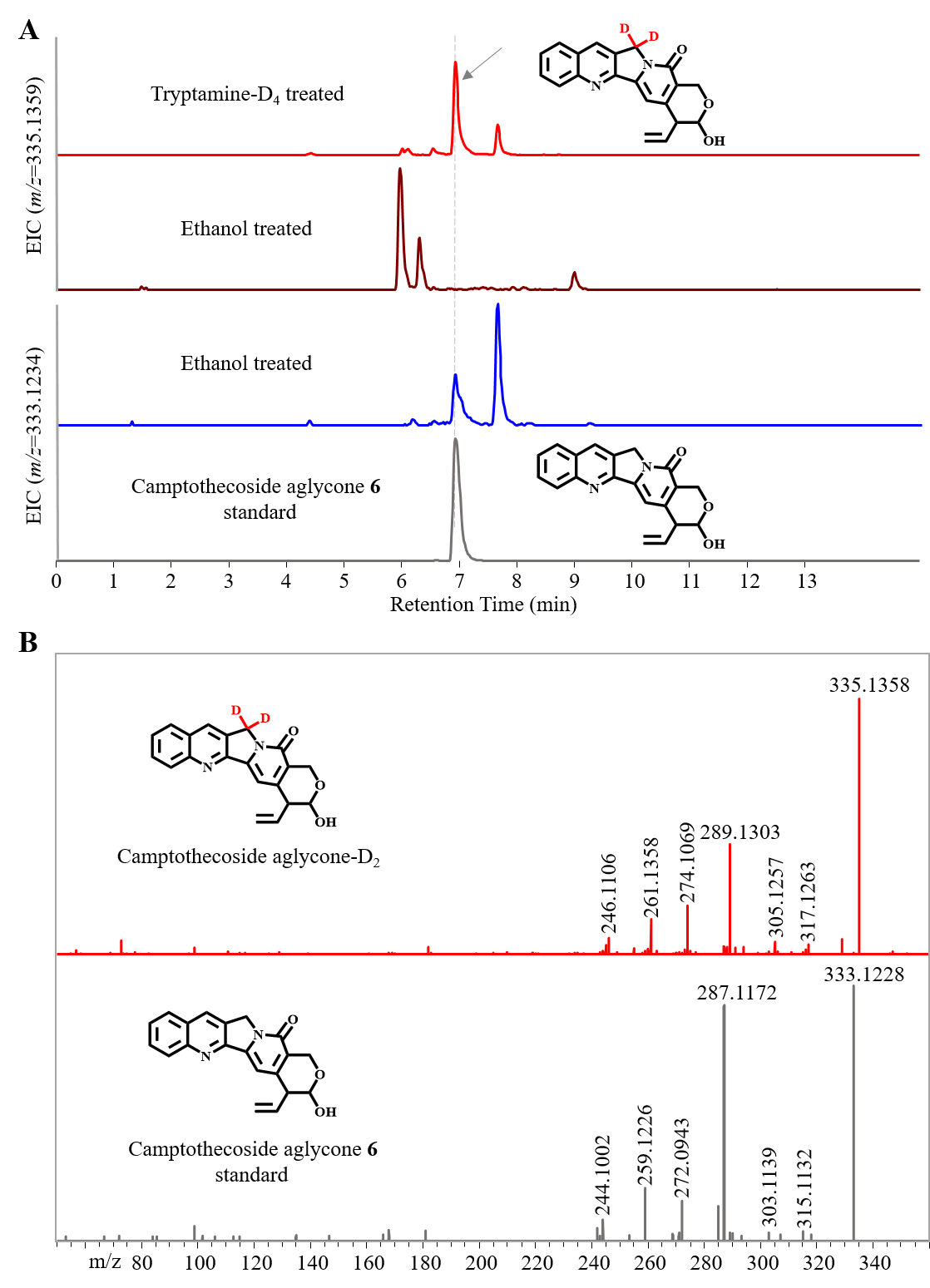

**Fig. S27. LC-MS analysis of camptothecoside aglycone 6 in the isotope feeding experiment.** (**A**). Extracted ion chromatogram for deuterium-labeled camptothecoside aglycone-D_2_ (m/z [M+H]^+^ = 335.1359) and camptothecoside aglycone **6** (m/z [M+H]^+^ = 333.1234) are shown. Tryptamine-D_4_ dissolved in the ethanol was added into the *O. pumila* hairy roots to label the intermediates involved in the camptothecin **9** biosynthesis. Ethanol was solely added into the *O. pumila* hairy roots as the negative control. (**B**). MS/MS spectra of deuterium-labeled camptothecoside aglycone-D_2_ and standard camptothecoside aglycone **6**. The main fragments of deuterium-labeled camptothecoside aglycone-D_2_ were compared with those of standard camptothecoside aglycone **6**.

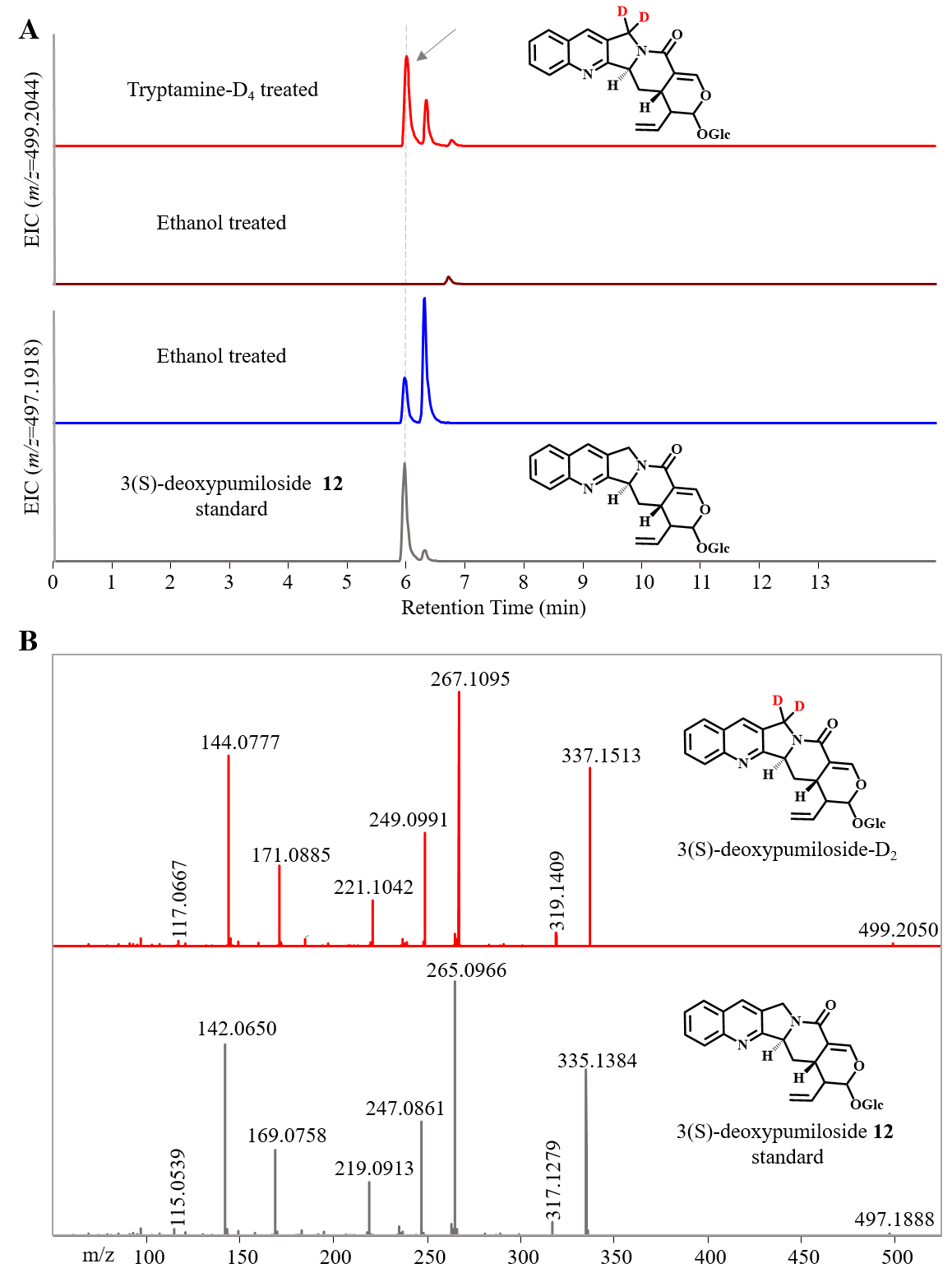

**Fig. S28. LC-MS analysis of 3(*S*)-deoxypumiloside 12 in the isotope feeding experiment.** (**A**). Extracted ion chromatogram for deuterium-labeled 3(*S*)-deoxypumiloside-D_2_ (m/z [M+H]^+^ = 499.2044) and 3(*S*)-deoxypumiloside **12** (m/z [M+H]^+^ = 497.1918) are shown. Tryptamine-D_4_ dissolved in the ethanol was added into the *O. pumila* hairy roots to label the intermediates involved in the camptothecin **9** biosynthesis. Ethanol was solely added into the *O. pumila* hairy roots as the negative control. (**B**). MS/MS spectra of deuterium-labeled 3(*S*)-deoxypumiloside-D_2_ and standard 3(*S*)-deoxypumiloside **12**. The main fragments of deuterium-labeled 3(*S*)-deoxypumiloside-D_2_ were compared with those of standard 3(*S*)-deoxypumiloside **12**.

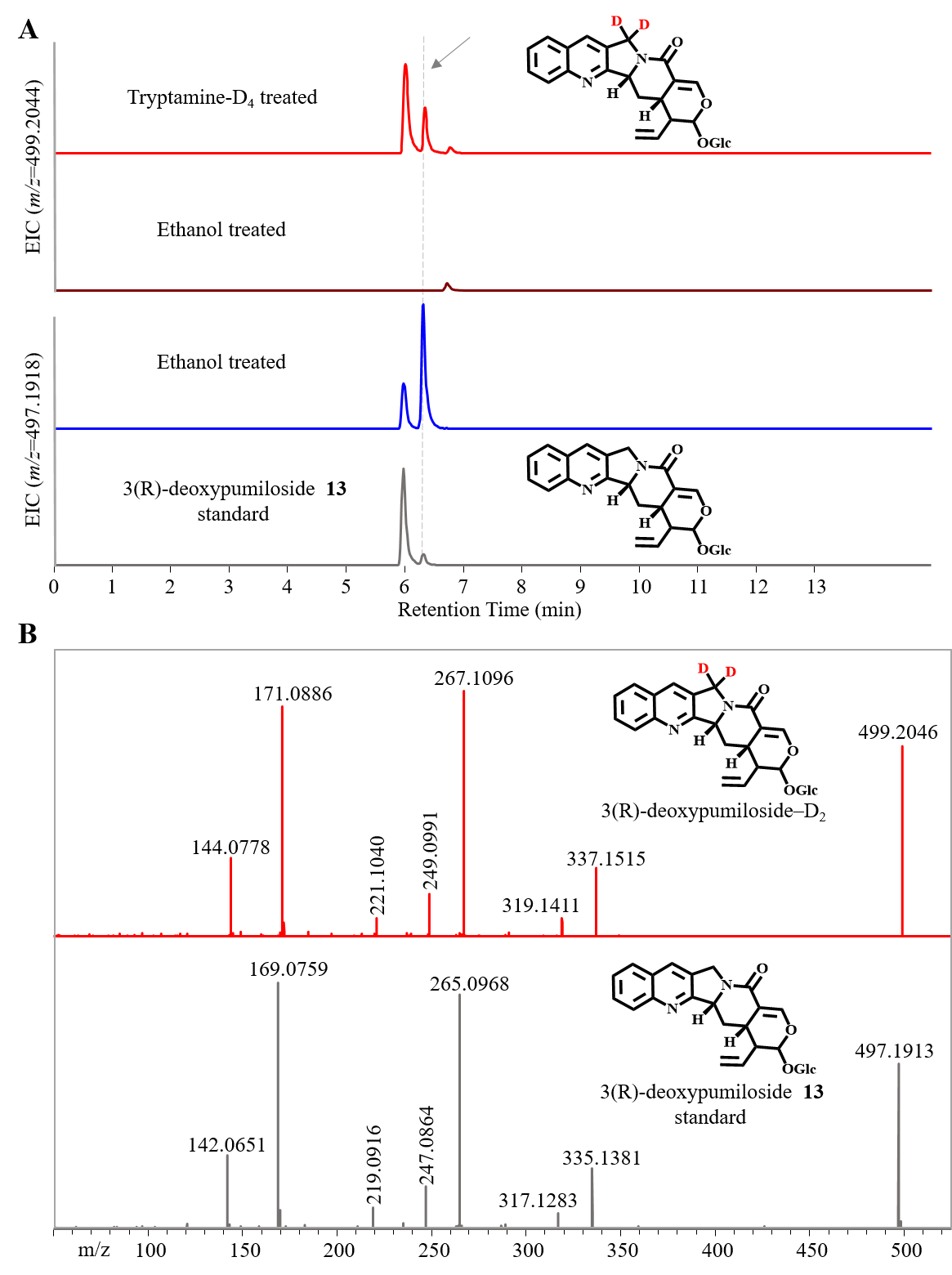

**Fig. S29. LC-MS analysis of 3(*R*)-deoxypumiloside 13 in the isotope feeding experiment.** (**A**). Extracted ion chromatogram for deuterium-labeled 3(*R*)-deoxypumiloside-D_2_ (m/z [M+H]^+^ = 499.2044) and 3(*R*)-deoxypumiloside **13** (m/z [M+H]^+^ = 497.1918) are shown. Tryptamine-D_4_ dissolved in the ethanol was added into the *O. pumila* hairy roots to label the intermediates involved in the camptothecin **9** biosynthesis. Ethanol was solely added into the *O. pumila* hairy roots as the negative control. (**B**). MS/MS spectra of deuterium-labeled 3(*R*)-deoxypumiloside-D_2_ and standard 3(*R*)-deoxypumiloside **13**. The main fragments of deuterium-labeled 3(R)-deoxypumiloside-D_2_ were compared with those of standard **13**.

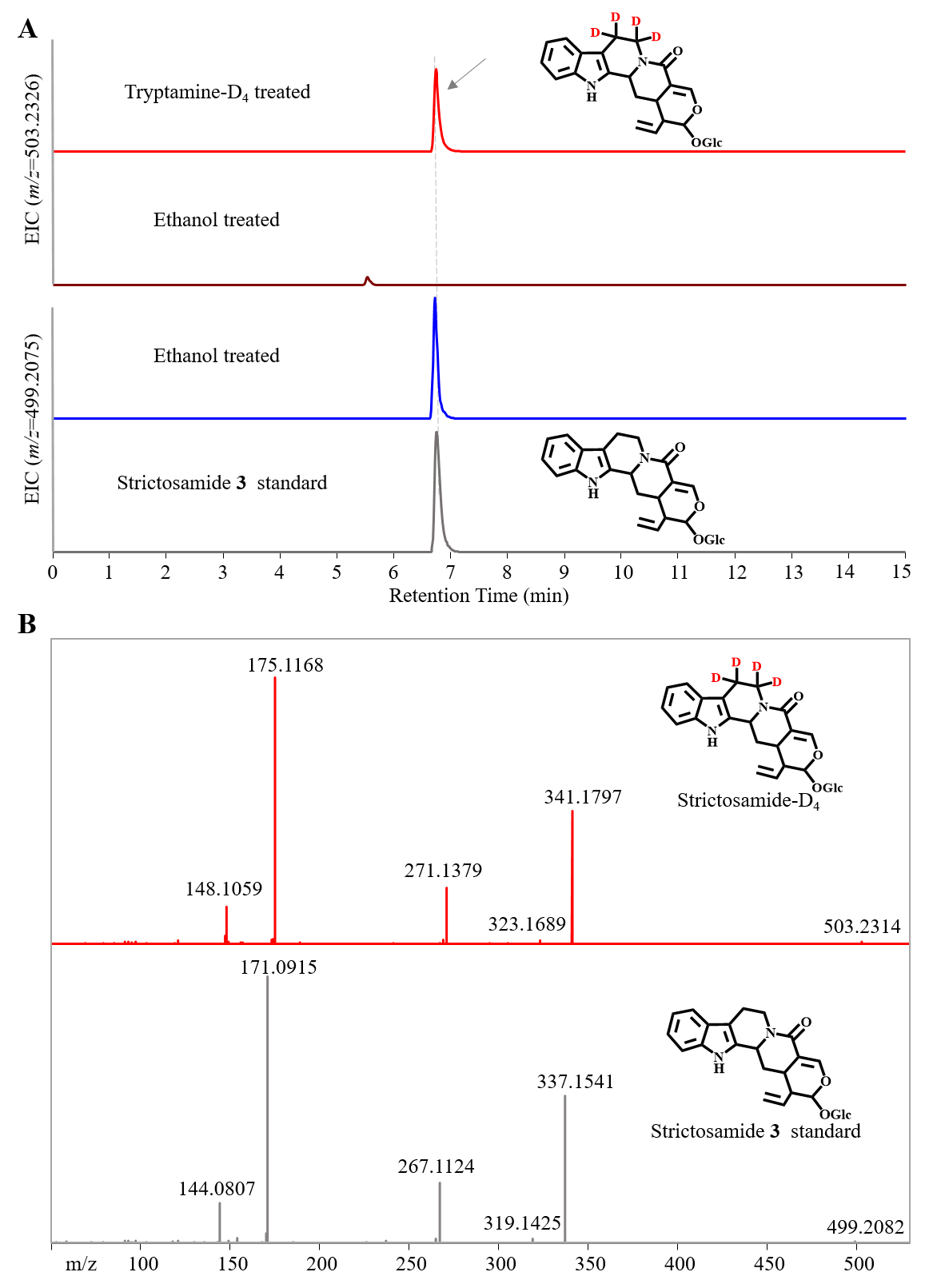

**Fig. S30. LC-MS analysis of strictosamide 3 in the isotope feeding experiment.** (**A**). Extracted ion chromatogram for deuterium-labeled strictosamide-D_4_ (m/z [M+H]^+^ = 503.2326) and strictosamide **3** (m/z [M+H]^+^ = 499.2075) are shown. Tryptamine-D_4_ dissolved in the ethanol was added into the *O. pumila* hairy roots to label the intermediates involved in the camptothecin **9** biosynthesis. Ethanol was solely added into the *O. pumila* hairy roots as the negative control. (**B**). MS/MS spectra of deuterium-labeled strictosamide-D_4_ and standard strictosamide **3**. The main fragments of deuterium-labeled strictosamide-D_4_ were compared with those of standard strictosamide **3**.

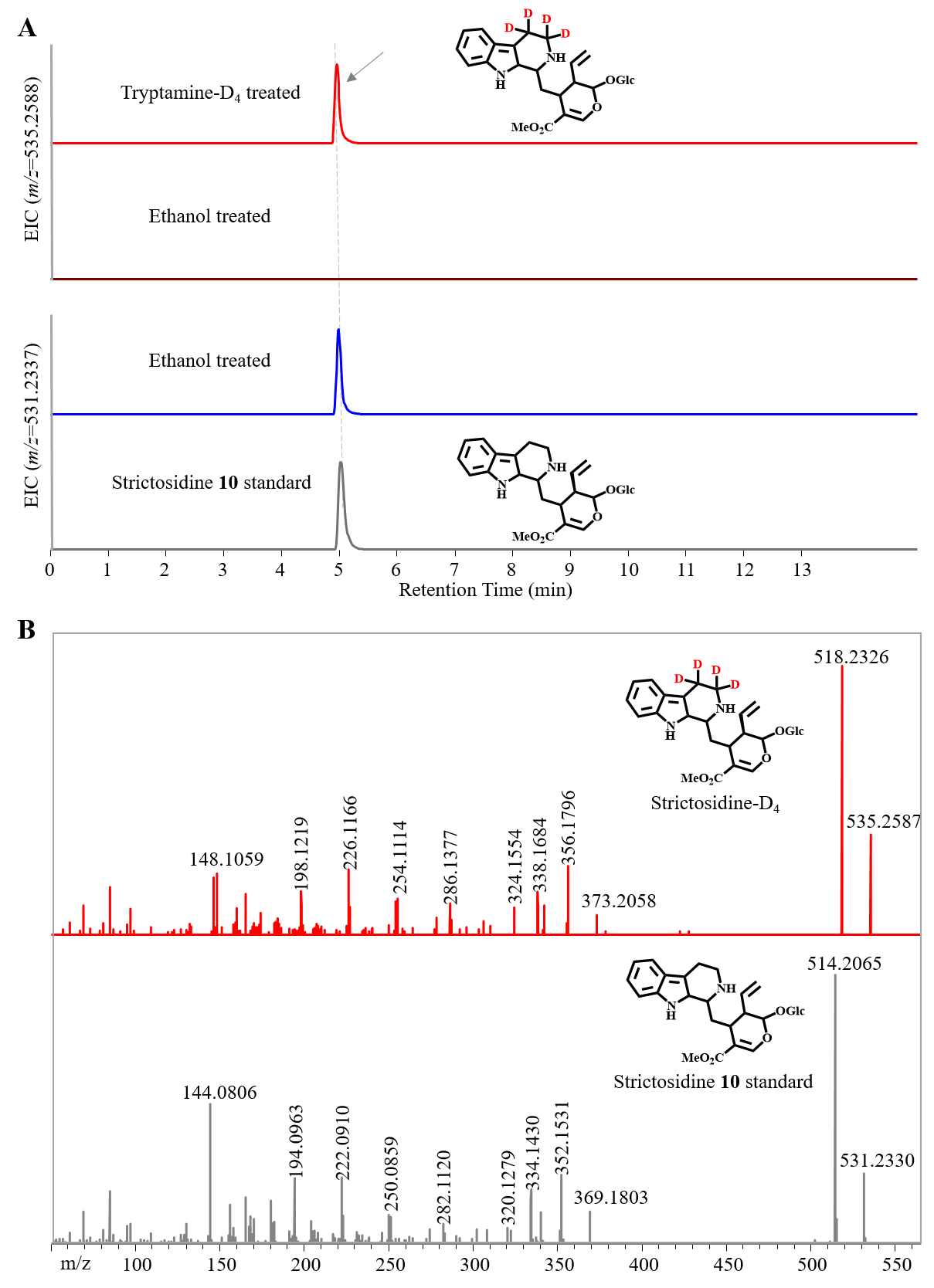

**Fig. S31. LC-MS analysis of strictosidine 10 in the isotope feeding experiment.** (**A**). Extracted ion chromatogram for deuterium-labeled strictosidine-D_4_ (m/z [M+H]^+^ = 535.2588) and strictosidine **10** (m/z [M+H]^+^ = 531.2337) are shown. Tryptamine-D_4_ dissolved in the ethanol was added into the *O. pumila* hairy roots to label the intermediates involved in the camptothecin **9** biosynthesis. Ethanol was solely added into the *O. pumila* hairy roots as the negative control. (**B**). MS/MS spectra of deuterium-labeled strictosidine-D_4_ and standard strictosidine **10**. The main fragments of deuterium-labeled strictosidine-D_4_ were compared with those of standard strictosidine **10**.

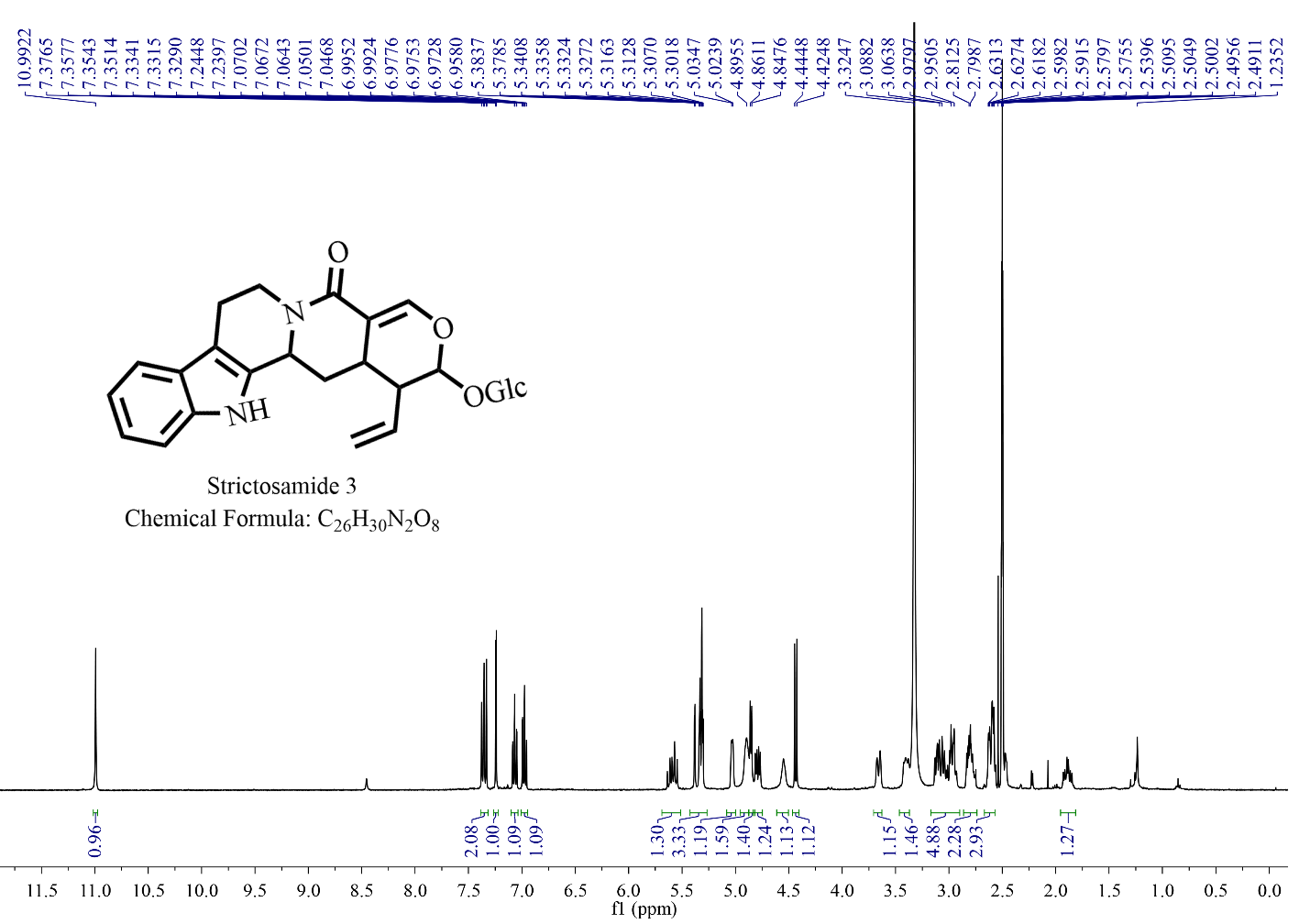

**Fig. S32. ^1^H NMR spectrum of strictosamide 3 produced by OpSTR in vitro.** (400MHz, DMSO-*d*_6_) δ 10.99 (s, 1H), 7.39 – 7.32 (m, 2H), 7.24 (d, *J* = 2.1 Hz, 1H), 7.07 (ddd, *J* = 8.2, 7.0, 1.3 Hz, 1H), 6.98 (ddd, *J* = 7.9, 7.0, 1.1 Hz, 1H), 5.69 – 5.51 (m, 1H), 5.42 – 5.27 (m, 3H), 5.08 – 5.00 (m, 1H), 4.90 (s, 2H), 4.85 (d, *J* = 5.4 Hz, 1H), 4.79 (dd, *J* = 12.6, 5.8 Hz, 1H), 4.55 (s, 1H), 4.43 (d, *J* = 8.0 Hz, 1H), 3.66 (d, *J* = 11.7 Hz, 1H), 3.46 – 3.37 (m, 1H), 3.17 – 2.90 (m, 5H), 2.81 (td, *J* = 8.3, 5.0 Hz, 2H), 2.67 – 2.57 (m, 3H), 1.89 (d, *J* = 5.7 Hz, 1H).

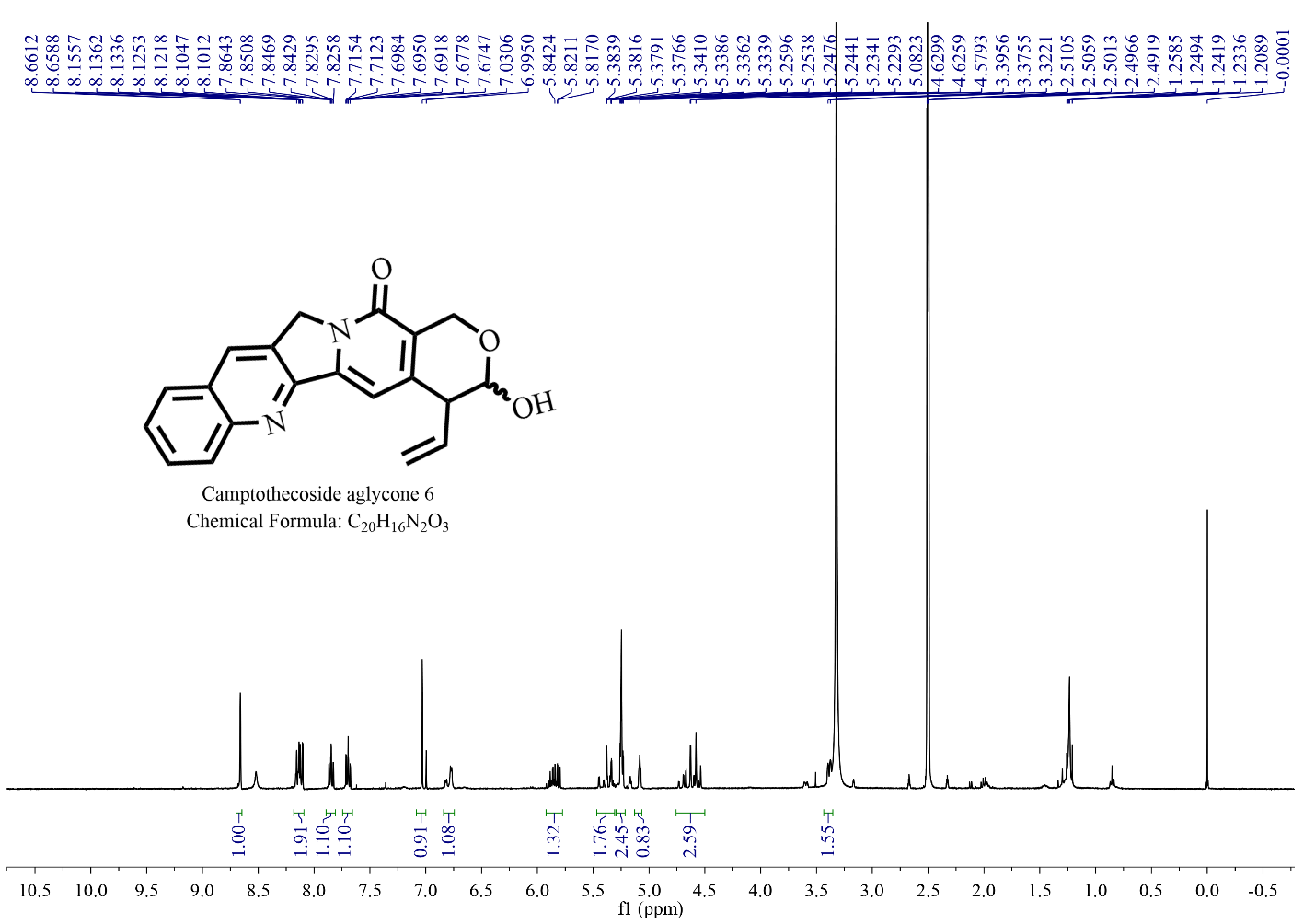

**Fig. S33. ^1^H NMR spectrum of camptothecoside aglycone 6 produced by OpGH1 in vitro.** (400MHz, DMSO-*d*_6_) δ 8.70 – 8.65 (m, 1H), 8.18 – 8.09 (m, 2H), 7.85 (ddd, *J* = 8.4, 6.8, 1.5 Hz, 1H), 7.70 (ddd, *J* = 8.1, 6.8, 1.2 Hz, 1H), 7.03 (s, 1H), 6.77 (d, *J* = 4.5 Hz, 1H), 5.84 (ddd, *J* = 17.1, 10.1, 8.5 Hz, 1H), 5.36 (ddd, *J* = 17.1, 1.9, 0.9 Hz, 2H), 5.29 – 5.21 (m, 2H), 5.08 (t, *J* = 3.3 Hz, 1H), 4.76 – 4.50 (m, 2H), 3.44 – 3.35 (m, 1H).

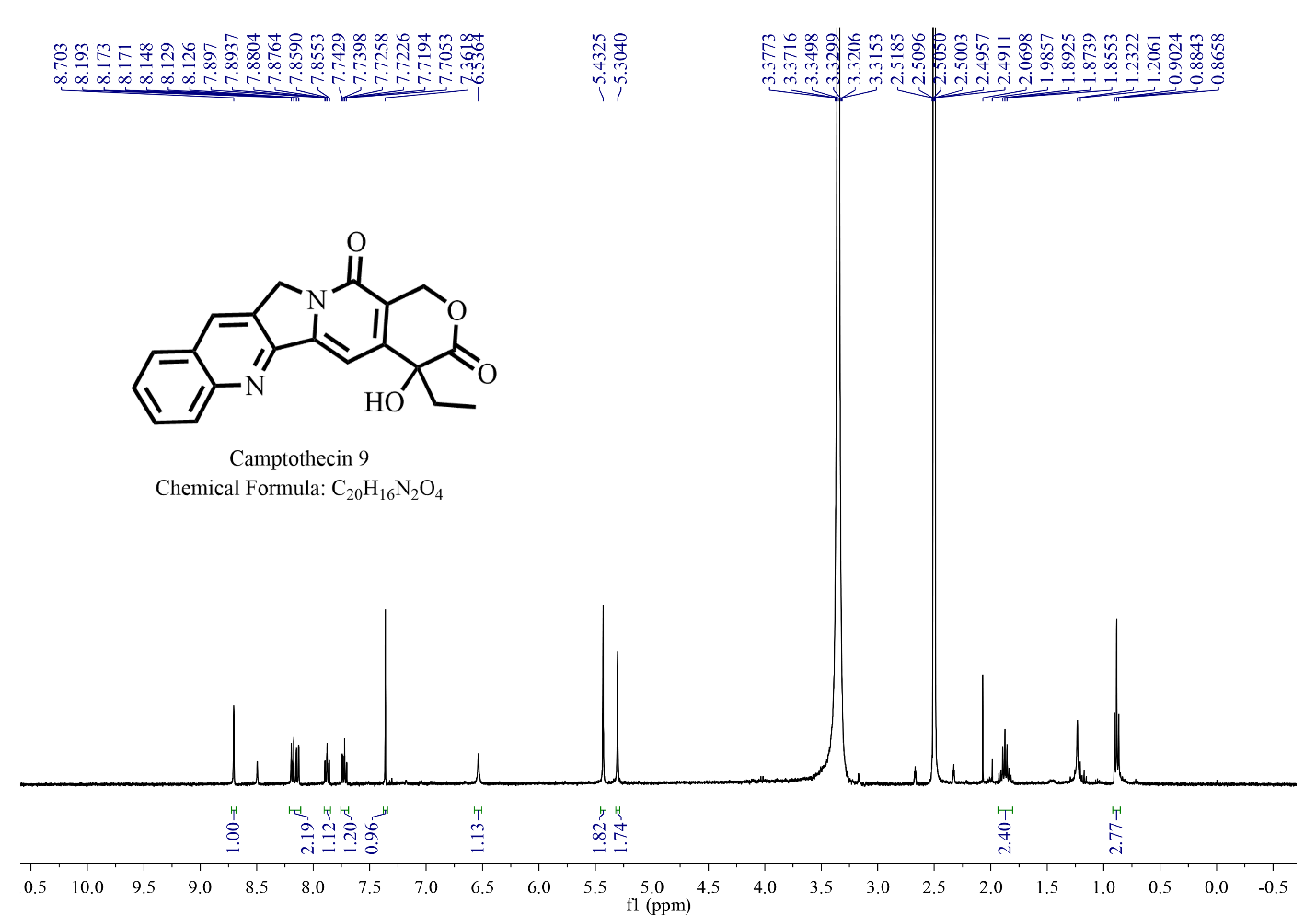

**Fig. S34. ^1^H NMR spectrum of camptothecin 9 produced by OpCS in vitro.** (400 MHz, DMSO-d₆) δ 8.70 (s, 1H), 8.21 – 8.11 (m, 2H), 7.88 (ddd, J = 8.4, 6.8, 1.5 Hz, 1H), 7.72 (ddd, J = 8.2, 6.8, 1.2 Hz, 1H), 7.36 (s, 1H), 6.54 (s, 1H), 5.43 (s, 2H), 5.30 (s, 2H), 1.94 – 1.81 (m, 2H), 0.88 (t, J = 7.3 Hz, 3H).

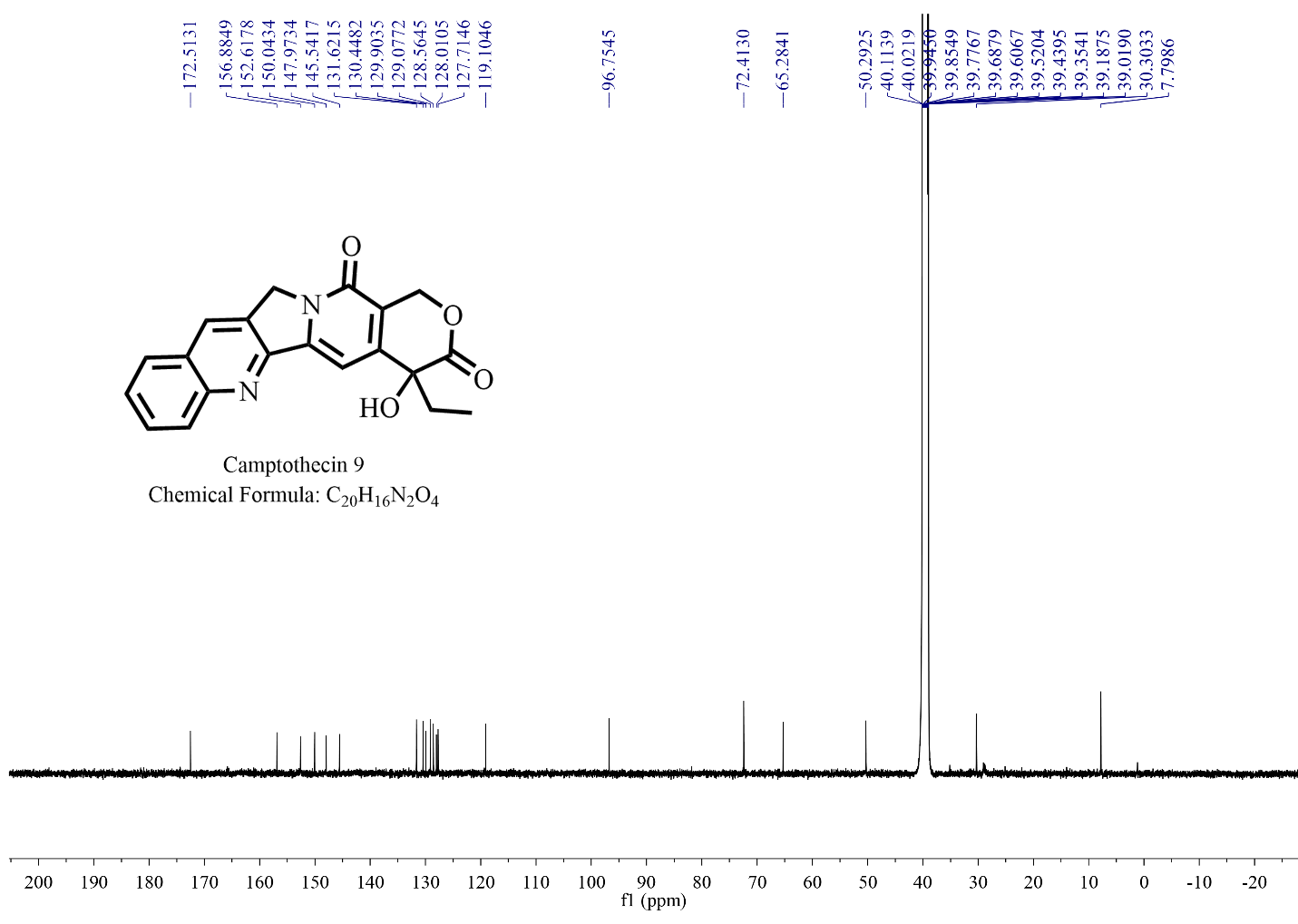

**Fig. S35. ^13^C NMR spectrum of camptothecin 9 produced by OpCS in vitro.** (125MHz, DMSO-d₆) δ 172.51, 156.88, 152.61, 150.04, 147.97, 145.54, 131.62, 130.45, 129.90, 129.08, 128.56, 128.01, 127.71, 119.10, 96.75, 72.41, 65.28, 50.29, 30.31, 7.79.

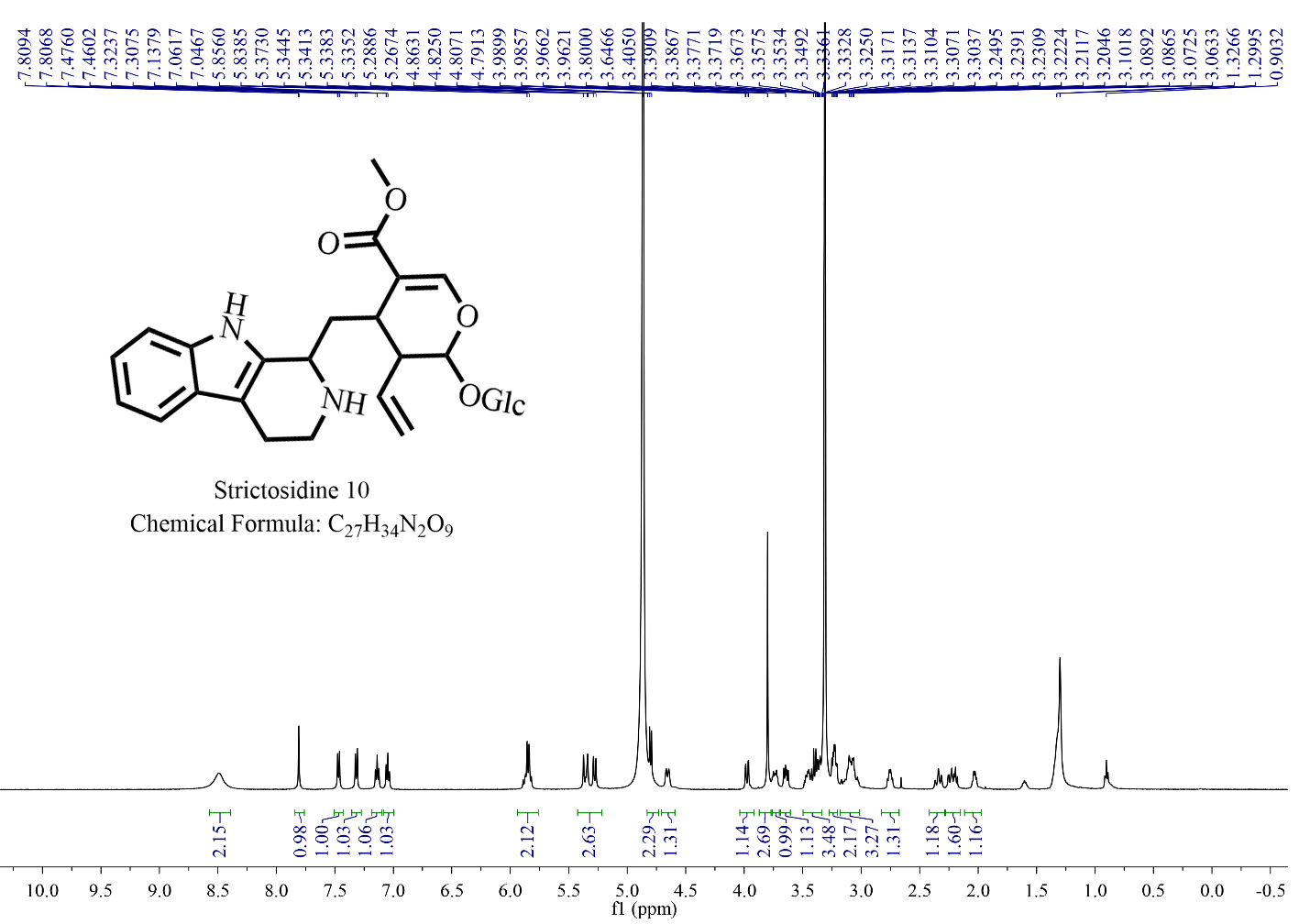

**Fig. S36. ^1^H NMR spectrum of strictosidine 10 produced by OpSTR in vitro.** (400MHz, Methanol-*d*_4_) δ 8.49 (s, 2H), 7.88 – 7.73 (m, 1H), 7.47 (d, J = 7.9 Hz, 1H), 7.32 (d, J = 8.1 Hz, 1H), 7.14 (t, J = 7.6 Hz, 1H), 7.05 (t, J = 7.5 Hz, 1H), 5.92-5.77 (m, 2H), 5.35 (d, J = 17.3 Hz, 1H), 5.28 (d, J = 10.6 Hz, 1H), 4.80 (d, J = 7.9 Hz, 1H), 4.65 (d, J = 11.5 Hz, 1H), 3.98 (dd, J = 11.9, 2.1 Hz, 1H), 3.80 (s, 3H), 3.77 – 3.70 (m, 1H), 3.64 (dd, J = 11.8, 7.0 Hz, 1H), 3.50 – 3.34 (m, 2H), 3.26-3.18 (m, 2H), 3.18 – 3.00 (m, 3H), 2.80 – 2.70 (m, 1H), 2.40 – 2.29 (m, 1H), 2.28 – 2.16 (m, 1H), 2.06 – 1.98 (q, J = 6.2 Hz, 1H).

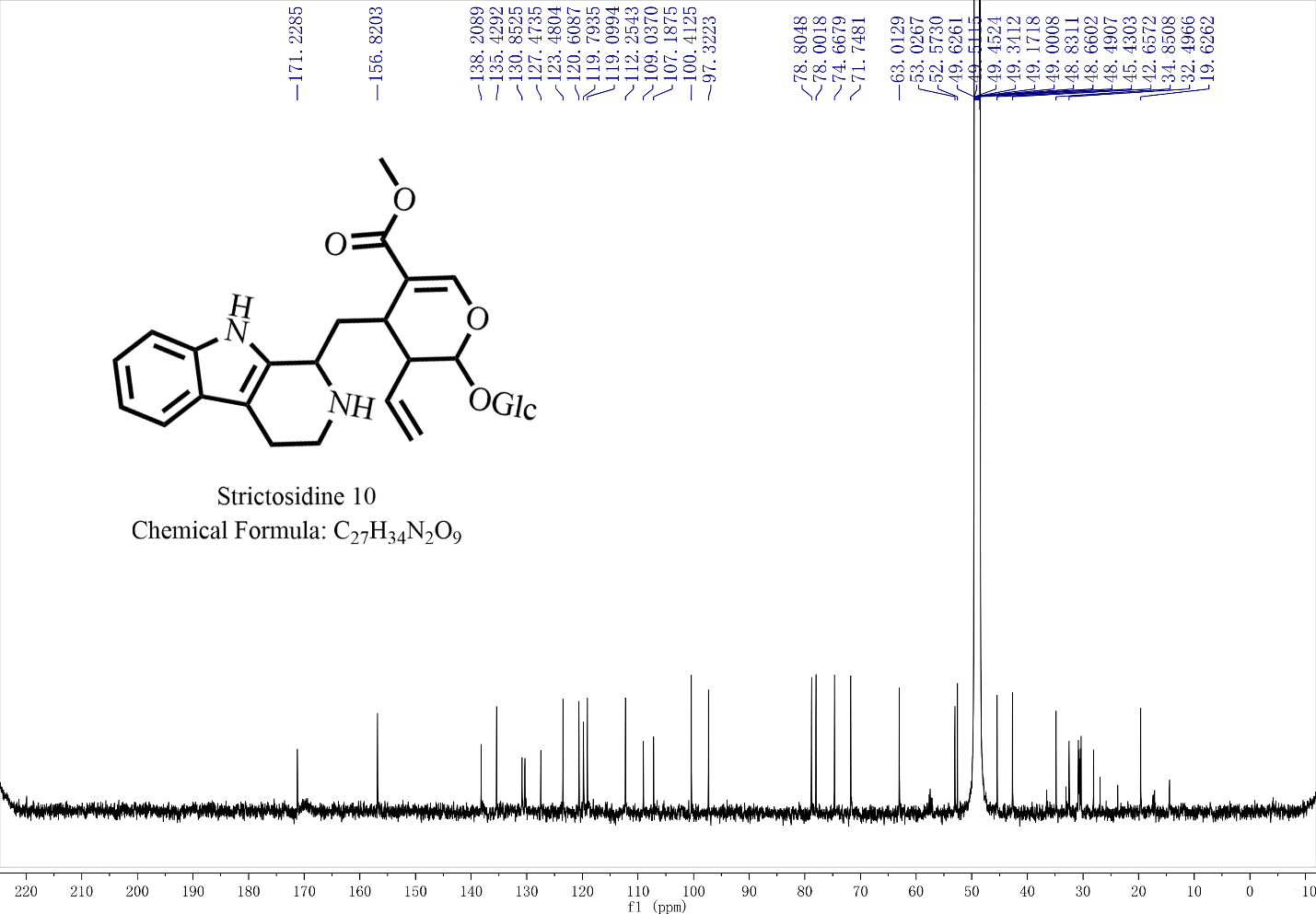

**Fig. S37. ^13^C NMR spectrum of strictosidine 10 produced by OpSTR in vitro.** (125MHz, Methanol-*d*_4_) δ 171.23, 156.82, 138.21, 135.43, 130.85, 127.47, 123.48, 120.61, 119.79, 119.10, 112.25, 109.04, 107.19, 100.41, 97.32, 78.80, 78.00, 74.67, 71.75, 63.01, 53.03, 52.57, 45.43, 42.66, 34.85, 32.50, 19.63.

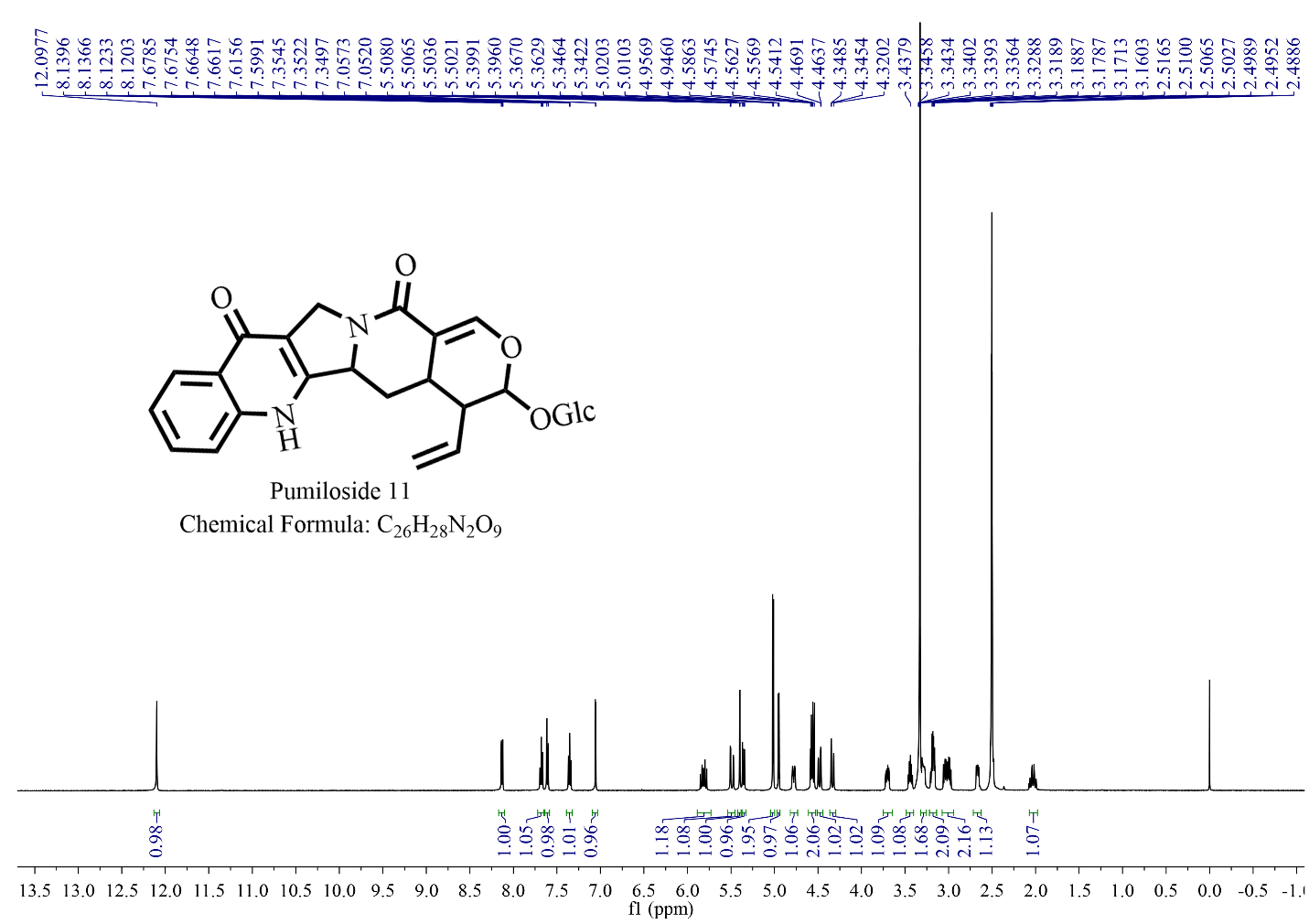

**Fig. S38. ^1^H NMR spectrum of pumiloside 11 produced by flavin mononucleotide.** (500MHz, DMSO-*d*_6_) δ 12.10 (s, 1H), 8.13 (dd, *J* = 8.2, 1.5 Hz, 1H), 7.68 (ddd, *J* = 8.4, 6.9, 1.6 Hz, 1H), 7.61 (d, *J* = 8.3 Hz, 1H), 7.35 (ddd, *J* = 8.1, 6.9, 1.1 Hz, 1H), 7.05 (d, *J* = 2.7 Hz, 1H), 5.81 (dt, *J* = 17.2, 9.8 Hz, 1H), 5.54 – 5.45 (m, 1H), 5.40 (d, *J* = 1.5 Hz, 1H), 5.35 (dd, *J* = 10.3, 2.1 Hz, 1H), 5.02 (d, *J* = 5.0 Hz, 2H), 4.95 (d, *J* = 5.4 Hz, 1H), 4.78 (d, *J* = 11.7 Hz, 1H), 4.61 – 4.53 (m, 2H), 4.48 (dd, *J* = 14.2, 2.7 Hz, 1H), 4.33 (dd, *J* = 14.2, 1.6 Hz, 1H), 3.75 – 3.65 (m, 1H), 3.44 (dt, *J* = 11.8, 5.9 Hz, 1H), 3.33 – 3.25 (m, 2H), 3.18 (tt, *J* = 8.2, 3.8 Hz, 2H), 3.07 – 2.94 (m, 2H), 2.67 (dd, *J* = 9.4, 5.1 Hz, 1H), 2.07 – 1.98 (m, 1H).

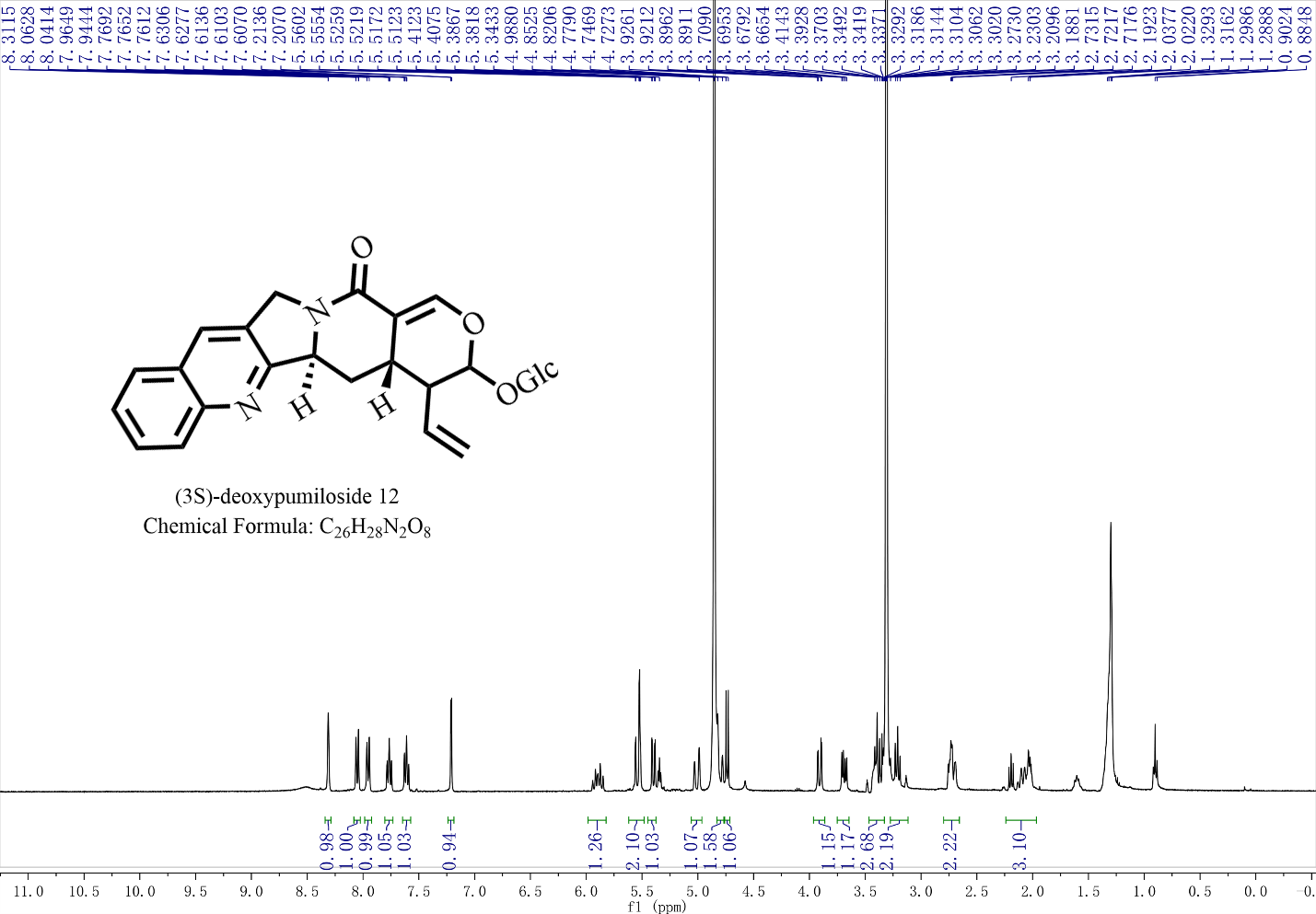

**Fig. S39. ^1^H NMR spectrum of (3*S*)-deoxypumiloside 12.** (400 MHz, Methanol-*d*_4_) δ 8.31 (s, 1H), 8.05 (d, *J* = 8.5 Hz, 1H), 7.95 (d, *J* = 8.2 Hz, 1H), 7.77 (ddd, *J* = 8.5, 6.8, 1.5 Hz, 1H), 7.65 – 7.57 (m, 1H), 7.21 (d, *J* = 2.6 Hz, 1H), 5.90 (dt, *J* = 17.9, 10.0 Hz, 1H), 5.62 – 5.48 (m, 2H), 5.40 (dd, *J* = 10.3, 1.9 Hz, 1H), 5.01 (d, *J* = 16.4 Hz, 1H), 4.80 (d, *J* = 16.6 Hz, 1H), 4.83-4.76 (m, 2H), 3.91 (dd, *J* = 12.0, 2.0 Hz, 1H), 3.69 (dd, *J* = 11.9, 5.5 Hz, 1H), 3.47 – 3.33 (m, 3H), 3.27 – 3.12 (m, 2H), 2.73 (tt, *J* = 11.8, 5.7 Hz, 2H), 2.24 – 1.97 (m, 1H).

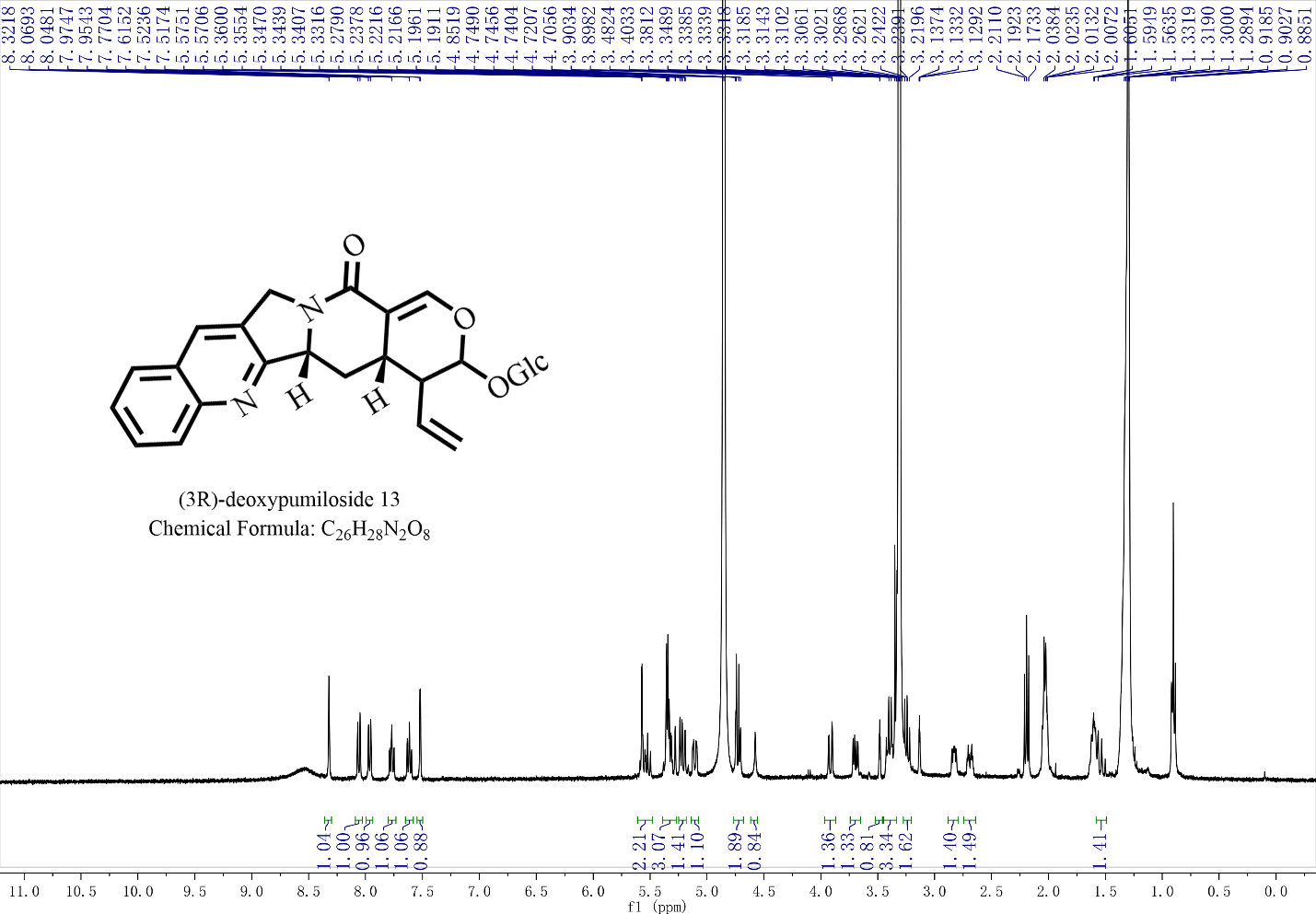

**Fig. S40. ^1^H NMR spectrum of (3*R*)-deoxypumiloside 13.** (400 MHz, Methanol-*d*_4_) δ 8.32 (s, 1H), 8.06 (d, *J* = 8.5 Hz, 1H), 7.96 (d, *J* = 8.2 Hz, 1H), 7.77 (ddd, *J* = 8.4, 6.8, 1.4 Hz, 1H), 7.65 – 7.58 (m, 1H), 7.52 (d, *J* = 2.5 Hz, 1H), 5.61 – 5.48 (m, 2H), 5.39 – 5.27 (m, 1H), 5.25 – 5.18 (m, 1H), 5.11 (dd, *J* = 11.2, 3.1 Hz, 1H), 4.76 – 4.68 (m, 2H), 3.92 (dd, *J* = 12.0, 2.0 Hz, 1H), 3.56 – 3.47 (m, 1H), 3.44 – 3.30 (m, 4H), 3.30 – 3.21 (m, 2H), 2.83 (dd, *J* = 9.4, 5.3 Hz, 1H), 2.74 – 2.64 (m, 1H), 1.61-1.50 (m, 1H).

**Table S1. The known genes involved in strictosamide 3 biosynthesis**

| **Gene name** | **Gene ID** |
| --- | --- |
| GES  G8H | Op10g01262  Op05g00545 |
| 8HGO | Op04g01394 |
| IS | Op03g01921 |
| IO | Op01g00532 |
| 7-DLGT | Op09g01577 |
| 7-DLH | Op10g00805 |
| LAMT | Op10g00724 |
| TDC | Op07g00629 |
| SLS | Op10g00801 |
| STR | Op10g01176 |

**Table S2. Functional characterization of OpCS with different substrates**

| **Protein name** | **Substrate** | **Conversion rate** |
| --- | --- | --- |
| OpCS | Deoxycamptothecin **8** | 10% |
| OpCS | Camptothecoside aglycone **6** | 0 |
| OpCS | Deoxypumiloside aglycone **5** | 0 |
| OpCS | Strictosamide aglycone **4** | 0 |
| OpCS | Strictosamide **3** | 0 |
| OpCS | (3*S*)-deoxypumiloside **12** | 0 |
| OpCS | (3*R*)-deoxypumiloside **13** | 0 |

**Table S3. Canonical marker genes of *A. thaliana* and *O. pumila***

| **Species** | **Canonical markers**  **(gene numbering)** | **Cell types** | **Homologous genes of *O. pumila*** |
| --- | --- | --- | --- |
| *A. thaliana* | AT5G53370 | Cortex | Opuchr06-g0003570-1.1 |
| *A. thaliana* | AT5G53370 | Cortex | Opuchr06-g0003570-1.1 |
| *A. thaliana* | AT5G53370 | Cortex | Opuchr06-g0003570-1.1 |
| *A. thaliana* | AT1G61590 | Endodermis | Opuchr08-g0062080-1.1 |
| *A. thaliana* | AT1G61590 | Endodermis | Opuchr08-g0062080-1.1 |
| *A. thaliana* | AT1G61590 | Endodermis | Opuchr08-g0062080-1.1 |
| *A. thaliana* | AT1G32450 | Pericycle | Opuchr02-g0008300-1.1 |
| *A. thaliana* | AT1G32450 | Pericycle | Opuchr02-g0008300-1.1 |
| *A. thaliana* | AT1G79430 | Phloem | Opuchr05-g0068550-1.1 |
| *A. thaliana* | AT1G79430 | Phloem | Opuchr05-g0068550-1.1 |
| *A. thaliana* | AT1G79430 | phloem | Opuchr05-g0068550-1.1 |
| *A. thaliana* | AT1G79430 | phloem | Opuchr05-g0068550-1.1 |
| *A. thaliana* | AT1G79430 | Phloem | Opuchr05-g0068550-1.1 |
| *A. thaliana* | AT1G79430 | Phloem | Opuchr05-g0068550-1.1 |
| *A. thaliana* | AT1G68810 | Xylem | Opuchr11-g0077910-1.1 |
| *A. thaliana* | AT1G68810 | Xylem | Opuchr11-g0077910-1.1 |
| *A. thaliana* | AT1G33280 | Root cap | Opuchr10-g0063950-1.1 |
| *A. thaliana* | AT3G20840 | Quiescent Center | Opuchr07-g0011840-1.1 |
| *A. thaliana* | AT3G20840 | Quiescent Center | Opuchr07-g0011840-1.1 |
| *A. thaliana* | AT3G20840 | Quiescent Center | Opuchr07-g0011840-1.1 |
| *A. thaliana* | AT3G20840 | Quiescent Center | Opuchr07-g0011840-1.1 |

**Table S4. The top 50 genes highly expressed in the cortex cells**

| Gene number | Average_logFC | Function analysis |
| --- | --- | --- |
| 1 | 2.7315788 | Hydrolase |
| 2 | 2.6378667 | Oxidase |
| 3 | 2.4783165 | Major allergen Pru ar 1-like |
| 4 | 2.4594207 | Major allergen Pru ar 1-like |
| 5 | 2.4587595 | CYP450 |
| 6 | 2.3930073 | WAT1-related protein |
| 7 | 2.3729146 | Glutathione S-transferase |
| 8 | 2.2427058 | Ethylene-responsive transcription factor |
| 9 | 2.2070534 | Glutathione S-transferase |
| 10 | 2.1959696 | WAT1-related protein |
| 11 | 2.186103 | 2-succinylbenzoate--CoA ligase |
| 12 | 2.1559355 | Peroxidase 47-like |
| 13 | 2.1009605 | Mediator of RNA polymerase II |
| 14 | 2.0778718 | Remorin family protein |
| 15 | 2.0591228 | Heavy metal-associated isoprenylated protein |
| 16 | 2.0221276 | Caffeic acid 3-O-methyltransferase-like |
| 17 | 2.0219374 | 12-oxophytodienoate oxidase 2-like |
| 18 | 1.9928836 | Mitogen-activated protein kinase |
| 19 | 1.9565735 | Protein EXORDIUM-like 2 |
| 20 | 1.9560789 | Peroxidase 42 |
| 21 | 1.9491706 | Heat shock cognate protein |
| 22 | 1.9359796 | Iridoid synthase CYC2-like |
| 23 | 1.8933716 | AP2/ERF transcription factor |
| 24 | 1.8520389 | UDP-glycosyltransferase |
| 25 | 1.8517765 | Plant invertase |
| 26 | 1.838793 | AAA-ATPase ASD, mitochondrial-like |
| 27 | 1.8320626 | Protein-serine,threonine phosphatase |
| 28 | 1.8015625 | Phosphatidyltransferase |
| 29 | 1.8001959 | Hydroquinone glucosyltransferase-like |
| 30 | 1.7958498 | Unknown protein |
| 31 | 1.7935997 | 1, 4-dihydroxy-2-naphthoyl-CoA synthase |
| 32 | 1.7803947 | AAA-ATPase |
| 33 | 1.7669822 | Adenine phosphoribosyltransferase |
| 34 | 1.7569678 | Phospholipase A1-Igamma1 |
| 35 | 1.752989 | Late embryogenesis abundant protein |
| 36 | 1.7440339 | Proteinase inhibitor I3 |
| 37 | 1.7294513 | Calcium-binding |
| 38 | 1.7290128 | Oxidoreductase (OpCAR) |
| 39 | 1.718333 | Calcium-binding protein CP1-like |
| 40 | 1.7142087 | Filament-like plant protein 7 |
| 41 | 1.696073 | Peroxidase |
| 42 | 1.6928388 | CYP450 |
| 43 | 1.6641101 | Disease resistance response protein |
| 44 | 1.6603848 | Late embryogenesis abundant protein |
| 45 | 1.6375517 | Sodium/calcium exchanger NCL2-like |
| 46 | 1.6343249 | Alpha/beta-hydrolases superfamily |
| 47 | 1.6126099 | Peroxidase 10 |
| 48 | 1.5859368 | Blue copper protein-like |
| 49 | 47.751007 | ABC transporter |
| 50 | 1.5610431 | Dehydrin family protein |
